# Tissue-specific activities of the Fat1 cadherin cooperate to control neuromuscular morphogenesis

**DOI:** 10.1101/207308

**Authors:** Françoise Helmbacher

## Abstract

Muscle morphogenesis is tightly coupled with that of motor neurons (MNs). Both MNs and muscle progenitors simultaneously explore the surrounding tissues while exchanging reciprocal signals to tune their behaviors. We previously identified the Fat1 cadherin as a regulator of muscle morphogenesis, and showed that it is required in the myogenic lineage to control the polarity of progenitor migration. To expand our knowledge on how Fat1 exerts its tissue-morphogenesis regulator activity, we dissected its functions by tissu-specific genetic ablation. An emblematic example of muscle under such morphogenetic control is the cutaneous maximus (CM) muscle, a flat subcutaneous muscle in which progenitor migration is physically separated from the process of myogenic differentiation, but tightly associated with elongating axons of its partner motor neurons. Here, we show that constitutive *Fat1* disruption interferes with expansion and differentiation of the CM muscle, with its motor innervation and with specification of its associated MN pool. *Fat1* is expressed in muscle progenitors, in associated mesenchymal cells, and in MN subsets including the CM-innervating pool. We identify mesenchyme-derived connective tissue as a cell type in which *Fat1* activity is required for the non-cell-autonomous control of CM muscle progenitor spreading, myogenic differentiation, motor innervation, and for motor pool specification. In parallel, *Fat1* is required in MNs to promote their axonal growth and specification, indirectly influencing muscle progenitor progression. These results illustrate how *Fat1* coordinates the coupling of muscular and neuronal morphogenesis by playing distinct but complementary actions in several cell types.

**Author summary:** Fat cadherins are evolutionarily conserved cell adhesion molecules playing key roles in modulating tissue morphogenesis, through the control of collective cell behavior and polarity. We previously identified the mouse *Fat1* gene as a regulator of muscle morphogenesis. The present study explores how *Fat1* influences neuromuscular morphogenesis in the context of development of a flat subcutaneous muscle, the cutaneous maximus muscle (CM), formed by migratory progenitors emerging from forelimb levels somites, and innervated by a pool of brachial spinal motor neurons (MNs). CM development involves the rostrocaudal planar migration of muscle progenitors and subsequent elongation of muscle fibers to form a fan-shaped muscle. We previously reported that *Fat1* was required in muscle progenitors to modulate their migration polarity. Here, these results were expanded by exploring the contribution of *Fat1* activities in two other cell types, mesenchymal cells and MNs. We show that *Fat1* disruption in connective tissue robustly alters CM muscle morphogenesis, affecting not only progenitor migration and myofiber expansion, but also subsequently impairing axon growth and specification of cognate MNs. In parallel, *Fat1* acts in MNs to modulate axonal growth and neuronal specification, modestly influencing muscle morphology. Together, these results show that *Fat1* coordinates the coupling between muscle and neuronal development by playing complementary functions in mesenchyme, muscles and MNs. These findings could guide research on muscle pathologies associated with *FAT1* alterations in humans.

## Introduction

Neuromuscular morphogenesis involves complex tissue interactions simultaneously governing the generation of skeletal muscles, and the production of the somatic motor neurons (MNs) that innervate them. Both processes independently rely on the execution of a generic regulatory program sequentially leading to cell fate determination, differentiation, and functional diversification [1–3]. These regulatory events are coupled with dynamic morphogenetic events leading to the definition of multiple muscle shapes, and the simultaneous topographical wiring of motor axonal projections. During muscle morphogenesis, myogenic progenitors migrate collectively from their origin in either somites or cephalic mesoderm, to their final position in the limb, trunk or face [4]. Trunk and limb connective tissues, which derives from lateral plate mesoderm, provide instructive signals for incoming somite-derived myogenic cells [5]. Reciprocal signals exchanged by muscle progenitors and the mesenchymal environment pattern muscle shapes by allowing the definition of fiber orientation, muscle attachment sites, and tendon development [5,6]. Muscle progenitors subsequently engage in a complex regulatory process through which they give rise to differentiating cells called myocytes, in charge of producing the contractile apparatus [2,3,7]. Myocytes then fuse with each other to form multinucleated muscle fibers. The process of muscle growth is determined by a tightly regulated balance between progenitor expansion and production of myocytes and differentiating muscle fibers [2,3]. In parallel with muscle morphogenesis, motor neurons emit axons, which grow in peripheral tissues, selecting a trajectory by responding to multiple guidance cues, allowing them to find their target muscles, within which they ultimately establish a selective pattern of intramuscular arborization [8]. During these processes, axons of motor neurons and migratory myogenic progenitors follow converging trajectories, along which they simultaneously probe the environment and respond to instructive cues, as evidenced by classical embryological studies [5,9–11]. Multiple signals emitted by peripheral tissues to instruct MNs specification and axonal pathfinding have been identified [12]. Likewise, some recent discoveries have started shedding light on how non-myogenic connective tissues exert their influence on muscle patterning [13]. In spite of such advances, what controls the coordinated behavior of the two cell types to orchestrate neuromuscular morphogenesis has not been studied.

In the present study, we have examined the possibility that the Fat1 cadherin, a planar cell polarity (PCP) molecule involved in tissue morphogenesis, could contribute to coordinate muscular and neuronal morphogenesis. We recently identified *Fat1* as a new player in muscle morphogenesis, which influences the shape of subsets of face and shoulder muscles, in part by polarizing the direction of collectively migrating myoblasts [14]. Fat1 belongs to the family of Fat-like atypical cadherins [15]. Together with their partner cadherins Dachsous, Fat-like cadherins are involved in regulating coordinated cell behaviors such as planar cell polarity in epithelia [15–17], collective/polarized cell migration [18–20], and oriented cell divisions [21,22]. Through these actions, Fat/Dachsous signaling modulates cell orientation, junctional tension [23,24], and microtubule dynamics [25], thereby influencing the mechanical coupling between cell behavior and tissue shapes [17]. Aside from their canonical role in regulating the planar cell polarity pathway [16], Fat-like cadherins also control tissue growth via the Hippo pathway [26,27], and were recently found to contribute to mitochondria function and metabolic state by interacting with the electron transport chain [28,29]. In vertebrates, the most studied *Fat* homologue, *Fat4,* plays multiple functions in development, to coordinate kidney [22,30–32], skeletal [21], heart [33], or neural morphogenesis [19,34,35]. The other family member, *Fat1,* is known for playing complementary functions during kidney [36,37], muscle [14], and neural [38] morphogenesis.

Here, to explore the mechanisms underlying the coupling of neural and muscular morphogenesis and to assess how *Fat1* contributes to this process, we focused on a large flat subcutaneous muscle, the Cutaneous Maximus (CM), which expands under the skin by planar polarized myoblast migration. This muscle is linked to its cognate spinal MN pool through the selective production by the CM muscle of Glial-derived-Neurotrophic-Factor (GDNF), a secreted growth factor required to control specification of the corresponding MNs [39]. Unlike limb-innervating MNs, which are born with an intrinsic molecular program specifying their anatomical characteristics [9,11], CM-innervating MNs are incompletely specified at the time they first send their axons, and are dependent on extrinsic signals from peripheral tissues [39–41]. GDNF, produced first by the plexus mesenchyme, and subsequently by the CM and LD muscles, is perceived by axons of a competent population of motor neurons when they reach the plexus and as they continue growing along the expanding muscle [39,42]. GDNF acts through the Ret tyrosine kinase receptor in a complex with a GPI-anchored co-receptor Gfra1 [43–45], by inducing expression of the transcription factor Etv4 (also known as Pea3) in the MN pools innervating the CM and LD muscles [39], in synergy with another limb-derived factor, Hepatocyte Growth Factor (HGF) [46,47]. Etv4 in turn influences MN cell body positioning, dendrite patterning, intramuscular axonal arborization, and monosynaptic reflex circuit formation [40,41]. Whereas MN cell body positioning is thought to involve the regulation of the Cadherin code by Etv4 [41,48], patterning of sensory-motor connections is accounted for by the *Etv4*-regulated Sema3E, acting as repellent cue for sensory neurons expressing its receptor PlexinD1 [49,50]. GDNF can also directly influence axon pathfinding, as demonstrated in the context of dorsal motor axon guidance [51–54], or of midline-crossing by commissural axons [55], and is subsequently required for survival of subsets of MNs [56,57].

We found that inactivation of the *Fat1* gene has a profound impact on the assembly of the CM neuromuscular circuit, affecting not only the rate of subcutaneous expansion of the CM muscle by progenitor migration, and the subsequent rate of differentiation, but also the acquisition of identity and projection patterns of their cognate MNs. Intriguingly, in addition to its function in myogenic cells [14], *Fat1* is also expressed in muscle-associated mesenchymal cells, and in the MN subset corresponding to the *Fat1-*dependent CM muscle. Through a series of genetic experiments in mice, we have selectively ablated *Fat1* functions in the distinct tissue types in which it is expressed along this neuromuscular circuit, and assessed the impact on muscular and neuronal morphogenesis. We uncovered two novel *Fat1* functions in the mesenchymal lineage and in motor neurons, which synergize to coordinate the development of the CM neuromuscular circuit. *Fat1* ablation in the mesenchymal lineage causes severe non-cell-autonomous alterations of CM morphogenesis, disrupting expansion of the GDNF-expressing CM progenitors and the subsequent processes of muscle fiber elongation, CM motor innervation, and the acquisition of MN pool identity. This identifies the mesenchymal lineage as a source of *Fat1*-dependent muscle- and MN-patterning cues. The neural consequences of mesenchyme-specific *Fat1* ablation partially mimic the effects of *Gdnf* or *Etv4* mutants, and can be aggravated by further reducing *Gdnf* levels genetically. In parallel, we find that MN-*Fat1* is required cell-autonomously for motor axon growth and MN specification. Unexpectedly MN-specific *Fat1* ablation also influences myogenic progenitor spreading in a non-cell-autonomous manner, demonstrating a reverse influence of MNs on muscle morphogenesis. Collectively, these data show that *Fat1* exerts complementary functions in several tissue-types along the circuit, each of which contributes to neuromuscular morphogenetic coupling through distinct mechanisms, coordinating the adaptation of MN phenotype to muscle shape.

## Results

### Loss of *Fat1* alters posterior expansion and differentiation of the Cutaneous Maximus muscle

In this study, we have used two phenotypically equivalent constitutive knockout alleles of *Fat1* (summarized in S1 Table): The first allele (*Fat1^-^* allele, also known as *Fat1^ΔTM^)* derives from the conditional *Fat1^Flox^* allele by CRE-mediated recombination of the floxed exons encoding the transmembrane domain, thus abrogating the ability of the Fat1 protein to transduce signals [14]. The second allele (*Fat1^LacZ^* allele) is a genetrap allele of *Fat1,* in which the inserted transgene results in expression of a Fat1-β-galactosidase chimeric protein, in the endogenous domain of *Fat1* expression [14]. Both alleles cause comparable phenotypes [14].

We focused on one of the muscles affected by constitutive loss of *Fat1* functions, the Cutaneous Maximus (CM) muscle (Fig 1A, B), a flat muscle emerging from the forelimb plexus (also called brachial plexus). Completing our previous analysis [14], we first followed the establishment of the CM muscle and its evolution during development by using a transgenic line (the *Mlc3f-nLacZ-2E* line, later referred to as *MLC3F-2E*) expressing an *nLacZ* reporter in differentiating muscle cells, thus behaving as a reporter of sarcomeric gene expression [58,59]. This line was combined to the constitutive *Fat1^-^* allele, and wild type and mutant embryos carrying the transgene were stained with X-gal. We previously reported that this approach reveals in *Fat1^-/-^* embryos: 1) myocytes dispersed in the forelimb region; 2) a supernumerary muscle in ectopic position in the upper part of the forelimb (see Fig 6 in ref [14], S1A Fig, S1B Fig). Both phenotypes can be quantified (S1B Fig), and can also be observed on histological sections, using additional markers of muscle development, such as Pax7 to label myogenic progenitors, and the sarcomeric protein Myh1 to visualize muscle fibers (S1C Fig, S1D Fig). The evolution of the CM muscle follows a very specific growth pattern. Muscle fibers can be viewed as “chains” of *MLC3F-2E* positive nuclei. In the CM, these fibers appear to originate from the brachial plexus, just posterior to the forelimb bud, and to extend posteriorly from this point. Individual fibers spread in a radial manner under the skin to form a fan shaped structure, ranging from dorsally directed fibers to ventrally directed fibers, the median direction being approximately horizontal. As development proceeds, the length of such fibers (yellow arrows, Fig 1A, B) increases posteriorly, and the overall area covered by β-Galactosidase-positive fibers (red dotted area, Fig 1A, B) expands. The rate of CM expansion can be measured by following the area containing differentiated fibers, plotted relative to the area of body wall (BW) muscles (yellow dotted area on upper pictures in Fig 1A, B), used as a proxy for the embryo stage, as these muscles are not affected by the mutation. In wild-type *MLC3F-2E^+^* embryos, the CM area follows a positive linear evolution, strongly correlated with expansion of the body wall muscle area (Fig 1C, left plot). In *Fat1^-/-^; MLC3F-2E^+^* embryos, CM expansion appears severely delayed, although not abolished, with a growth rate reduced by more than twofold compared to control embryos (Fig 1C, left plot). As a result, at comparable stages, the differentiated CM area is systematically smaller in absence of functional *Fat1*. Given the highly dynamic nature of CM expansion over just one day, in order to pool data from all embryos examined, the ratio between the CM and BW areas was calculated, and normalized to the median ratio of control littermate embryos, corresponding to 100% (Fig 1C, right plot). Overall, loss of *Fat1* functions causes the area of differentiated CM to be reduced to a median value of ~32% compared to control embryos. Interestingly, observation of older embryos (Fig 1B) reveals that fiber length and *LacZ*-positive nuclei density appear more drastically reduced in the ventral part of the CM than in the dorsal part. We previously documented that at later stages, in this ventral area, occurrence of misoriented fibers crossing each other can frequently be observed [14].

**Fig 1:**
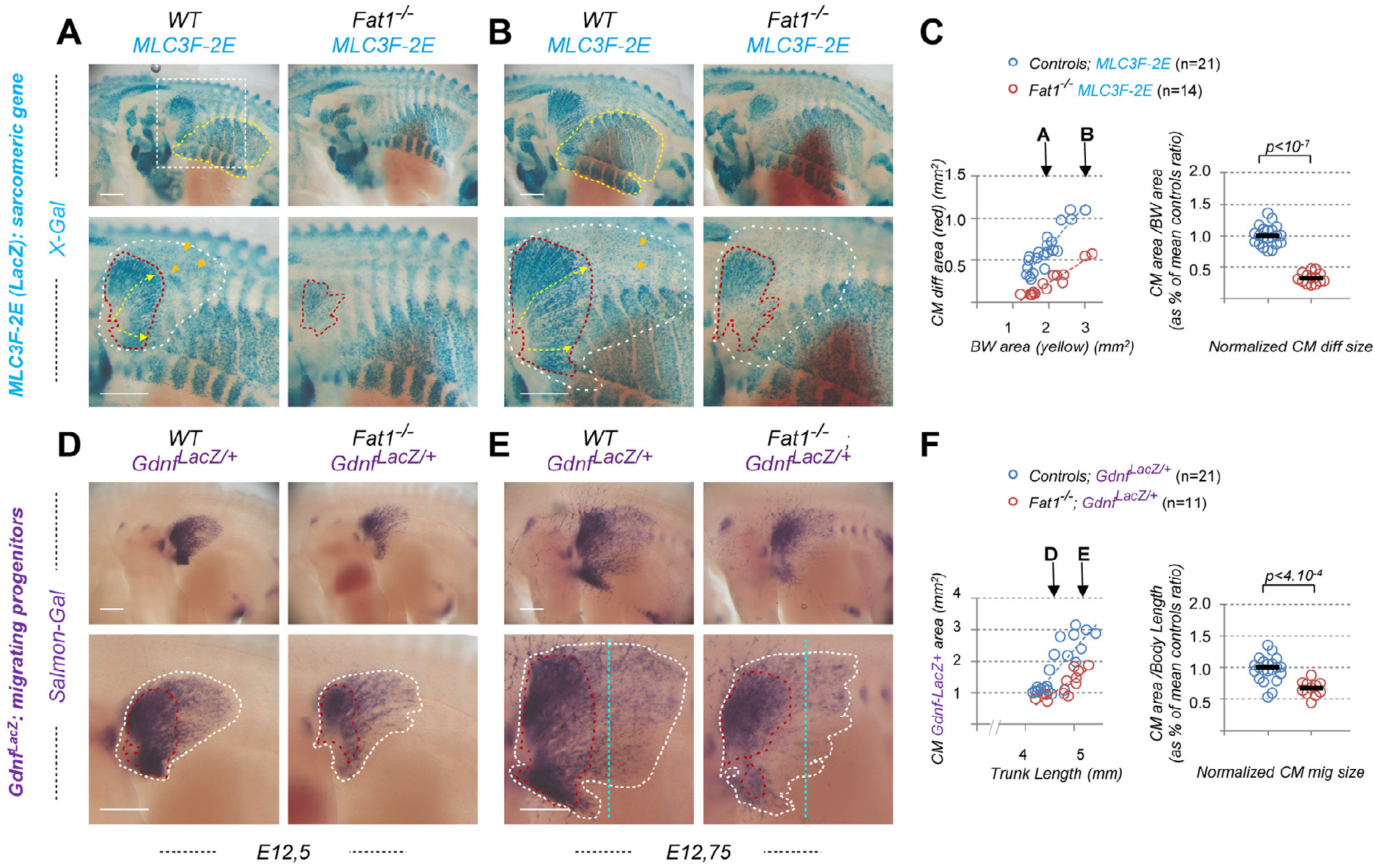
*Fat1* knockout alters expansion of the subcutaneous muscle Cutaneous Maximus. Whole-mount β-galactosidase staining was performed using X-gal as substrate on embryos carrying the *MLC3F-2E* transgene **(A, B)**, or using Salmon-Gal as substrate on embryos carrying the *Gdnf^LacZ/+^* allele **(D, E)**. In each case, two successive stages are shown, E12.5 **(A,D)** and E12.75 **(B, E)**, respectively with *Fatl^+/+^;* (left) and *Fatl^-/-^* (right) embryos, with the lower panels showing a higher magnification of the flank in which the Cutaneous Maximus muscle (CM) spreads. On upper panels in **(A, B)**, the yellow dotted line highlights the body wall muscles, which area is being measured. The white square highlights the area shown on lower panels. Lower panels: the red dotted line highlights the area covered by differentiating *MLC3F-2E^+^* muscle fibers constituting the CM muscle in **(A, B)**, also matching an area of higher *Gdnf^LacZ^* intensity in **(D, E)**; the white dotted lines highlight the area corresponding to the full shape of the *Gdnf^LacZ+^* area in **(D, E)**, in which a low density of blue *(MLC3F-2E^+^)* nuclei can also be observed in **(A,B)**. **(C)** Quantifications of the relative expansion of the *MLC3F-2E^+^* CM differentiated area. Left plot: for each embryo side, the area of differentiated CM is plotted relative to the body wall area. Arrows represent the stages shown in **(A)** and **(B)**, respectively. Right plot: for each embryo, the CM area/Body wall area was normalized to the median ratio of control embryos. Blue dots: *Fat1^+/+^; MLC3F-2E* (n=21); Red dots: *Fat1^-/-^; MLC3F-2E* (n=14). Underlying data are provided in S1 Data. **(F)** Quantifications of the relative expansion of the *Gdnf^LacZ+^* area. Left plot: for each embryo side, the *Gdnf^LacZ+^* area is plotted relative to the length of the trunk (measured between two fixed points). Arrows represent the stages shown in **(D)** and **(E)**, respectively. Right plot: for each embryo, the CM area/Trunk Length was normalized to the median ratio of control embryos. Blue dots: *Fat1^+/+^; Gdnf^LacZ/+^* (n=21); Red dots: *Fat1^-/-^; Gdnf^LacZ/+^* (n=11). Underlying data are provided in S1 Data. Scale bars: 500 μm.

### *Fat1* controls spreading of the GDNF-expressing CM muscle

We next took advantage of the fact that the CM is a selective source of GDNF, thus offering an excellent marker to follow development of this muscle [39,42]. Alterations of the CM muscle shape resulting from disrupted *Fat1* functions can be visualized by following β-galactosidase activity in embryos carrying a *Gdnf^LacZ^* allele (Fig 1D, E). We therefore produced embryos carrying one copy of the *Gdnf^LacZ^* allele in wild type or *Fat1^-/-^* contexts and performed staining with Salmon Gal, a substrate more sensitive than X-gal, adapted to the low level of *Gdnf* expression (see experimental procedures). *Gdnf^LacZ^* expression can be detected as early as E11.5, prior to the emergence of the CM, at the level of the plexus mesenchyme (at fore and hindlimb levels), where it serves to guide motor axons and instruct them of their identity [39,42,54]. *Gdnf* expression is then detected in the CM and in the underlying Latissimus Dorsi (LD) muscles, as they emerge (around E12.0) from the brachial plexus [14,39]. The LD muscle is not visible on our pictures because it is hidden by the CM, but can be recognized on embryo sections. From that stage onward, these muscles progress by migrating under the skin in a posterior direction, radiating from their point of origin. As development proceeds, the area occupied by CM progenitors expands (and can be viewed through the skin by transparency in whole embryos). We focused on the time window when most of the subcutaneous progression is occurring (E12.0-E12.75). To analyze the rate of expansion, the *Gdnf^LacZ^*-positive area (white dotted area in Fig 1D, E) was plotted relative to the trunk length, used as a value that increases regularly as the embryo grows, thus reflecting the stage of development. At any stage examined, the area covered by *Gdnf^LacZ^*-positive cells is smaller than in control embryos (Fig 1D-F). The rate of CM expansion is significantly reduced in *Fat1^-/-^* embryos, with a median area reduced to 66% of controls. As seen with *MLC3F-2E,* this effect also appears more pronounced in the ventral part of the CM. Furthermore, staining intensity in the *Gdnf^LacZ^*-positive zone behind the progression front appears reduced in *Fat1* mutants (compare intensity along the vertical blue dotted line in Fig 1E). Rather than reflecting a reduction in the level of *Gdnf^LacZ^* expression per progenitors, this effect appears to result from a reduction in the density of *LacZ*-expressing cells in this front of migration.

This effect on *Gdnf* expression can also be observed in the context of the other null allele of *Fat1 (Fat1^LacZ^),* by following *Gdnf* expression by *in situ* hybridization on embryo sections (S2A Fig). In this context, reduced thickness of the *Gdnf*-expressing cell layer is observed in posterior CM sections of *Fat1^LacZ/LacZ^* embryos, reflecting a reduced amount of *Gdnf*-expressing cells. The reduced density in muscle progenitors in constituting the CM is also visualized by following markers of myogenic cells or subsets of migratory muscle populations such as *MyoD, Six1,* and *Lbx1* on sections at posterior levels (S2B Fig). A reduced *Gdnf* expression level was also detected at the level of the plexus mesenchyme in *Fat1^LacZ/LacZ^* embryos (S2A Fig). Overall, we confirm using two independent alleles that *Fat1* is required for the development of the CM muscle.

### *Fat1* ablation disrupts innervation of the CM muscle

The CM muscle represents an excellent example in which to study the coupling between muscle morphogenesis and neuronal specification. Given the strong effect of *Fat1* loss-of-function on expansion and differentiation of the CM muscle, we next wondered if changes could also be observed in the pattern of innervation. E12.5 control and *Fat1* mutant embryos were therefore stained by whole-mount immunohistochemistry with an anti-neurofilament antibody, and visualized after clearing in Benzyl-benzoate (Fig 2). Embryos were cut in half and flat-mounted for imaging. Motor axons innervating the CM muscle can be recognized on the embryo flank by their horizontal progression, as they intersect the vertically orientated thoracic nerves. After initial imaging of flat-mounted embryos halves (Fig 2A, B, top and middle images), all inner structures including thoracic nerves were manually removed to better distinguish CM axons (Fig 2A, and B; bottom pictures). CM motor axons cover an area with a shape very similar to that occupied by *Gdnf^LacZ^* expressing cells. This shape was affected by *Fat1* loss-of-function in a similar way as was the *Gdnf^LacZ^* expressing region. At comparable stages, the area covered by CM axons was smaller in *Fat1^-/-^* embryos, which exhibited shorter CM axons than wild types (Fig 2A, and B). As seen with muscle markers, the dorsal part of the muscle appears less affected. In the ventral muscle, mutant motor axons were shorter, with an apparent lower density of axons bundles than in controls (inserts in Fig 2A, bottom images). Throughout the period considered, the subcutaneous expansion of CM-innervating axons is fast and dynamic. It is therefore best represented by showing two consecutive stages, and quantified by measuring the area covered by CM-axons, plotted relative to a reference structure (such as the length of the T10 nerve, black dotted line) used as an indicator of developmental age (Fig 2C). In wild type embryos, the area covered by CM axons expanded steadily, covering the embryo flank in little more than half a day. In contrast, the rate of progression of CM innervation is reduced by approximately two fold in *Fat1^-/-^* embryos compared to controls. Similar observations can be made with the other null allele (*Fat1^LacZ^*, S3 Fig). In both mutants, in contrast to the abnormal behavior of CM innervating axons, most other limb innervating nerves are preserved and appeared unaffected in *Fat1^-/-^* embryos (Fig 2). There was no obvious ectopic nerve corresponding to the supernumerary muscle in the scapulohumeral region. Thus, loss of *Fat1* functions appears to predominantly affect the development of axons innervating the CM, matching the pronounced effect on muscle spreading and differentiation.

**Fig 2:**
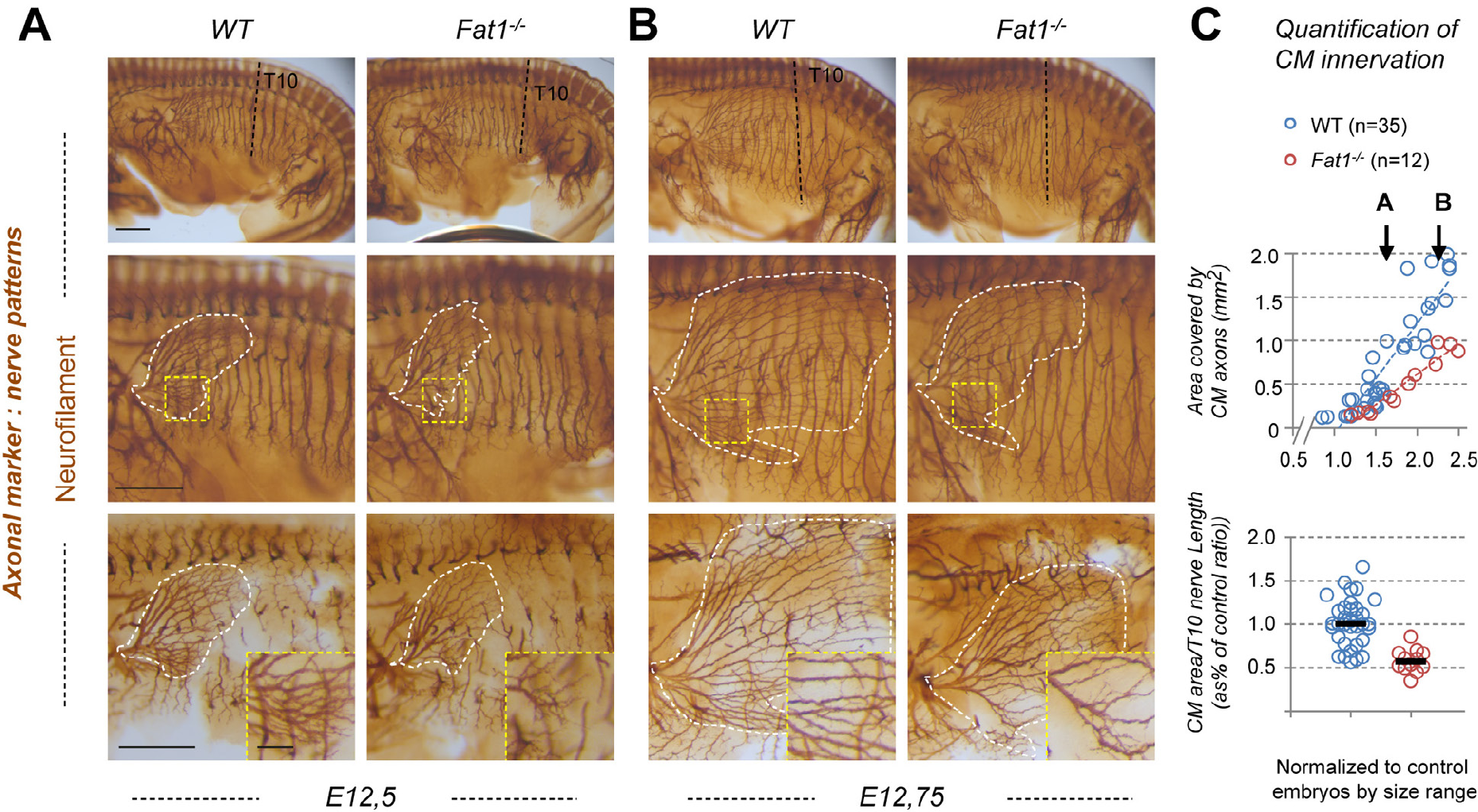
*Fat1* knockout alters motor innervation of the Cutaneous Maximus muscle. **(A, B)** The nerve pattern was analyzed by immunohistochemistry with antibodies against Neurofilament (2H3) in E12.5 **(A)** to E12.75 **(B)** wild type and *Fatl^-/-^* embryos. Embryos were cut in half, cleared in BB-BA, and flat-mounted. Upper panels are low magnification images of the left flank, showing the whole trunk. Lower panels show high magnification views of the area containing the CM muscle. The area covered by CM innervated axons is highlighted in white (middle panels). Axons of vertically oriented thoracic spinal nerves have been manually removed by dissection in the lower panels to improve visibility of CM axons. Inserts in the lower panels represent higher magnification of the area in the yellow squares. **(C)** Quantifications of the relative expansion of the area covered by CM-innervating axons. Upper plot: for each embryo side, the area covered by CM-innervating axons is plotted relative to the length of a thoracic nerve (T10, from dorsal root origin to ventral tip). Arrows point the stages of representative examples shown in **(A)** and **(B)**. Bottom plot: for each embryo, the CM-innervated area/T10 Length was normalized to the median ratio of control embryos, by size range. Blue dots: *Fatl^+/+^* (n=35, same sample set as in controls of S3 Fig); Red dots: *Fatl^-/-^* (n=12). Underlying data are provided in S1 Data. Sale bars: 500 m (large images); 100 μm (inserts in lower panels).

### Topographical organization of the CM muscle

A number of important features emerge when carefully comparing *MLC3F-2E^+^* embryos and *Gdnf^LacZ^* embryos during the developmental progression of the CM muscle in wild type embryos. For clarity in the following description, it is necessary to recall a few notions of orientation. Since the muscle progresses from anterior to posterior (see Fig 1, Fig 3A), the front of muscle progression is located posteriorly, whereas the rear corresponds to the point of origin of the muscle, at the brachial plexus, located anteriorly. When comparing *MLC3F-2E^+^* embryos and *Gdnf^LacZ^* embryos at similar stages (compare Fig 1A with 1D, and 1B with 1E), the area occupied by *Gdnf^LacZ^* cells (marked with a white dotted line) appears larger than the area occupied by *MLC3F-2E^+^* muscle fibres (marked with the red line). This *MLC3F-2E^+^* CM-differentiation has a specific fan-like shape, in which multinucleated muscle fibers extend from a narrow zone at the anterior origin, to a posterior side distributed along a wider dorso-ventral extent (Fig 1A, B). Since progression occurs at this posterior front (called front of fiber elongation), this indicates that multinucleated fibers elongate by adding new nuclei at the posterior side. This posterior front of fiber elongation appears to be situated approximately in the middle of the muscle, at a regular distance from the even more posterior front of progression of the *Gdnf^LacZ+^* area, likely composed of migrating muscle progenitors (Fig 3A). Nevertheless, when carefully observing X-gal stained *MLC3F-2E^+^* embryos, one can also distinguish, beyond the front of multinucleated fiber elongation, some *LacZ^+^* nuclei expressing β-Galactosidase at lower levels, within an area matching in size and shape the *Gdnf^LacZ+^* area (white dotted line, Fig 1). This suggests that the posterior half of the CM muscle (between the two fronts) is essentially occupied by *Gdnf^LacZ^*-positive migrating myogenic progenitors, and a few scattered mono-nucleated *MLC3F-2E^+^* myocytes. In contrast, the anterior half of the muscle contains elongating *MLC3F-2E^+^* fibers and *Gdnf^LacZ^*-positive cells. This was confirmed by analyzing serial sections of *Gdnf^LacZ/+^* embryos by immunohistochemistry, following β-galactosidase, the sarcomeric protein Myh1 in muscle fibers, and the muscle progenitor marker Pax7 (Fig 3B, C). Throughout the extent of the CM muscle (not considering the plexus mesenchyme region), we found that *Gdnf^LacZ^* was co-expressed with Pax7, confirming that it labels progenitors (Fig 3B), whereas it is not co-expressed with the sarcomeric protein Myh1 (Fig 3C inserts). Thus, *Gdnf^LacZ^* can be used as a specific marker of CM (and LD) progenitors. When following the CM muscle on serial sections (Fig 3C), we confirmed that only the anterior and middle sections contained a mixture of Pax7+*/Gdnf^LacZ^+* progenitors and Myh1+ fibers, whereas in the posterior sections, the CM only contained Pax7+*/Gdnf^LacZ^+* progenitors, but no fibers. Thus, there is a physical separation between the front of progenitor migration, and the front of differentiation, where new differentiating myocytes are being added to growing muscle fibers on their posterior side.

**Fig 3:**
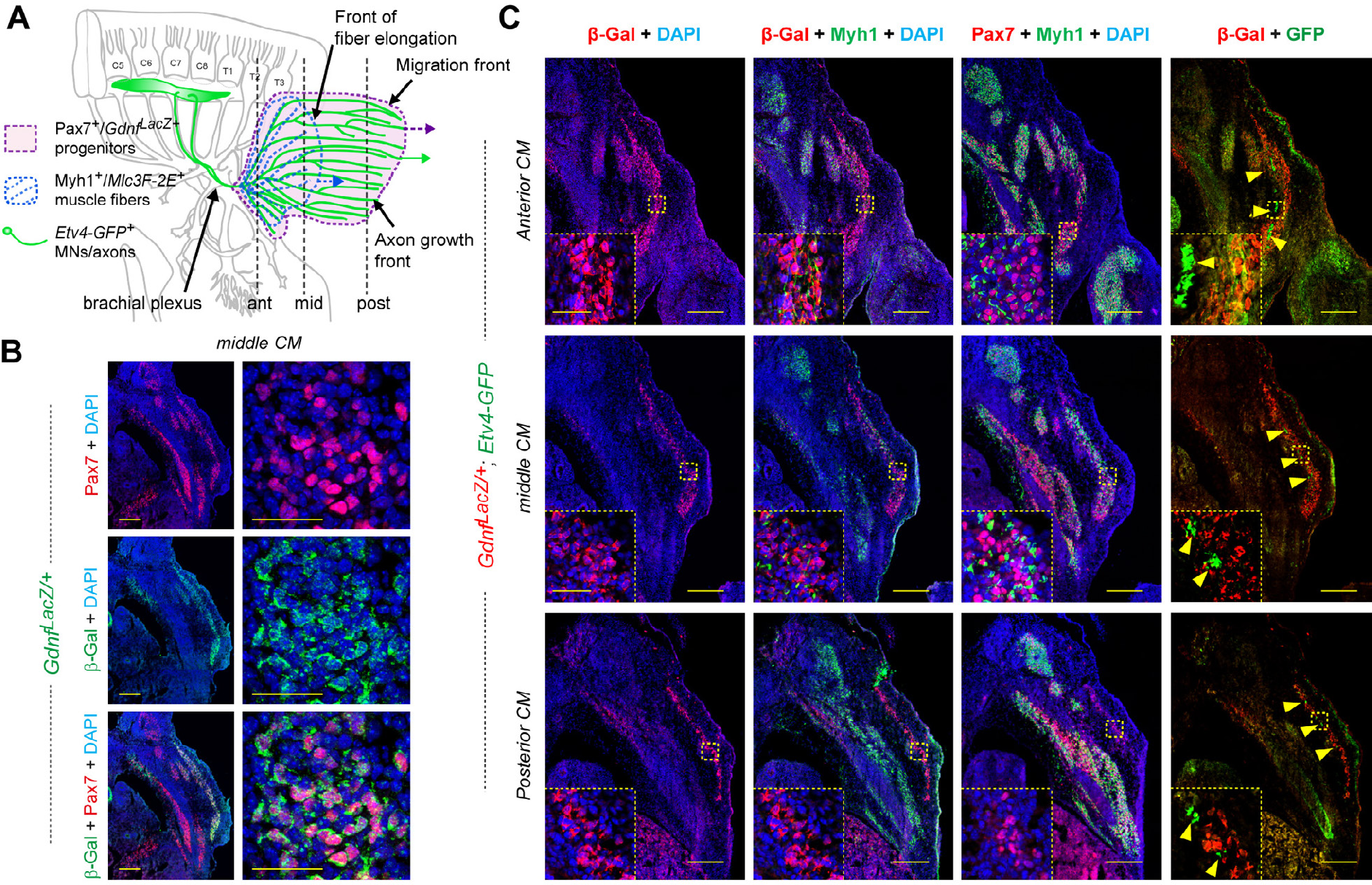
Topographic organization of myogenesis and nerve pattern in the CM muscle. **(A)** Scheme representing the shape of the CM muscle seen from the size of an embryo, featuring the area covered by *Gdnf^LacZ+^* muscle progenitors in purple, *MLC3F-2E^+^* muscle fibers represented in blue, and CM-innervating axons represented in green, indicating (vertical lines) the level corresponding to sections shown in **(B)** and **(C)**. **(B)** Cross section of an E12.5 *Gdnf^LacZ/+^* embryo, at the middle CM level, immunostained with antibodies to Pax7 (red) and β-galactosidase (green), showing that *Gdnf^LacZ^* is expressed in Pax7^+^ progenitors. **(C)** Cross sections of an E12.5 *Gdnf^LacZ/+^; Etv4-GFP^+^* embryo, at the anterior CM (top pictures), middle CM (middle row), and posterior CM (bottom row) levels, immunostained with antibodies to Pax7 (red), β-galactosidase (red), Myh1 (Type I myosin heavy chain, green), GFP (green), and with DAPI (blue). At each level, three neighboring sections of the same embryo were used with the indicated antibody combinations. In **(C)**, inserts show high magnifications of the area highlighted with the yellow dotted square. Scale bars: Low Magnification pictures in C: 200 μm; Inserts in C: 40 μm; High magnification (right) pictures in B: 40 μm; Low magnification pictures (Left) in B: 200 μm.

Interestingly, the shape of the area covered by CM-innervating motor axons (Fig 2) also appears more similar to the shape covered by *Gdnf^LacZ^-expressing* progenitors than to the shape the area covered by differentiated fibers (as visualized with the *MLC3F-2E* transgene) (Fig 1). The population of spinal motor neurons innervating the *Gdnf*-producing muscles CM (Fig 3A) and LD is characterized by expression of the transcription factor Etv4 [39,41]. We therefore took advantage of an *Etv4-GFP* transgene [60], in which GFP expression reproduces that of *Etv4* and enables detection of the corresponding axons, to follow CM-innervating motor axons on serial sections of *Gdnf^LacZ^; Etv4-GFP^+^* embryos (Fig 3A, C). GFP-positive axons can be seen throughout the extent of the CM, initially running as large bundles located along the interior side of the muscle in anterior sections, and progressively detected as smaller bundles posteriorly (Fig 3C). Even in the posterior-most sections, in which Myh1^+^ fibers are no longer detected in the *Gdnf^LacZ^*-positive CM sheet, GFP-positive axon bundles are found intermingled with *Gdnf^LacZ^* progenitors. Thus, *Etv4-GFP^+^* motor axons cover the entire zone enriched in *Gdnf^LacZ^*-expressing progenitors, as previously observed [39]. These observations imply that the front of migration contains *Gdnf^LacZ^* progenitors and distal tips of *Etv4-GFP^+^* motor axons, which appear to progress hands in hands, but not differentiated fibers. Thus, there is a significant topographical separation between the front of progenitor migration and axonal elongation, and the front of progression of muscle fiber elongation, where new myocytes are added to growing muscle fibers on their posterior side. These observations raise the interesting possibility that the speed and direction of CM muscle fiber elongation could be influenced by the direction/speed of progenitor migration, of axon elongation, or by a combination of both. In addition, the simultaneous progression of CM progenitors and *Etv4-GFP^+^* motor axons raise the issue of determining whether axons or muscle progenitors influence most the progression of CM expansion, and the subsequent expansion of muscle fibers.

### *Fat1* is expressed in multiple cell types delineating the CM neuromuscular circuit morphogenesis

We next asked in which cell type *Fat1* is required to exert the function(s) underlying this complex event of CM muscle and nerve morphogenesis. Although we previously showed that *Fat1* is required in the myogenic lineage to modulate myoblast migration polarity, the consequences of *Fat1* ablation driven in trunk myoblasts by *Pax3^cre^* were milder than in constitutive knockouts [14], suggesting that *Fat1* might be required in other cellular components of the circuit for its muscle patterning function. We therefore first analyzed *Fat1* expression during CM development, focusing on all the cell types involved, including muscles, surrounding connective tissues, and motor neurons. *Fat1* expression was followed either with anti-β-galactosidase antibodies on serial sections of *Fat1^LacZ/+^* embryos (Fig 4A, C, S3 Fig, and S4 Fig), by anti-Fat1 immunohistochemistry on sections of *Gdnf^LacZ/+^* embryos (Fig 4B, D, in which the β-galactosidase pattern reproduces *Gdnf* expression), or by *in situ* hybridization with a *Fat1* RNA probe (Fig 5). As previously reported [14], in addition to its expression in migrating myogenic cells (with *Fat1^LacZ^* expression detected in both Pax7-positive progenitors and Myh1-positive muscle fibers, Fig 4C), we also found that *Fat1* was expressed in mesenchymal cells surrounding the CM muscle, thus constituting a sheet of *Fat1* expressing cells through which the CM muscle expands (Fig 4C, D; see also [14]). Similarly, Fat1 protein is detected not only in *Gdnf^LacZ^*-positive muscle precursor cells (Fig 4D) but also in cells surrounding the layer of *Gdnf^LacZ^*-expressing progenitors (Fig 4D). β-galactosidase staining intensity in *Fat1^LacZ/+^* embryos appears more intense in the subcutaneous mesenchymal layer at the level of posterior CM sections than it is at anterior levels (Fig 4C and S3B). This expression was preserved in genetic contexts leading to depletion of migratory muscles, as evidenced by the robust *Fat1^LacZ^* expression detected in embryos in which myoblast migration is abrogated, such as *Met* [61,62] or *Pax3* [63–65] mutants (S5A Fig, and S5B Fig). In the latter case, following cells derived from the *Pax3^cre^* lineage using the R26-YFP reporter [66] (S5B Fig), reveals that most of the Fat1-β-galactosidase fusion protein detected in the dorsal forelimb remains unchanged, even in absence of the YFP^+^ myogenic component in *Pax3^cre/cre^; Fat1^LacZ/+^* compared to *Pax3^cre/+^; Fat1^LacZ/+^* embryos (the remaining YFP^+^ cells correspond to Schwann cells along the nerves). Together, these findings indicate that a large part of *Fat1* expression surrounding or within the CM muscle corresponds to mesenchymal cells (or connective tissue).

**Fig 4:**
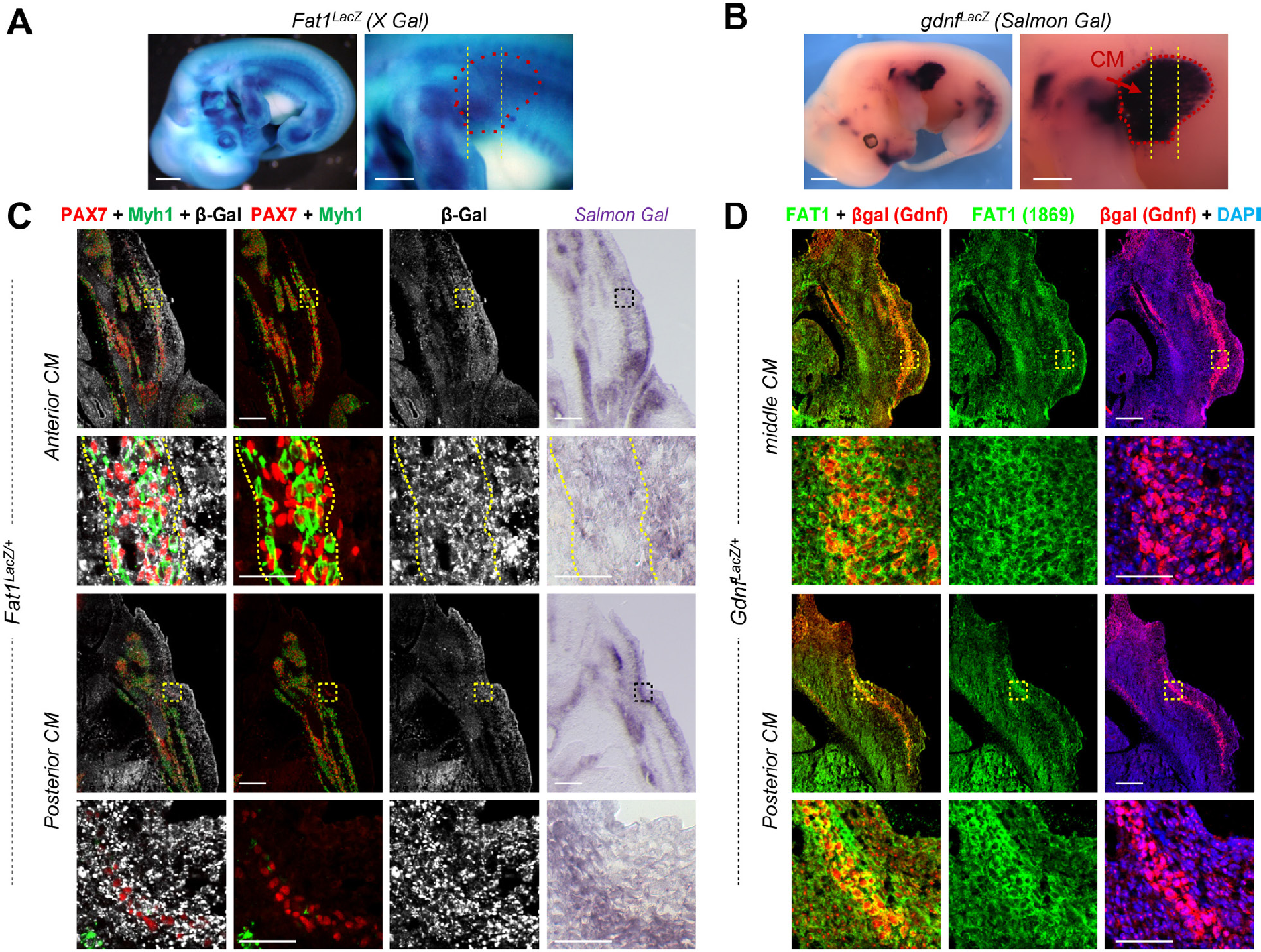
*Fat1* is expressed in CM progenitors and the surrounding subcutaneous mesenchyme. **(A)** *Fatl* expression is visualized in an E12.5 *Fatl^LacZ/+^* embryo by X-gal staining. Left panel: whole embryo picture; right panel: higher magnification of the forelimb and flank region in which the CM spreads. In the right panel, the approximate CM shape is highlighted by red dotted lines, and the level of sections shown in **(C)** is indicated by vertical lines. **(B)** *Gdnf* expression is visualized in an E12.5 *Gdnf^LacZ/+^* embryo by Salmon gal staining. Left panel: whole embryo left side view. Right panel: higher magnification of the upper forelimb and flank region, showing that the CM exhibits a high level of *Gdnf^LacZ+^* expression (highlighted with red dotted lines). The level of sections shown in **(D)** is represented by vertical bars. (C) Cross sections of an E12.5 *Fatl^LacZ/+^* embryo at anterior and posterior CM levels were immunostained with antibodies against Pax7 (red), Myh1 (green) and β-galactosidase (white). The right panels show neighboring sections of the same *Fatl^LacZ/+^* embryo in which β-galactosidase activity was revealed by Salmon-Gal staining. **(D)** Comparison between expression of *Gdnf^LacZ^* (visualized with an anti-β-galactosidase antibody (red)) and that of Fat1 (green, Ab FAT1-1869 Sigma) on two cross-sections of an E12.5 *Gdnf^LacZ/+^* mouse embryo at middle and posterior CM levels as indicated in **(B)**. Fat1 protein is detected both within and around the *Gdnf^LacZ/+^* CM progenitors. Scale bars: (A, B): 1 mm (left), 500 μm (right); (C, D): 200 μm (low magnification), 50 μm (high magnification).

**Fig 5:**
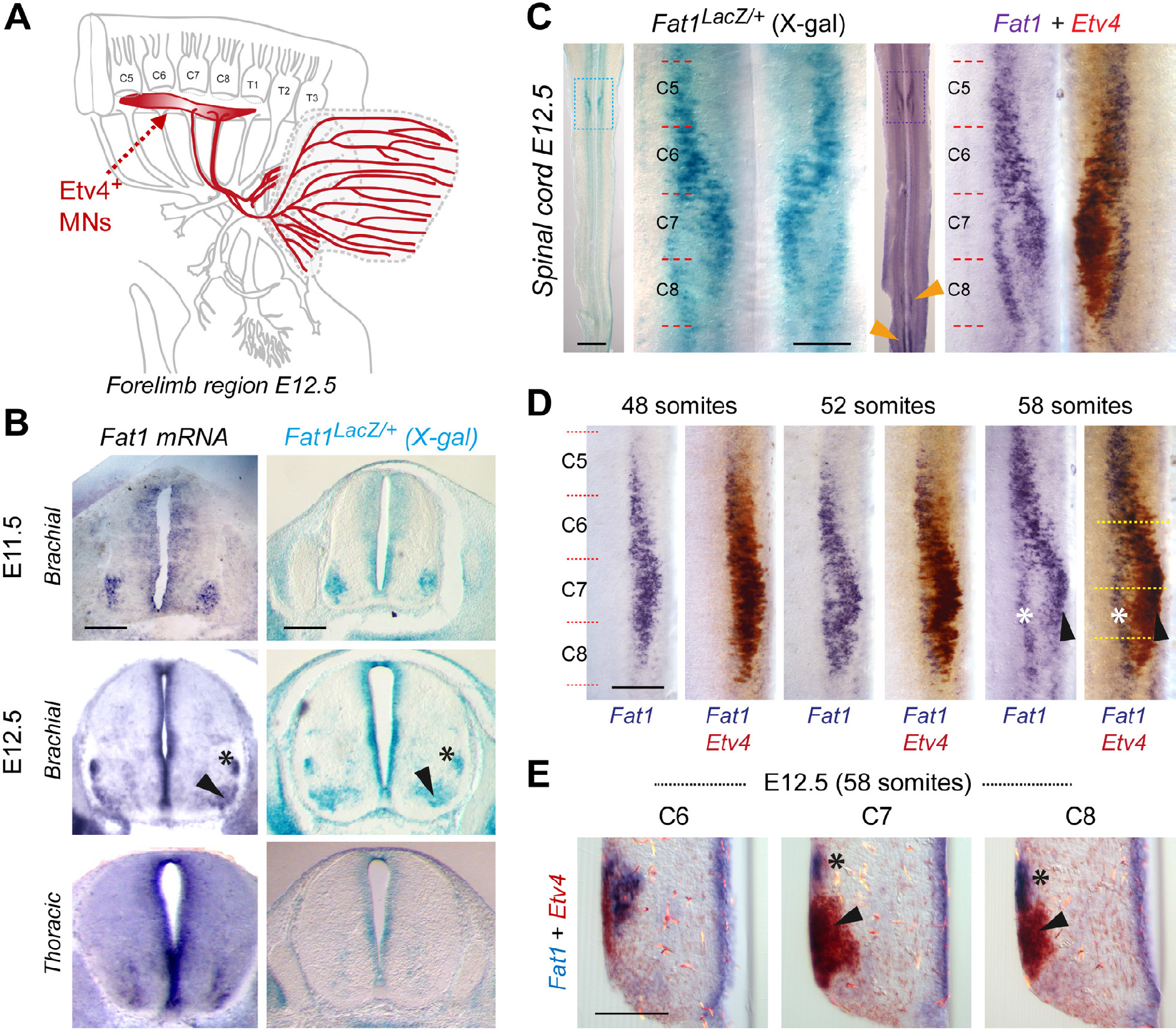
*Fat1* is expressed in subsets of brachial MN pools including CM-innervating *Etv4^+^* MNs. **(A)** Scheme representing the brachial region of a mouse embryo at E12.5 with the C4-T2 portion of the spinal cord, the corresponding spinal nerves and their projections to the forelimb, with Etv4+ MNs and their axons highlighted in red, whereas the target muscles CM and LD are delineated in blue (the LD being underneath the CM). **(B)** *Fatl* expression in the mouse brachial spinal cord is shown by *in situ* hybridization in wild-type embryo sections (left panels) and by X-gal staining on sections of *Fatl^LacZ/+^* spinal cords at E11.5 (top) and E12.5 (bottom), showing expression in all neural progenitors in the ventricular zone, and in pools of MNs, visible as one single cluster at E11.5, and two separate pools (arrowhead and asterisk) at E12.5. **(C)** *Fatl* expression in the mouse brachial spinal cord at E12.5 is shown through an X-gal staining of a *Fatl^LacZ/+^* spinal cord (left) or a double in situ hybridization with *Fatl* (purple) and *Etv4* (brown) RNA probes on a wild type spinal cord. *Fatl* expression is detected in *Etv4-*expressing MN pools (arrowheads) but also expressed in a distinct dorsal column (asterisk). Spinal cords are flat-mounted, such that the ventral midline is seen as a vertical line in the middle, and motor columns are seen on both sides. For both stainings, the entire spinal cord is shown on the left, and a magnification of the brachial region is shown on the right (corresponding to the delineated zone). Left and right sides of the in situ hybridization panel show a mirror image of the same spinal cord side before and after developing the brown *(Etv4)* reaction. **(D)** double in situ hybridizations with *Fatl* (purple) and *Etv4* (brown) RNA probes on wild type spinal cords at three successive time points, 48 somites (E11.5), 52 somites (E12.0) and 58 somites (E12.5), showing the left side of the brachial spinal cord, after developing Fat1 only (purple, left), or after developing the second reaction, in brown (right). **(E)** cross section on the 58 somite spinal cord shown in (**D**, right panel), at the three levels indicated by dotted line in **(D)**, showing the partial overlap (arrowhead) between *Fatl* and *Etv4* expression, and the dorsal pool of MNs expressing Fatl only. Scale bars: (B) 200 μm; (C): low magnification: 1mm; (C) high magnification: 200 μm; (D) 200 μm; (E) 100 μm.

**Fig 6:**
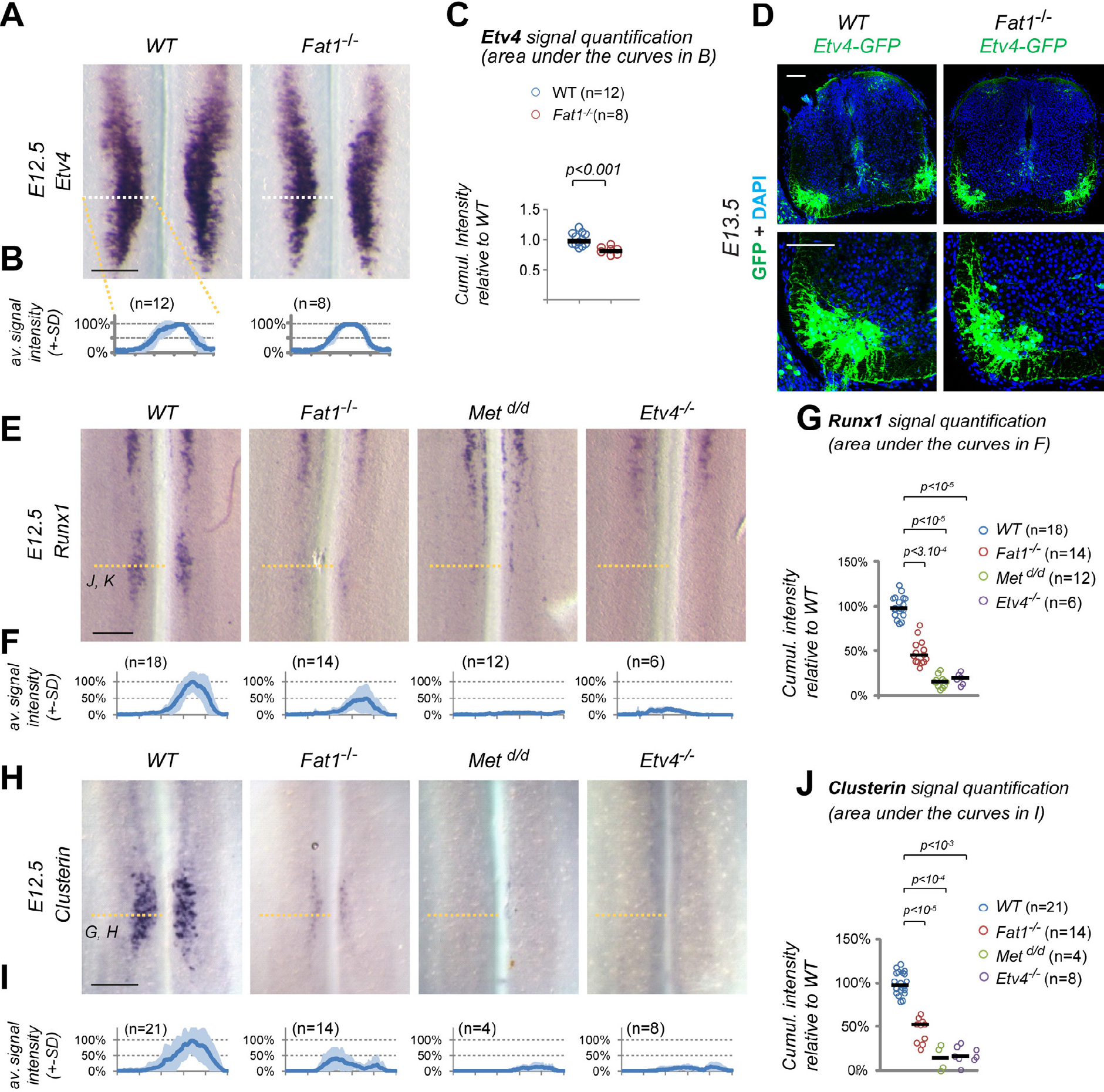
*Fat1* knockout alters the specification of CM Motor neuron pools. **(A)** *Etv4* expression was analyzed by in situ hybridization in E12.5 wild type and *Fatl^-/-^* embryos. The images represent flat-mounted spinal cords in the brachial region. **(B)** Quantifications of *Etv4* signal: each plot represents the average signal distribution (± standard deviation in light blue) measured on the indicated number of spinal cord sides along the white dotted line in each image in **(A)** *(Fatl^+/+^* (n=12; this set of controls includes the same samples as those shown in S7 Fig); *Fatl^-/-^*(n=8). **(C)** Quantifications and statistical analyses of the sum of signal intensity corresponding to the area under the curves in plots shown in **(C)**: each dot represents the sum of *Etv4* intensity for each spinal cord side, the number of samples being indicated (the two sides of each embryo considered independent). (B-C): Underlying data are provided in S1 Data. **(D)** Sections of spinal cords from E13.5 *Fatl^+/+^;Etv4-GFP* and *Fatl^-/-^;Etv4-GFP* embryos, were stained with antibodies against GFP, and with DAPI. **(E-J)** Analysis by ISH of *Runxl* **(E)** and *Clusterin* **(H)** expression in flat-mounted brachial spinal cords from E12.5 wild type, *Fatl^-/-^, Met^d/d^* and *Etv4^-/-^* embryos: Expression of *Clusterin* and *Runxl* in the C7-C8 segments is lost in both *Etv4* and *Met* mutants, and severely reduced in *Fatl^-/-^* spinal cords, whereas the rostral domain of *Runxl* expression is independent of *Met, Etv4* and *Fatl.* Quantifications of *Runxl* **(F, G)** and *Clusterin* **(I, J)** signal intensity: each plot in **(F, I)** represents the average signal distribution (± standard deviation in light blue) measured on the indicated number of spinal cord sides along the orange dotted line in each image above (with the corresponding genotype), in **(F)** for *Clusterin* and **(I)** for *Runxl.* **(F, G)** *Clusterin* probe: *Fatl^+/+^* (n=21); *Fatl^-/-^* (n=14); *Met^d/d^* (n=4); *Etv4^-/-^* (n=8); **(I, J)** *Runxl* probe: *Fatl^+/+^* (n=18); *Fatl^-/-^* (n=14); *Met^d/d^* (n=12); *Etv4^-/-^* (n=6)). Underlying data are provided in S1 Data. **(G, J)** Quantifications and statistical analyses of the sum of signal intensity corresponding to the area under the curves in plots shown in **(F)** and **(I)**, respectively: each dot represents the sum of *Runxl* or *Clusterin* intensity for each spinal cord side, the number of samples being indicated. Underlying data are provided in S1 Data. Scale bars: 200 μm (A, E, H); 100 μm (D).

The population of spinal motor neurons innervating the *Gdnf*-producing muscles CM and LD is characterized by expression of the transcription factor Etv4 (Fig 5A) [39,41]. Interestingly, in addition to its peripheral expression, we also detected *Fat1* expression in groups of MNs at brachial levels encompassing the pools of *Etv4*-expressing MNs (Fig 5C-E). Aside from expression in neural precursors in the ventricular zone all along the dorso-ventral axis (Fig 5B, S6C Fig), this brachial MN column was the main site of high *Fat1* expression in the spinal cord (Fig 5C). *Fat1* expression was also detected in a column of ventral neurons at thoracic levels (S6A Fig), and with later onset, in subsets of lumbar and sacral MNs (Fig 5C (orange arrowheads) and S4B Fig). The overlap between *Etv4* and *Fat1* expression in brachial MNs was maximal at E11.5 (Fig 5D), whereas additional groups of *Fat1*-positive; *Etv4*-negative (*Fat1*-only) neurons become detectable in dorsal positions at E12.5 (Fig 5D, E). At that stage, the CM MN pools have completed their shift in cell body position in the spinal cord [41], resulting in the dorso-ventral split of *Fat1* expression domain at C7-C8 levels into a dorsal *Fat1*-only pool (Fig 5D, E, asterisk), and a ventral, *Fat1^+^/Etv4^+^* pool (Fig 5D, E, arrowheads), matching position of CM MNs.

Given the *Fat1/Etv4* co-expression, we asked whether *Fat1* could be a transcriptional target of Etv4, or whether its expression was dependent on factors acting upstream of *Etv4*, such as GDNF and HGF [39,41,47]. *Fat1* expression only appeared modestly reduced in shape in *Etv4^-/-^* and *Gdnf^-/-^* spinal cords and unchanged in *Met^d/d^* spinal cords at E11.5 (S6A Fig, S6B Fig). These data indicate that *Fat1* induction occurred independently of HGF/Met, GDNF and Etv4, in spite of the subtle changes in shape of *Fat1*-expressing columns in *Etv4* and *Gdnf* mutants. Nevertheless, at E12.5, after the dorsoventral split into two *Fat1-*expressing columns, the ventral *Fat1* expressing pool was missing in *Etv4^-/-^, Gdnf^-/-^* and *Met^d/d^* spinal cord (S6A Fig, S6B Fig), consistent with previously reported changes in the fate of CM MNs [39,41,47]. In contrast, the dorsal column appeared increased in *Etv4^-/-^* and *Gdnf^-/-^* spinal cord (S6B Fig), also consistent with the altered positioning of some CM neurons [39,41]. This dorsal column appeared reduced in *Met^d/d^* spinal cord, possibly resulting from the onset of enhanced MN death in the absence of the target muscle [46,47,60]. Altogether, these data are consistent with *Fat1* being expressed in CM motor pools. The reiterated use of *Fat1* expression in several components of the GDNF/Etv4 circuit and the altered shape of CM muscle and innervations pattern in *Fat1* mutants raise the possibility that loss of *Fat1* functions might influence development of this neuromuscular circuit either by acting directly in MNs or as an indirect consequence of its role in muscle patterning.

### Loss of *Fat1* causes CM-innervating Motor neuron specification defects

We next asked whether the CM muscle phenotype and the changes in motor axon patterns observed in *Fat1* mutants were also associated with molecular defects in the corresponding spinal MN pools. The co-expression of *Fat1* with *Etv4* in MNs and the selective alteration in shape and nerve projections to the CM prompted us to focus on specification of the CM motor pools and to examine whether *Etv4* expression was altered in *Fat1* mutants. This analysis revealed that *Fat1* is dispensable for the establishment of *Etv4* expression domain in *Fat1^-/-^* spinal cord (Fig 6A-C, and S7A Fig). Using the *Etv4-GFP* transgene [60] also allowed detecting a near normal appearance of the GFP-positive motor columns in *Fat1^-/-^* spinal cords (Fig 6D). Nevertheless, analysis of signal intensity of *Etv4* mRNA detected a modest but significant lowering of *Etv4* signal intensity of around 20% (Fig 6B and C). We next asked whether such modest changes in *Etv4* levels may be sufficient to impact expression of some of its transcriptional targets (for example affecting low affinity but not high affinity targets).

We therefore analyzed expression of several markers of subsets of *Etv4^+^* neurons, which expression is dependent on *Etv4* activity, some of those being also affected by loss of its upstream regulator *Met,* a situation which we showed only partially alters *Etv4* expression [41,47]. We first studied *Sema3E* and *Cadherin 8,* two known Etv4 targets in the CM motor pool [41]. Consistent with previous reports [41], their expression is absent in *Etv4^-/-^* spinal cords, whereas it was reduced but not lost in *Met^d/d^* spinal cords (S7E Fig). *Sema3E* and *Cadherin 8* expression appeared unaffected in *Fat1^-/-^* spinal cords (S7E Fig), these two genes behaving as expected for robust “high affinity” Etv4-targets. We next studied expression of *Clusterin* and *Runx1*, two genes which we selected for their expression in the CM motor pool. *Runx1* is a transcription factor expressed in a rostral column of ventrally located neurons spanning C1-C6 [67], and in a separate pool at C7-C8 matching the CM subset of *Etv4^+^* MNs (Fig 6E and S5C Fig), where its expression was shown to require Met signaling [60]. *Clusterin* is a glycoprotein, known to accumulate in neurons after axotomy, injury, or in neurodegenerative diseases (such as ALS, Alzheimer), which was proposed to modulate cell death, autophagy and clearance of proteins aggregates or cell debris [68–71]. *Clusterin* expression in the developing spinal cord was restricted to a subset of MNs matching the position of the *Etv4^+^* CM pool (Fig 6H). In contrast to *Sema3E* and *Cadherin 8,* expression of both *Clusterin* and *Runx1* was completely abolished in *Etv4* and *Met* knockout spinal cords (Fig 6E-J). *Clusterin* and *Runx1* signal intensities were severely reduced in the C7-C8 MN pool in *Fat1^-/-^* spinal cords, although not as severely as in *Met* and *Etv4* mutants (Fig 6E-J). The severe effect on *Clusterin* and *Runx1* expression and mild effect on *Etv4* expression, detected at E12.5 in *Fat1* mutant spinal cords, is unlikely to result from increased cell death. First, *Runx1* is coexpressed with *Sema3E* (S6C Fig), which expression is unaffected by loss of *Fat1* (S7E Fig), indicating that the neurons that fail to express *Clusterin/Runx1* in *Fat1* mutants are still present and retain *Sema3E* expression. Second, naturally ongoing MN death only peaks at E13.5 in mice, one day after the stage analyzed here. Absence of muscle development in mutants of genes such as such as *Pax3* or *Met* was shown to have minor effect on MN numbers prior to the establishment of the trophic dependency of MNs for muscle-derived factors [47]. Thus, at E12.5, stage of our current analysis, cell death is unlikely to contribute to the effects of *Fat1* loss-of-function on *Clusterin/Runx1* expression. Thus their sensitivity to the absence of *Fat1* is consistent with *Clusterin* and *Runx1* expression being low affinity targets of Etv4, affected by subtle changes in *Etv4* levels. Altogether, these data confirm that loss of *Fat1* compromises the correct acquisition of molecular identity of the CM innervating MN pool, in addition to its effect on CM muscle growth and differentiation.

### *Fat1* ablation in mesenchymal cells, but not in MNs, causes drastic muscle shape patterning defects

There is a strong topological connection between altered CM muscle morphology observed in *Fat1* mutants and the selective defects in the corresponding MN population. This topology is even more puzzling when considering that *Fat1* is expressed in several cell types including *Etv4^+^* MN, along the GDNF/Etv4 circuit. Neural and muscular aspects of the phenotypes could represent the consequences of one single primary phenotype resulting from *Fat1* deletion in one single cell type. Given the known dependency of *Etv4* induction on GDNF [39,41,47], the neural phenotypes could arise as a consequence of the reduced amount of GDNF-producing cells. However, the discovery of *Fat1* selective expression in *Etv4^+^* MNs raises the alternative possibility that some aspects of these phenotypes may result from its ablation in MNs. Thus, the overall knockout phenotype could represent the cumulated consequences of phenotypes resulting from ablation of independent *Fat1* assignments in distinct cell types. To tackle this question, we used a conditional approach to selectively interfere with *Fat1* functions in a tissue-specific manner. This allowed assessing in which cell type it was necessary and sufficient to ablate *Fat1* functions to reproduce each aspect of the muscular and neural phenotypes observed in constitutive mutants. We combined our *Fat1^Flox^* conditional allele [14] with several Cre lines matching the different sites of *Fat1* expression (see S1 Table for descriptions): *Pax3-cre* knock-in for ablation in the premigratory myogenic lineage [64], *Olig2-cre* knock-in for MN precursors [72], *Prx1-cre* transgenic line for limb mesenchymal cells [73,74], and *Wnt1-cre* transgenic line in neural crest derivatives [75]. We first focused on the strong alterations of CM muscle appearance observed in the constitutive knockout. We followed muscle differentiation using the *MLC3F-nLacZ-2E* transgene. Although we previously described significant alterations of myogenesis resulting from *Fat1* ablation in the premigratory myogenic lineage (driven by *Pax3-cre* [14]), the appearance of the CM muscle in E14.5 *Pax3-cre; Fat1^Flox/Flox^* embryos was grossly normal, albeit with reduced density of differentiated muscle cells. This contrasts with constitutive *Fat1* knockouts, where the severe effect on CM shape persists at later stages, when the muscle has further extended to cover most of the trunk [14]. This discrepancy indicated that the effect on muscle growth caused by premigratory ablation in myogenic cells was attenuated at later stages. Thus *Fat1* function underlying proper CM development must therefore also be exerted in a distinct cell type.

Given the selective co-expression of *Fat1* and *Etv4* in the CM MN pool, and the fact that the front of CM expansion is led by both muscle progenitors and axons, we next asked if *Fat1* deletion in the corresponding motor neurons had an impact on CM muscle development. However, MN-specific *Fat1* deletion in *Olig2-cre; Fat1^Flox/Flox^; MLC3F-2E* embryos did not cause any detectable change in the appearance (S8A Fig) and expansion rate of the CM as assessed by measuring the CM area in X-gal stained embryos carrying the *MLC3F-2E-LacZ* transgene (S8B Fig). MN-specific mutants also lacked all the other muscle phenotypes observed in constitutive knockouts, with no ectopic muscle in the scapulohumeral region (Figure S8A), no myocyte dispersed in the forelimb (S8A Fig, B Fig), and an overall normal appearance at E14.5 (S9 Fig). Together, these observations establish that the progression of myogenic differentiation in the CM is not significantly influenced by *Fat1* activity in the pool of MNs innervating this muscle.

In contrast, we found that *Fat1* ablation driven in the limb and trunk mesenchyme by *Prx1-cre* led to a severe and robust change in the appearance of the CM muscle (Fig 7A, B) and of upper forearm muscles (S8A Fig, S8B Fig). This phenotype was already visible at E12.5, during the phase of CM posterior extension, with a significant reduction of the rate of CM expansion assessed by X-gal staining in *Prx1-cre; Fat1^Flox/Flox^; MLC3F-2E* embryos (Fig 7A, B). This leads to a phenotype as severe as that observed in constitutive knockouts as measured when comparing the growth rate of the CM differentiation area to that of body wall muscles (Fig 7B). Again, the ventral half of the CM was severely shortened, whereas spreading in the dorsal part appeared less affected (Fig 7A). Moreover, *Prx1-cre; Fat1^Flox/Flox^; MLC3F-2E* embryos consistently exhibited changes in forearm muscles, such as the appearance of ectopic muscles in the scapulohumeral area (S8A Fig), or a mild but significant increase in the amount of dispersed myocytes in the forelimb (S8B Fig). At E14.5, stage at which the CM is fully extended in the trunk, up to the point of emergence of the hindlimbs, *Prx1-cre; Fat1^Flox/Flox^* embryos exhibited a pronounced hypoplasia of the ventral CM, with only very few *MLC3F-2E-*positive fibers having successfully extended in random orientations (S9A Fig, S9C Fig), thus strongly recapitulating the changes seen in the germline deletion. Again, the dorsal CM appeared less affected in E14.5 *Prx1-cre; Fat1^Flox/Flox^;MLC3F-2E* embryos, although the density of *LacZ^+^* nuclei nevertheless appeared reduced compared to *Fat1^Flox/Flox^;MLC3F-2E* embryos (S9A Fig).

**Fig 7:**
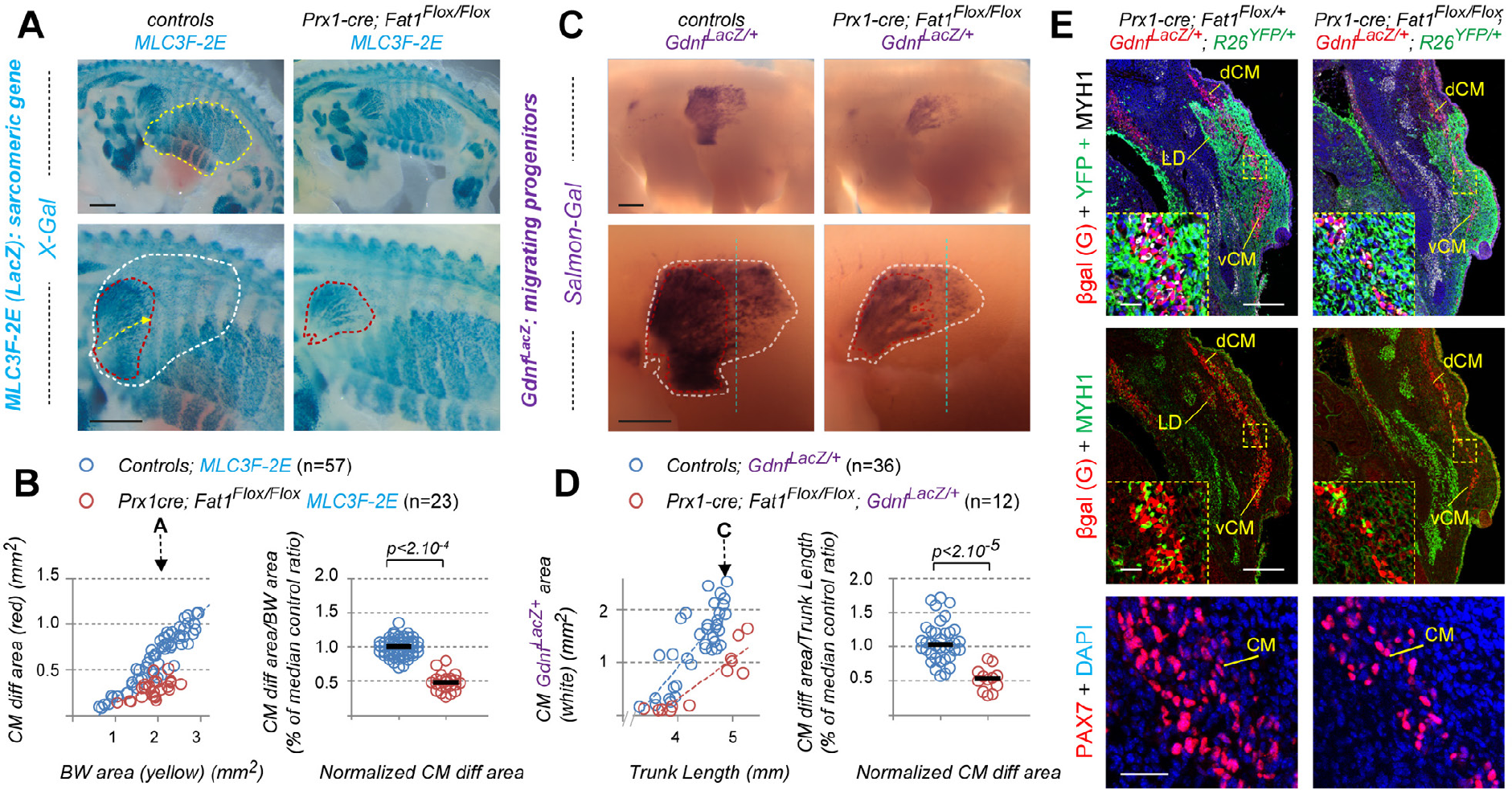
Mesenchyme-specific *Fat1* deletion non-cell-autonomously alters CM expansion. **(A, C)** Whole-mount β-galactosidase staining was performed using X-gal as substrate on embryos carrying the *MLC3F-2E* transgene **(A)**, or using Salmon-Gal as substrate on embryos carrying the *Gdnf^LacZ/+^* allele **(C)**, in the context of mesenchyme-specific deletion of *Fat1,* driven by Prx1-cre at E12.5. Top images show a side view of the whole flank of an embryo. Yellow dotted lines highlight the area occupied by body wall muscles. Lower images are higher magnification of the area in which the CM spreads. Red and white dotted lines correspond (as in Fig 1) to the areas covered respectively by *MLC3F-2E^+^* CM-fibers (red, **A**), and to the area covered by *Gdnf^LacZ+^* progenitors (white,**C**), respectively. **(B)** Quantification of the expansion rate of differentiated CM fibers. Left graph: For each embryo side, the area covered by differentiated CM fibers was plotted relative to the area occupied by body wall muscles. Right plot: for each embryo, the CM area/Body wall area was normalized to the median ratio of control embryos. Blue dots: *Fat1^Flox/Flox^; MLC3F-2E* (n=57, includes the same set of controls as S8C Fig); Red dots: *Prx1-cre; Fatl^Flox/Flox-^; MLC3F-2E* (n=23). Underlying data are provided in S1 Data. (D) Quantification of the expansion rate of the area occupied by *Gdnf^LacZ+^* progenitors. Left plot: For each embryo side, the area covered by *Gdnf^LacZ+^* progenitors was plotted relative to the trunk length. Right plot: for each embryo, the *Gdnf^LacZ+^CM* area/Trunk Length was normalized to the median ratio of control embryos. Blue dots: *Fat1^Flox/Flox^; Gdnf^LacZ/+^* (n=36, pooling respective littermates); Red dots: *Prx1-cre; Fatl^Flox/Flox-^; Gdnf^LacZ/+^* (n=12). Underlying data are provided in S1 Data. **(E)** Cross-sections of E12.5 *Prx1-cre; Fatl^FIox/+^; Gdnf^LacZ/+^; R26^YFP/+^ and *Prx1-cre*; Fatl^Flox/Flox^; Gdnf^LacZ/+^; R26^YFP/+^* embryos at equivalent rostro-caudal positions (caudal CM level), were immunostained with antibodies against GFP/YFP (green) to reveal the domain of *Prx1-cre* activity (green), against β-galactosidase (red), against Myh1 (white on top panels and inserts, green on middle panels), against Pax7 (bottom panels), and with DAPI (blue). The yellow dotted boxes indicate the areas magnified in inserts and in the bottom panels, in equivalent positions of the CM. Images show that lowered *Gdnf* levels represent a non-cell-autonomous consequence of lack of *Fat1* signaling in the mesenchyme of *Prxl-cre; Fatl^Flox/Flox^* embryos, and result from a reduced number of Pax7-GDNF-expressing progenitors cells, rather than from a lower level of *Gdnf^LacZ^* expression per cell. Scale bars: **(A, C)** 500 μm; (E) low magnification: 200 μm; inserts: 20 μm; lower panels: 40 μm.

Constitutive *Fat1* knockouts also exhibit abnormal morphology of subcutaneous muscles in the face, mostly visible at E14.5 [14]. Unlike the CM muscle, this group of facial subcutaneous muscles appeared unaffected in E14.5 *Prx1-cre; Fat1^Flox/Flox^; MLC3F-2E* embryos (S9 Fig), consistent with the lack of *Prx1-cre* activity in craniofacial mesenchyme [73,74]. Ablation of *Fat1* functions in the craniofacial mesenchyme was achieved using *Wnt1-cre* [75], which drives cre expression in all neural crest including cephalic neural crest, from which most craniofacial mesenchyme derives [76]. This led to profound morphological alterations in the appearance of facial subcutaneous muscles in *Wnt1-cre; Fat1^Flox/Flox^; MLC3F-2E* embryos, with a severely reduced fiber density and drastic changes in fiber orientation and position of fiber origins (S9 Fig). The same severe effect on morphology of facial subcutaneous muscles was also observed in E14.5 *Pax3^cre/+^; Fat1^Flox/Flox^* embryos (S9 Fig). This reflects the known activity of *Pax3-cre* in neural crest as well. By contrast, the observation of intact morphology of scapulohumeral muscles, normal rate of progression of the CM muscle, and lack of myocyte dispersion in the forelimb in *Wnt1-cre; Fat1^Flox/Flox^; MLC3F-2E* embryos (S8 Fig, S9 Fig), indicates that *Fat1* is dispensable in trunk neural crest to pattern trunk muscle development. Altogether, these data identify mesenchyme as a cell type in which *Fat1* signaling is required for muscle morphogenesis, with the trunk mesenchyme deriving from *Prx1-cre* lineage, and the craniofacial mesenchyme deriving from the *Wnt1-cre/*neural crest lineage. In contrast, these results establish that *Fat1* activity in MNs and trunk neural crest is dispensable for myogenic differentiation.

### *Fat1* ablation in mesenchymal cells non-cell-autonomously disrupts posterior spreading of GDNF-expressing CM progenitors

So far, we have established that *Fat1* is required in the mesenchyme for the progression of differentiation and fiber elongation in the CM. Since the delay in CM expansion observed in constitutive knockouts was associated with a reduced rate of expansion of the sheet of *Gdnf^LacZ^* expressing progenitors, we next asked if this progenitor progression was also affected, by bringing the *Gdnf^LacZ^* allele in the mesenchyme-specific mutant context, and performing Salmon Gal staining on whole-mounts embryos. As in knockouts, we observed an important reduction in the rate of progression of the area occupied by *Gdnf^LacZ^*-expressing progenitors (related to the evolution of trunk length) in *Prx1-cre; Fat1^Flox/Flox^; Gdnf^LacZ/+^* embryos compared to *Fat1^Flox/Flox^; Gdnf^LacZ/+^* embryos (Fig 7C, D). When comparing embryos of similar stage, the *Gdnf^LacZ^* sheet appeared truncated in the ventral region (vCM), and shorter in the dorsal CM (dCM), with an apparent reduction in staining density, visible by comparing staining intensity along a dorsoventral line positioned in similar position (blue dotted line in (Fig 7C,). This approach does not distinguish a reduction in expression level from a reduced number/density of cells expressing *Gdnf^LacZ^.* Therefore, to discriminate between the two options, we next analyzed the level of β-galactosidase protein (visualized by immunohistochemistry) driven by the *Gdnf^LacZ^* allele on control and mutant embryo sections. At middle and posterior CM levels, there was a clear reduction in thickness of the sheet of *Gdnf^LacZ+^* cells detected in a *Prx1-cre; Fat1^Flox/Flox^; Gdnf^LacZ/+^* embryo compared to a control embryo, resulting from a lowered number of stained cells rather than a reduced expression level per cells (Fig 7E, inserts). *Gdnf^LacZ^*-expressing cells however exhibited comparable β-galactosidase staining intensity, ruling out an effect on *Gdnf* expression levels. Staining with antibodies against Pax7 to mark progenitors, confirmed a reduction in the number of Pax7^+^ progenitors at comparable AP levels (Fig 7E). Thus, the reduced CM thickness in *Prx1-cre; Fat1^Flox/Flox^; Gdn^LacZ/+^* embryos results for a large part from a reduced amount of Pax7^+^ progenitors, and from a consequent reduction in the amount of differentiated fibers (highlighted with anti-Myh1 antibodies), compared to *Prx1^cre^; Fat1^Flox/+^; Gdnf^LacZ/+^* controls.

Our histological analysis of serial sections of mesenchyme-specific mutants and controls was done in a genetic context allowing lineage tracing of *Prx1-cre* activity with a *R26-YFP* reporter [66], to highlight cells in which cre-mediated recombination is occurring (Fig 8A). This context *(Prx1-cre; R26YFP; Gdnf^LacZ/+^)* allowed visualizing both reporters (β-galactosidase and YFP) simultaneously by immunohistochemistry on transverse sections spanning from the brachial plexus to the CM muscle, comparing *Fat1^Flox/+^* (Fig 8B) with *Fat1^Flox/Flox^* (Fig 8C) mutant settings (see also Fig 7E). This analysis confirmed that *Gdnf* expression domain is subdivided in two compartments: In posterior sections, *Gdnf^LacZ^*-expressing CM muscle progenitors detected throughout the length of the muscle do not derive from mesenchymal progenitors (Fig 7E, Fig 8). These myogenic cells appear to slide along and be surrounded by a territory of YFP-expressing connective tissue mesenchymal lineage composed of *Prx1-cre; R26YFP* cells, up to a dorso-ventral boundary corresponding to the limits of the *Prx1-cre* lineage [73] (Fig 7E, and Fig 8). In contrast, in anterior sections, at the level of the brachial plexus, *Gdnf^LacZ^*-expressing cells co-express β-galactosidase and YFP, indicating that the plexus component of *Gdnf^LacZ^* domain is constituted of mesenchyme-derived cells (Fig 8). Thus, *Gdnf* expression domain is constituted of two sub-domains of distinct developmental origins (myogenic and mesenchymal, respectively), which are anatomically connected at the position of origin of migration of the CM (and LD progenitors).

**Fig 8:**
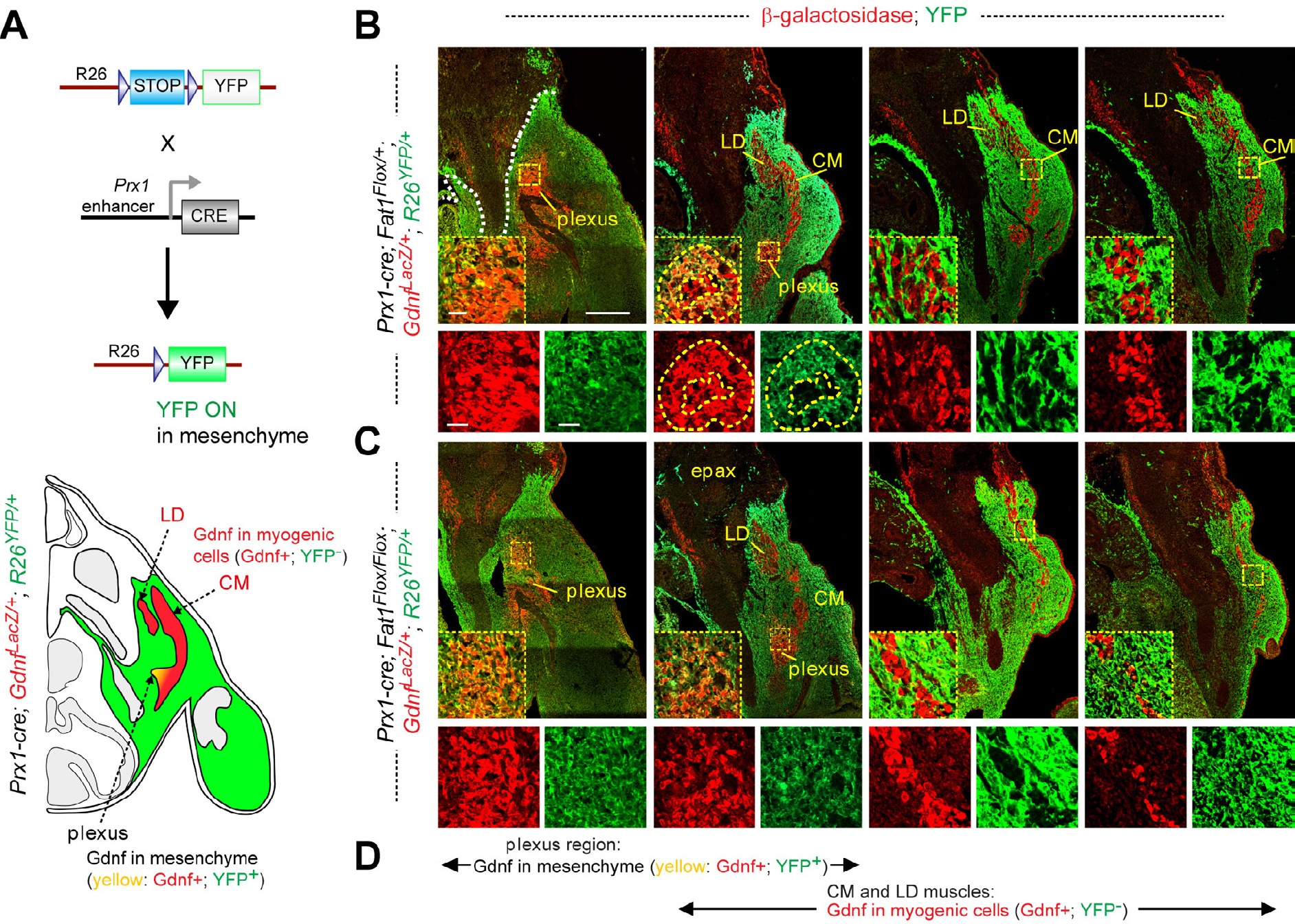
Mesenchymal-*Fat1* is required for expansion of the myogenic component of *Gdnf* expression domain, but dispensable for *Gdnf* expression in plexus mesenchyme. **(A) Top:** Principle of the genetic paradigm used to follow the *Prx1-cre* lineage, using the *R26^Lox-STOP-Lox-YFF^* reporter line combined with *Prx1-cre.* In tissues where cre is not expressed, YFP expression is prevented by the STOP cassette. In CRE-expressing mesenchymal cells, STOP cassette excision allows YFP expression. Bottom: scheme of a cross section of an *Prx1-cre*; *R26^YFP^; Gdnf^LacZ/+^* embryo, highlighting in green the cells in which YFP expression is activated, in red, the cells expressing *Gdnf^LacZ^,* and in white or grey, the other non-recombined tissues. **(B, C)** Cross-sections of E12.5 *Prx1-cre; Fatl^Flox/+^; Gdnf^LacZ/+^*; *R26^YFP/+^* **(B)** *and Prx1-cre; Fatl^Flox/+^; Gdnf^LacZ/+^*; *R26^YFP/+^* **(C)** embryos, stained with antibodies against GFP (YFP; to reveal the domain of *Prx1-cre* activity, in green), with and anti-β-galactosidase antibody (for *Gdnf^LacZ^,* red), visualized at four successive rostrocaudal positions spanning from the brachial plexus to the caudal half of the CM muscle. For each level, the inserts below represent a high magnification view of the area indicated in the yellow dotted boxes, showing red only, green only, and overlay. **(D)** Visual summary of the two components of *Gdnf^LacZ^* expression domain, spanning the sections shown in **(B)** and **(C)**: At the plexus level *Gdnf^LacZ^* is expressed in YFP+cells derived from *Prx1-cre* mesenchyme; whereas in the CM and LD muscles (emerging from the plexus and extending dorsally and caudally), *Gdnf^LacZ^*-positive cells do not express YFP as they are from the myogenic rather than from the mesenchymal lineage. At the point of emergence of the plexus myogenic patches (red only, yellow dotted line in **(B)**, second section) can be surrounded by mesenchymal-Gdnf cells (red+green=yellow, white dotted lines). The overall analysis shows that whereas *Prx1-cre-*mediated Fat1 ablation does not affect *Gdnf* expression in the plexus mesenchyme, but causes non-cell-autonomous reduction in the myogenic component of *Gdnf* expression domain, through a reduction of the number of *Gdnf^LacZ^*-expressing myogenic progenitors. Scale bars: **(B, C)** low magnification: 200 μm; inserts: 20 μm;

Among the two sub-domains of *Gdnf^LacZ^* expression, only the myogenic component was affected in *Prx1-cre; Fat1^Flox/Flox^; Gdnf^LacZ/+^; R26^YFP/+^* embryos (Fig 8C), whereas the mesenchymal subdomain of *Gdnf^LacZ^* expression appeared unaffected, both anatomically and in β-galactosidase intensity. In this mutant context, *Fat1* activity is disrupted in the YFP-positive cells, and not in the *Gdnf^LacZ^*/Pax7^+^ expressing progenitors. Thus, mesenchymal *Fat1* depletion has no effect on mesenchymal *Gdnf* expression at plexus level, in contrast with the constitutive knockout (S2 Fig). This indicates that this part of the knockout phenotype did not result from depletion of mesenchymal *Fat1*, and most likely reflects *Fat1* activity in another cell type. In contrast, *Fat1* ablation in the *Prx1-cre* lineage has a drastic impact on the myogenic component of *Gdnf^LacZ^* domain, not derived from the *Prx1-cre* lineage. This demonstrates that *Fat1* acts in a non-cell-autonomous manner. It is required in YFP-positive cells of the mesenchymal lineage to promote expansion of the sheet of migrating *Gdnf^LacZ^*/Pax7^+^ progenitors, possibly by modulating the production by mesenchymal cells of signals controlling progenitor pool expansion and/or migration. Interestingly, at E12.5, the CM and LD muscles are almost entirely surrounded by *Prx1-cre*-derived mesenchymal cells, with exception of the dorsal-most tip of the CM, which lies beyond the dorso-ventral limit of lateral-plate mesoderm derived mesenchyme [73] (Fig 7E and Fig 8). Interestingly, dorsal to this dorso-ventral limit, thickness of the CM appears reinforced; suggesting that once the myoblasts reach the non-recombined mesenchymal zone, the unaltered *Fat1* activity available in this dorsal environment allows them to resume their normal growth behavior, providing a possible explanation for the apparent sparing of dorsal CM at E14.5 (Fig 5C). Altogether, these data support a model in which muscle-associated mesenchymal cells exert a *Fat1*-dependent positive influence on CM muscle growth/extension.

### *Fat1* activity in the Pdgfrα-expressing connective tissue lineage

Thus far, we have identified the *Prx1-cre* lineage as the cell type in which *Fat1* is required to control CM expansion and scapulo-humeral muscle patterning. However, the *Prx1-cre*-derived lineage is broad (Fig 8) and includes several distinct subtypes of connective tissues (CT). These comprise specialized CT such as bones and cartilage, dense regular CT such as tendons, and dense irregular CT (here referred to as loose CT) such as muscle-resident mesenchymal derivatives or the peri-muscular environment [13]. All of these subtypes express *Fat1* at relatively high levels (Fig 4, S3 Fig, S4 Fig). The CM grows towards a subcutaneous layer of connective tissue, in which we observed increasing levels of *Fat1* expression. Furthermore, CM extension appears to be affected on the side of its growth towards the skin interface. Altogether, this suggests that this subcutaneous interface with the CM muscle might be where this *Fat1* function is taking place. Whole-mount *in situ* hybridization with the tenocyte/tendon marker *Scleraxis* highlights sites of intense expression in the limb tendons, or at the interface between intercostal muscles and ribs (visible by transparency, beneath the CM), but only shows background *Scleraxis* levels in the region in which CM is migrating (S5A Fig). When analyzing expression of other markers of CT subtypes by immunohistochemistry on embryo sections, we found that this subcutaneous CT expresses high levels of Pdgfrα and TenascinC, but not Tcf4 (also known as Tcf7L2) (S10 Fig), which was otherwise detected at other muscle extremities and in subsets of limb muscle progenitors (S10 Fig), as previously reported [77,78].

We next asked whether restricting *Fat1* ablation to this subtype of mesenchyme could be sufficient to interfere with CM spreading and differentiation. Our characterization prompted us to consider inactivating *Fat1* in the *Pdgfrα*-expressing lineage. However, as *Pdgfrα* is expressed at early stages in cells from which a large lineage will derive, we chose to use an inducible *Pdgfrα-cre/ERT2* transgenic line (here referred to as *Pdgfrα-iCre),* in which CRE is expressed in the *Pdgfrα* domain, but remains catalytically inactive unless tamoxifen is provided [79]. This strategy is expected to highlight only a subset of mesenchymal cells expressing Pdgfrα at the time of Tamoxifen administration, including the loose connective tissue at the skin-muscle interface (Fig 9A, B). To establish the conditions to obtain reliable excision rate, we first combined the *Pdgfrα-iCre* line to the R26-YFP reporter [66], and exposed pregnant females to tamoxifen treatment. Optimal excision efficacy was obtained when injecting a first dose of 50mg/kg at E9.75 (earlier injections frequently lead to developmental arrest), followed by a second injection of 100mg/kg at E10.5 (Fig 9A, B). This allowed consistent detection of the R26-YFP signal when screening whole embryos just after dissection (S10B Fig). Analysis of YFP expression by IHC on sections of *Pdgfrα-iCre; R26^YFP/+^* embryos confirmed that recombined YFP-positive cells *(“PdgJm-iYFP”* cells) were mostly localized in the loose connective tissue surrounding the CM, whereas none of the Myh1-positive fibers exhibited any detectable recombination (Fig 9A). Although our experimental conditions do not allow simultaneous detection of YFP and Pax7 (the heat-induced-epitope-recovery treatment required to detect Pax7 protein abrogates detection of YFP), analysis of neighboring sections was consistent with the Pax7-containing area not exhibiting any YFP activity (Fig 9A). Thus, these experimental conditions allow an approximate excision rate of 30% in the loose subcutaneous connective tissue surrounding the CM, whereas no activity in the myogenic lineage was detected.

**Fig 9:**
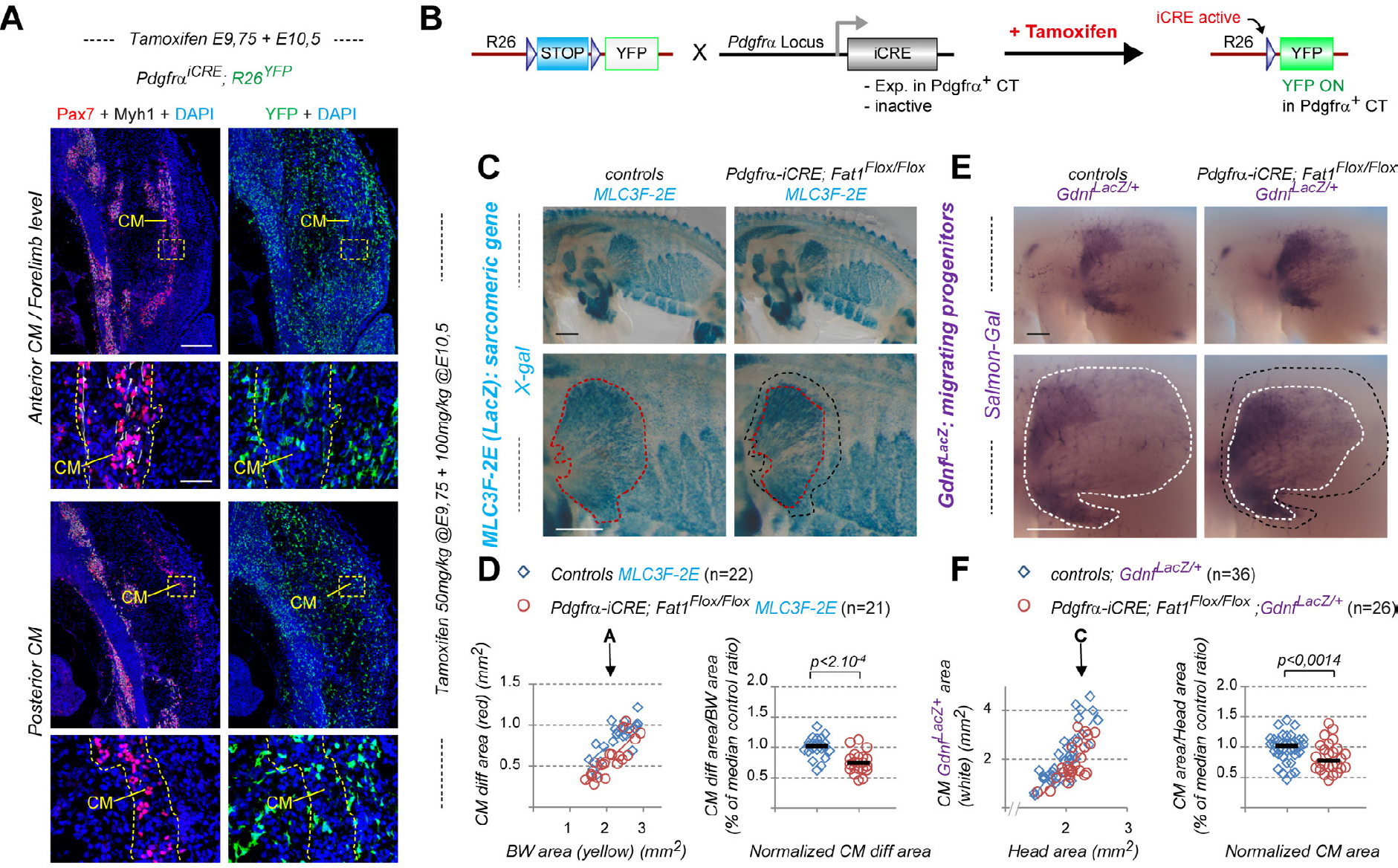
Inducible *Fat1* deletion in the pdgfrα connective tissue lineage alters progression of CM migration and differentiation. **(A)** Cross-sections of a *Pdgfrα-iCre; R26^YFP/+^* embryo collected at E12.5, after in utero administration of Tamoxifen at E9.75 (50mg/kg) + E10.5 (100mg/kg). Alternate sections at anterior and posterior CM levels, respectively, were immunostained with antibodies against: Left panels: Pax7 (red) and myh1 (white), plus DAPI (blue); Right panels: GFP (green), plus DAPI (blue), to reveal the outcome of Pdgfrα-iCre-mediated R26-YFP recombination (Right panels). The yellow dotted boxes indicate the areas magnified in the bottom panels. **(B)** Principle of the genetic paradigm used to follow the *Pdgfrα-iCre* lineage, using the *R26^Lox-STOP-Lox-YFP^* reporter line combined with *Pdgfrα-iCre.* iCRE (CRE/ERT2) is expressed in the domain of *Pdgfrα* expression, but remains catalytically inactive. iCRE activity is permitted by in utero treatment with Tamoxifen. Catalytic activity is triggered in the cells expressing iCRE at the time of Tamoxifen treatment, thus allowing the stop cassette to be deleted, and YFP to be permanently expressed. **(C, E)** Whole-mount β-galactosidase staining was performed using X-gal as substrate on embryos carrying the *MLC3F-2E* transgene **(C)**, or using Salmon-Gal as substrate on embryos carrying the *Gdnf^LacZ/+^* allele **(E)**, in the context of Tamoxifen-induced *Fat1* deletion in the *Pdgfrα* lineage driven by *Pdgfrα-iCre* at E12.5. Top images show a side view of the whole flank of an embryo. Lower images are higher magnification of the area in which the CM spreads. Red- and white-dotted lines correspond to the areas covered in control embryos by *MLC3F-2E^+^* CM-fibers **(C)**, and by *Gdnf^LacZ+^* progenitors **(E)**, respectively. In comparison, the corresponding areas observed in mutants are indicated as black-dotted lines in both cases. **(D)** Quantification of the expansion rate of differentiated CM fibers. Left graph: For each embryo side, the area covered by differentiated CM fibers was plotted relative to the area occupied by body wall muscles. Right plot: for each embryo, the CM area/Body wall area was normalized to the median ratio of control embryos. Blue dots: *Fat1^Flox/Flox^; MLC3F-2E* (n=22, control embryos from Tamoxifen-treated litters); Red dots: *Pdgfrα-iCre; Fatl^Flox/Flox-^; MLC3F-2E* (n=21). Underlying data are provided in S1 Data. **(F)** Quantification of the expansion rate of the area occupied by *Gdnf^LacZ+^* progenitors. Left plot: For each embryo side, the area covered by *Gdnf^LacZ+^* progenitors was plotted relative to the trunk length. Right plot: for each embryo, the *Gdnf^LacZ+^* CM area/Trunk Length was normalized to the median ratio of control embryos. Blue dots: *Fat1^Flox/Flox^; Gdnf^LacZ/+^* (n=36; control embryos from Tamoxifen-treated litters); Red dots: *Pdgfrα-iCre; Fatl^Flox/Flox^; Gdnf^LacZ/+^* (n=26). Underlying data are provided in S1 Data. Scale bars: **(A)** low magnification: 200 μm; high magnification: 50 μm; **(C, E)** 500 μm.

We next asked whether *Fat1* ablation in around 30% of the loose CT surrounding the CM was sufficient to interfere with its expansion and with progression of differentiation. The *Pdgfrα-iCre* line was combined to the *Fat1^Flox^* allele, and with either the *MLC3F-2E* transgenic line to follow muscle differentiation, or with *Gdnf^LacZ^* to follow the progression of progenitor migration (Fig 9C, E). Pregnant females were treated with Tamoxifen as defined above, and embryos collected at E12.5. In all cases, mutants were compared to control embryos from Tamoxifen-treated litters. Analysis was performed as previously, by measuring the area occupied by the *MLC3F-2E^+^* fibers in the CM, as compared to the area occupied by body wall muscles, or by assessing the area occupied by the *Gdnf^LacZ^*-expressing progenitors as compared to the head area and/or trunk length. *Fat1* ablation driven by such restricted recombination paradigm leads to a significant delay in the progression of both CM muscle fiber elongation (Fig 9C, D), and CM progenitor migration (Fig 9E, F). The median differentiated CM area measured in Tamoxifen-treated *Pdgfrα-iCre; Fat1^Flox/Flox^; MLC3F-2E* embryos (ratio of CM differentiated area/body wall area, normalized to median control ratio) was reduced by approximately 30% compared to Tamoxifen-treated control embryos (Fig 9C, D). Similarly, we observed a 30% reduction of the median area covered by CM progenitors observed in Tamoxifen-treated *Pdgfrα-iCre; Fat1^Flox/Flox^; Gdnf^LacZ/+^* embryos (Fig 9E, F). In contrast, there was no apparent effect on the shape of scapulohumeral muscles, and we didn’t detect any significant enhancement of myocyte dispersion in the forelimb (S10C Fig, S10D Fig). In the conditions of Tamoxifen treatment we used for the *Pdgfrα-iCre* line, the percentage of cells in which the reporter R26-YFP expression indicates cre-mediated recombination (Fig 9A) is much more restricted compared to the *Prx1-cre* lineage (Fig 7E, Fig 8). This indicates that *Fat1* activity is required in a significant proportion of cells among these recombined *“Pdgfrα-iYFP”* cells for CM muscle spreading. Overall, these data are consistent with *Fat1* being required in the loose connective tissue for its non-cell-autonomous influence on CM progenitor spreading and muscle fiber extension. In conclusion, we have identified the mesenchymal lineage as the place where *Fat1* activity is required to promote CM muscle expansion. We have refined our knowledge on this lineage by uncovering that for a large part, this function occurs in the *Pdgfrα-*dependent loose connective tissue, such as the subcutaneous layer in which CM expansion occurs. Finally, we find that this lineage represents a subset of the *Pdgfrα-iCre* lineage, and corresponds to the subset of cells expressing *Pdgfrα* between E9.5 and E10.5.

### *Fat1* ablation in the mesenchyme alters the pattern of axon innervating the CM

Having refined the knowledge on which connective tissue type is required for the regulation of CM muscle expansion by *Fat1*, we next asked which cell type was accountable for the alteration of the CM innervation pattern observed in constitutive knockouts. We asked whether this activity of *Fat1* in the mesenchyme-derived connective tissue was sufficient to explain alterations of CM innervation and changes in MN pool specification, or whether *Fat1* is also additionally required in MNs. We first asked which site of *Fat1* expression was accountable for the altered pattern of axonal arborization of the target muscle CM seen in constitutive *Fat1* knockouts. We therefore performed whole-mount anti-neurofilament immunohistochemistry on embryos lacking *Fat1* either in MNs (*Olig2^cre/+^*; *Fat1^Floxl/Flox^)* or in mesenchyme (*Prx1-cre*; *Fat1^Flox/Flox^*) and examined the pattern of CM motor innervation. After imaging of BB-BA cleared embryo flanks (Fig 10A, upper images), the thoracic cage, and associated thoracic nerves, was removed to allow easier visualization of CM-innervating axons (Fig 10A, lower panels).

**Fig 10:**
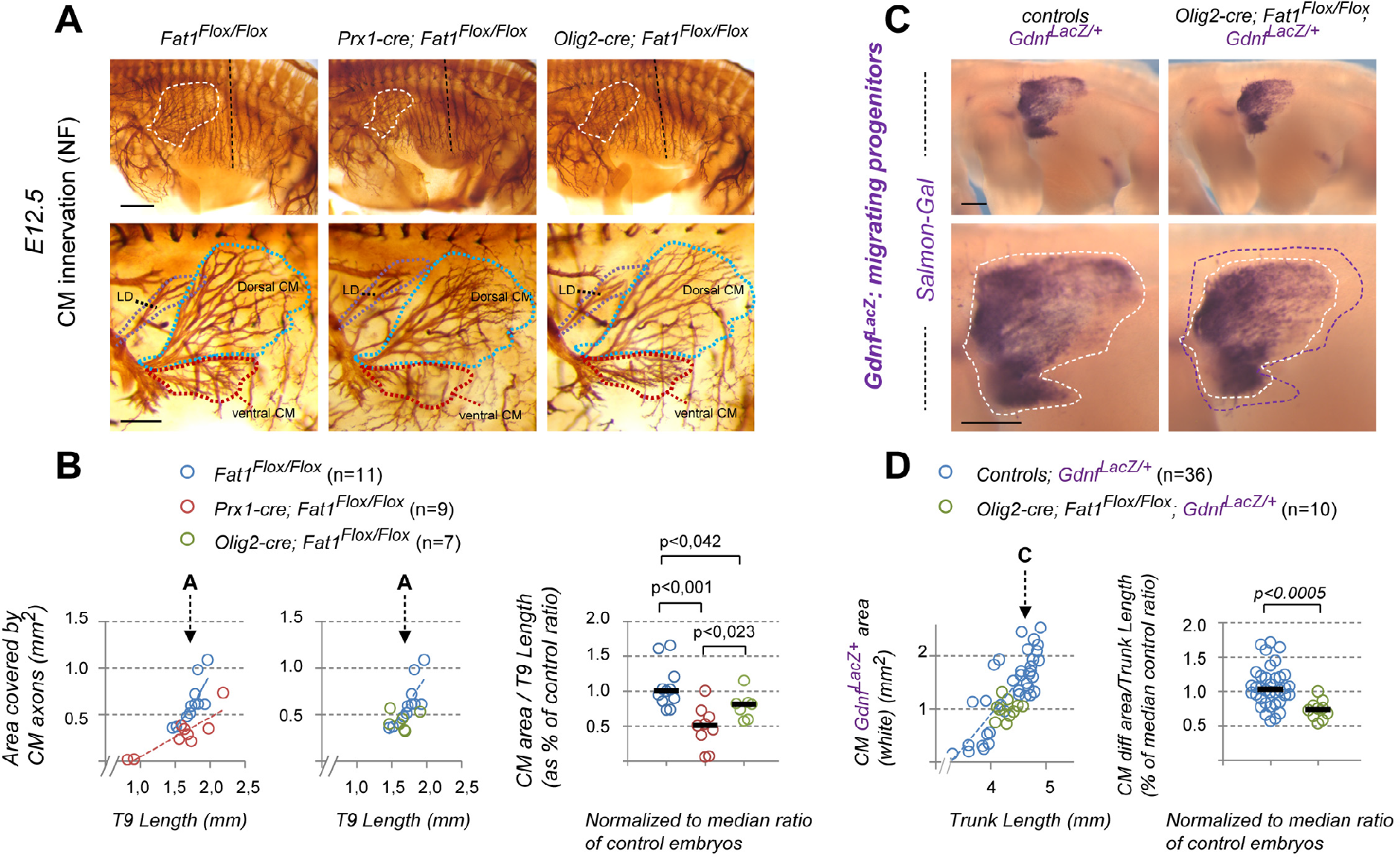
Dual control of CM innervation by Fat1 activity in MNs and mesenchyme. **(A)** Anti-Neurofilament immunohistochemistry was performed on E12.5 embryos in the context of *Prx1-cre*-mediated mesenchyme deletion of Fat1 or of *Olig2-cre*-mediated Fat1 deletion in Motor neurons. After BA-BB clearing, the embryos were cut in half and the internal organs and skin removed to visualize the CM and brachial plexus. Upper panels show the entire flank of *Fatl^Flox/Flox^* (left), *Prxl-cre; Fatl^Flox/Flox^* (middle), or *Olig2-cre; Fatl^Flox/Flox^* (right) at comparable stages. The area covered by CM-innervating axons is outlined with white-dotted lines. Lower panels represent a higher magnification of the flank area containing the CM and corresponding motor axons, after manual removal by dissection of a large part of the thoracic cage and nerves. The shape of the control area corresponding to dorsal CM (dCM) and ventral CM (vCM) is outlined in green- and red-dotted lines, respectively. **(B)** quantifications of the progression of CM innervation. Left and Middle plots: For each embryo side, the area covered by CM-innervating axons was plotted relative to the length of the T9 thoracic nerve, comparing *Prxl-cre; Fatl^Flox/Flox^* embryos (red dots) to controls (blue dots) on the left plot, and *Olig2-cre; Fatl^Flox/Flox^* embryos (green dots) to the same set of controls (blue dots) on the middle plot. Arrows point to the stage of representative examples shown in **(A)**. Right plot: for each embryo, the CM-innervated area/T9 Length was normalized to the median ratio of control embryos, by size range. Blue dots: Controls (n=11, same sample set in left and middle plot, pooling respective littermates); Red dots: *Prx1-cre; Fatl^Flox/Flox^* (n=9); Green dots: *Olig2-cre; Fatl^Flox/Flox^* (n=7). Underlying data are provided in S1 Data. (C) Whole-mount β-galactosidase staining was performed using Salmon-Gal as substrate on embryos carrying the *Gdnf^LacZ/+^* allele, in the context of MN-specific deletion of *Fat1,* driven by *Olig2-cre* at E12.5. Top images side view of the whole flank. Lower images: higher magnification of the area in which the CM spreads. White dotted lines correspond to the area covered by *Gdnf^LacZ+^* progenitors, whereas the purple line on the *Olig2-cre; Fatl^Flox/Flox^* image indicates the shape of the control area to highlight the difference. **(D)** Quantification of the expansion rate of the area occupied by *Gdnf^LacZ+^* progenitors. Left plot: For each embryo side, the area covered by *Gdnf^LacZ+^* progenitors was plotted relative to the trunk length. Right plot: for each embryo, the *Gdnf^LacZ+^*CM area/Trunk Length was normalized to the median ratio of control embryos. Blue dots: *Fat1^Flox/Flox^; Gdnf^LacZ/+^* (n=36; same control set as in Fig 7D); Red dots: *Olig2-cre; Fatl^Flox/Flox-^; Gdnf^LacZ/+^* (n=10). Underlying data are provided in S1 Data. Scale bars: (A) top: 500 μm; (A) bottom: 200 μm; (B) 500 μm.

We found that mesenchyme-specific mutants (*Prx1-cre*; *Fat1^Flox/Flox^* embryos) exhibited an overall shortening of the area covered by CM-innervating axons compared to stage-matched controls. This effect was very severe ventrally, with a complete absence of motor axon extending in the ventral part of the CM (vCM, Fig 10A), whereas axons extending in the dorsal CM (dCM) were present but shorter than in control embryos. To quantify this effect, the area covered by CM-innervating axons was plotted relative to a landmark indicating developmental age, using the length of the T9 nerve (Fig 10B). The net effect quantified by following the ratio area of CM axons/T9 length was almost as severe as the effect measured in constitutive knockouts (Fig 2). This effect was strikingly reminiscent of the profound shortening and shape change of the area occupied by *Gdnf^LacZ+^* CM progenitors seen earlier (Fig 7). Thus, *Fat1* ablation in the mesenchyme non-cell-autonomously influences not only expansion of the muscle progenitor area, but also extension of motor axons projecting in this muscle. This indicates that even in this context of severe perturbation of CM expansion, there remains a strong coupling between motor axons and muscle progenitors, and the topography of muscle fiber extension. To determine whether growth of CM-innervating axon was dependent on muscle, genetic depletion of the myogenic lineage was achieved through *Myf5-cre*-driven [80] conditional expression of diphtheria toxin A (DTA) regulated by the *R26* locus *(R26^Lox-LacZ-STOP-Lox-DTA^* locus, [81]). This paradigm resulted in a severe loss of muscle mass, as assessed with the *MLC3F-2E-LacZ* reporter. In the trunk, X-gal staining shows the loss of myogenic cells corresponding to the CM muscle, to intercostal muscles, and to the trapezius muscle, whereas epaxial muscles, although irregular, were less efficiently deleted (S11B Fig). As previously shown [46,82], limb innervating nerves were present and had extended in spite of the absence of muscles (S11B Fig). In contrast, the thoracic intercostal nerves, the CM-innervating motor nerve, and the spinal accessory nerve (or nerve XI), which innervates the trapezius muscle, were severely affected by muscle depletion (S11A Fig, S11C Fig) compared to stage matched control embryos. These results indicate that genetic suppression of myogenic cells was sufficient to cause a drastic shortening of CM-innervating axons, in contrast to the lack of effect on the trajectory of limb innervating nerves (S11 Fig). This supports the idea that the coupling between motor and muscle phenotypes observed in *Prx1-cre*; *Fat1^Flox/Flox^* embryos could be mediated by muscle-derived factors.

Although the effect of MN-specific *Fat1* ablation appeared less obvious, our method of quantification nevertheless uncovered a mild but significant reduction (around 20%) in the rate of expansion of the area covered by CM-innervating axons in *Olig2^cre/+^; Fat1^Flox/Flox^* embryos compared to age-matched controls (Fig 10A,B). Overall the shape of the innervated area does not change qualitatively, but axons are shorter in *Olig2^cre/+^; Fat1^Flox/Flox^* embryos than in age-matched controls. This indicates that in absence of *Fat1*, CM motor axons extend at a slower rate. Even though the net effect was small compared to the phenotype of constitutive knockouts, this result supports the notion that *Fat1* is required cell-autonomously for CM motor axon growth. Since CM motor axons progress hand in hand with Pax7^+^/*Gdnf^LacZ+^* progenitors, ahead of muscle fiber extension (Fig 3), we next asked if this mild but significant shortening of axons had any impact on the expansion of the CM progenitor area. We therefore brought the *Gdnf^LacZ^* allele in the context of MN-specific *Fat1* mutants, and followed the area covered by *Gdnf^LacZ^* progenitors (Fig 10C,D). Unexpectedly, whereas the shape of the *Gdnf^LacZ+^* area appeared qualitatively unchanged, area measurements uncovered a significant lowering (of about 25%) of the ratio *Gdnf^LacZ^* area / trunk length in *Olig2-cre; Fat1^Flox/Flox^; Gdnf^LacZ/+^* embryos, compared to *Fat1^Flox/Flox^; Gdnf^LacZ/+^* controls. This result uncovers that *Fat1* ablation in motor neurons non-cell-autonomously interferes with expansion of CM progenitors. This effect may either reflect a direct control by *Fat1* in motor neurons of the process of muscle progenitor spreading, or a *Fat1*-independent consequence of the slower progression of motor axons. This finding is counter-intuitive, as we had previously excluded that MN-specific *Fat1* loss had any impact of muscle fiber extension (S8 Fig, S9 Fig). Thus, the shortening of the area covered by *Gdnf^LacZ^* progenitors is not associated with a reduced progression of myogenic differentiation in the CM. This indicates that the slower migration seen in *Olig2-cre; Fat1^Flox/Flox^; Gdnf^LacZ/+^* embryos does not significantly influence efficiency of muscle fiber extension.

Thus, the alterations in CM innervation pattern observed in constitutive *Fat1* knockouts appear to result in part from removal of *Fat1* axon growth promoting activity in motor neurons, and to a large extent from the non-cell-autonomous consequences of Fat1 ablation in the mesenchyme, which simultaneously affects expansion of the CM progenitor area and axonal growth. Although these two non-cell autonomous consequences of mesenchymal Fat1 ablation could represent two independent phenotypes, the fact that axonal extension of CM-innervating axons requires the presence of myogenic cells supports the possibility that the two phenotypes might be causally linked. Given its known axonal growth promoting activity, GDNF is a likely candidate to explain how impaired myogenic progenitor expansion could lead to inhibited axon elongation in *Prx1-cre; Fat1^Flox/Flox^* embryos. In contrast, the observation of a non-cell-autonomous impact of reduced axonal elongation on progenitor expansion in *Olig2-cre; Fat1^Flox/Flox^; Gdnf^LacZ/+^* embryos indicate that motor axons also play a role in promoting expansion of the CM progenitor domain.

### Mesenchyme and MN activities of *Fat1* are required for MN fate acquisition

We next asked which tissue-specific *Fat1* mutant was most closely reproducing the effects on expression of *Runx1* and *Clusterin* in CM motor pools observed in constitutive knockouts (comparing *Prx1-cre* and *Olig2-cre).* We found that expression of both genes in the CM motor pools was significantly reduced not only in mesenchyme-specific *Prx1-cre; Fat1^Flox/Flox^* mutants, but also in MN-specific *Fat1* mutants (*Olig2^cre/+^; Fat1^Flox/Flox^),* (Fig 11A-D). These observations demonstrate that *Fat1* simultaneously exerts two complementary functions in MNs and in the mesenchyme, each of which cooperatively contributing to the acquisition of complete CM motor pool identity. Thus, the constitutive knockout phenotype results from the cumulated consequences of abrogating *Fat1* functions in both MNs and mesenchyme. Interestingly, the reduction of *Runx1/Clusterin* expression in *Fat1* conditionals is partially reminiscent of the phenotypes of *Etv4* (Fig 6E-J) and of *Gdnf* mutants (Fig 11A-D). Furthermore the degree of reduction in *Runx1* and *Clusterin* expression correlates with the degree of reduction of the area covered by *Gdnf* expressing myogenic progenitors observed in *Prx1-cre; Fat1^Flox/Flox^* and in *Olig2^cre/+^; Fat1^Flox/Flox^* embryos, respectively (Fig 7 and Fig 10). This suggests that the lowering of *Runx1* and *Clusterin* expression could be a consequence of the lowering in *Gdnf* levels, and of the subsequent changes in *Etv4* expression, even though subtle (Fig 6A-C). However, although comparable, the MN pool specification phenotype of either conditionals or constitutive *Fat1* mutants is less severe than the effect of complete *Gdnf* elimination. In spite of the reduced expression of the myogenic component of *Gdnf* expression domain in *Prx1-cre; Fat1^Flox/Flox^* mutants, the overall GDNF level is maintained owing to the remaining mesenchymal-*Gdnf* expression at plexus levels (Fig 8). This minimizes the impact on induction of Etv4, the transcription factor acting upstream of *Runx1* and *Clusterin* (Fig 6). As a result, *Etv4* expression is less severely affected in *Prx1-cre; Fat1^Flox/Flox^* embryos than in embryos with only one functional copy of *Gdnf* (compare *Fat1^Flox/Flox^; Gdnf^LacZ/+^* and *Fat1^Flox/Flox^* (Fig 11E, F)). As a consequence, the residual expression of *Etv4* is sufficient to ensure correct mediolateral positioning of MNs in the spinal cord of *Prx1-cre; Fat1^Flox/Flox^* embryos (S12A Fig), contrasting with what occurs in *Gdnf^-/-^* or *Etv4^-/-^* spinal cords (S12A Fig, [39,41]). However, genetic lowering of *Gdnf* exacerbates the effect of mesenchyme-specific *Fat1* ablation on *Etv4* expression, when both conditions are combined (Fig 11E, F). Consequently, *Etv4* expression is significantly lower in *Prx1-cre; Fat1^Flox/Flox^; Gdnf^LacZ/+^* embryos than in either *Fat1^Flox/Flox^; Gdnf^LacZ/+^* or *Prx1^cre/+^; Fat1^Flox/Flox^* embryos (Fig 11E, F). The possibility that *Gdnf* could itself be involved in regulating the subcutaneous progression and number of CM progenitors (thereby indirectly influencing the amount of other factors secreted by CM progenitors) is ruled out by the observation of normal expansion of the area covered by *Gdnf^LacZ^* expressing CM progenitors in *Gdnf^LacZ/LacZ^* embryos, compared to *Gdnf^LacZ/+^* (S12B Fig). Altogether, these data identify *Gdnf* as an essential mediator of the effect of mesenchyme-specific *Fat1* ablation on acquisition of CM motor pool fate, part of which involves the dose-dependent fine tuning of *Etv4* expression.

**Fig 11:**
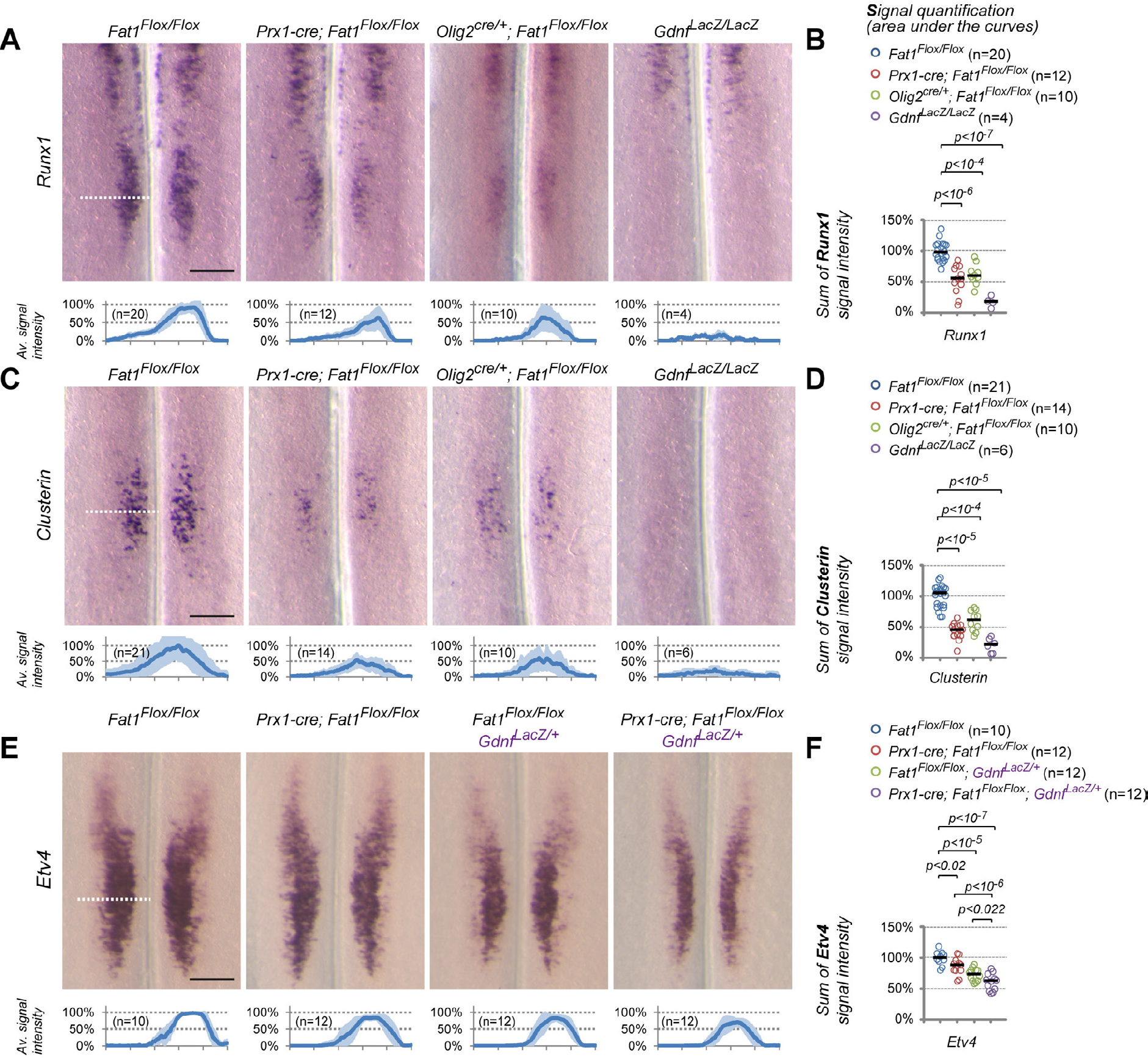
*Fat1* is required in both peripheral mesenchyme and motor neurons for proper specification of CM motor pool identity. **(A-B)** Analysis of *Runxl* **(A,B)** and *Clusterin* **(C,D)** mRNA expression in brachial spinal cords from E12.5 *Fatl^Flox/Flox^, Prxl-cre; Fatl^Flox/Flox^, Olig2^cre/+^; Fatl^Flox/Flox^* and *Gdnf^LacZ/LacZ^* embryos. Spinal cords are presented as open-book, flat-mounted, so that the line in the middle of each picture corresponds to the ventral midline, and the motor columns are visible on each side. Fat1 ablation in both the mesenchyme and MNs lead to significant lowering of *Clusterin* and *Runxl* expression levels, whereas expression is absent in the *Gdnf^LacZ/LacZ^* embryos. Quantifications in **(A-D)**: each plot in **(A)**, and **(C)** represents respectively, the average *Runxl* and *Clusterin* signal distribution (± standard deviation in light blue) measured on the indicated number of spinal cord sides along the white dotted line in each image above. Underlying data are provided in S1 Data. **(B)** The cumulated *Runxl* signal intensity (area under the curves in **(A)** was plotted for each spinal cord side. *Fatl^Flox/Flox^* (n=20) *Prxl-cre; Fatl^Flox/Flox^* (n=12), *Olig2^cre/+^; Fatl^Flox/Flox^* (n=10) and *Gdnf^LacZ/LacZ^* (n=4). Underlying data are provided in S1 Data. **(D)** The cumulated *Clusterin* signal intensity (area under the curves in **(C)** was plotted for each spinal cord side. *Fatl^Flox/Flox^* (n=21) *Prxl-cre; Fatl^Flox/Flox^* (n=14), *Olig2^cre/+^; Fatl^Flox/Flox^* (n=10) and *Gdnf^LacZ/LacZ^* (n=6). Underlying data are provided in S1 Data. **(E)** In situ hybridization analysis of the *Etv4* expression domain in brachial spinal cord of flat-mounted E12.5 *Fatl^Flox/Flox^, Prxl-cre; Fatl^Flox/Flox^, Fatl^Flox/Flox^; Gdnf^LacZ/+^* and *Prxl-cre; Fatl^Flox/Flox^; Gdnf^LacZ/+^* embryos at E12.5. Whereas *Fatl* ablation in mesenchyme mildly affects *Etv4* expression, genetic lowering of *Gdnf* levels further prevents *Etv4* induction in *Prxl-cre; Fatl^Flox/Flox^; Gdnf^LacZ/+^* embryos, when compared to either *Prxl-cre; Fatl^Flox/Flox^* or *Fatl^Flox/Flox^; Gdnf^LacZ/+^.* **(E,F)** Quantifications and statistical analyses of the sum of signal intensity corresponding to the area under the curves in plots shown in **(E)**: Each dot represents the sum of intensity for one spinal cord side. *Fatl^Flox/Flox^* (n=10) *Prxl-cre; Fatl^Flox/Flox^* (n=12), *Fatl^Flox/Flox^;Gdnf^LacZ/+^* (n=12) and *Prxl-cre; Fatl^Flox/Flox^; Gdnf^LacZ/+^* (n=12)). Underlying data are provided in S1 Data. Scale bars: (A, C, E): 200 μm.

### Impact of MN-specific and Mesenchyme-specific *Fat1* depletion on adult muscle phenotypes

Finally, we sought to evaluate the respective functional relevance of the two sites of *Fat1* expression for NMJ integrity in the adult CM muscle in both MN-specific and mesenchyme-specific mutants. We focused on the dorsal CM, where in spite of delayed myogenesis in mesenchyme-specific mutants, muscle fibers and motor axon have extended. In this region, muscle fibers, visualized with F-actin staining with fluorescent phalloidin, have formed in both mutants (Fig 12A), and have been innervated by motor axons, leading to a normal intramuscular pattern of NMJ distribution in both MN-specific and mesenchyme-specific *Fat1* mutants. We analyzed synapse morphology in NMJ-enriched regions of the dCM, by labeling the post-synaptic site with alpha-bungarotoxin (BTX, which binds Acetylcholine receptors; AchR), and the nerve endings with anti-neurofilament antibodies. As a measurable feature of NMJ integrity, we analyzed the surface area of AchR-rich zones for an average of 23 synapses per mouse, comparing *Olig2^cre/+^*; *Fat1^Flox/Flox^* and *Prx1^cre/+^; Fat1^Flox/Flox^* mice with *Fat1^Flox/Flox^* mice, aged 9-13 months. This analysis uncovered that synapses are reduced in size in both genotypes compared to controls, this effect being more pronounced in *Prx1^cre/+^; Fat1^Flox/Flox^* mice than in *Olig2^cre/+^; Fat1^Flox/Flox^,* with a median synapse area of 420 μm^2^ (81%) and 220 μm^2^ (42%), respectively, compared to 512 μm^2^ (100%) in controls (Fig 12B). A careful analysis of the distribution of synapse areas in each genotype confirmed a significant lowering in the percentage of large synapses, and an increase in the percentage of smaller synapses, with up to 68% of synapses in *Prx1^cre/+^; Fat1^Flox/Flox^* mice, compared to 32% in *O1ig2^cre/+^; Fat1^Flox/Flox^* and 12% only in *Fat1^Flox/Flox^* mice, were smaller than 300 μm^2^ (Fig 12B). Analysis on the same sections/samples of the distribution of fiber width revealed no significant change in fiber diameter, arguing that the reduced size does not simply reflect adaptation to a change of the muscle fibers themselves (Fig 12C). Instead, this smaller synapse area is indicative of altered NMJ integrity, and was accompanied by spots of local denervation of the synapses, or by fragmentation of the AchR-rich synaptic area (Fig 12A).

**Fig 12:**
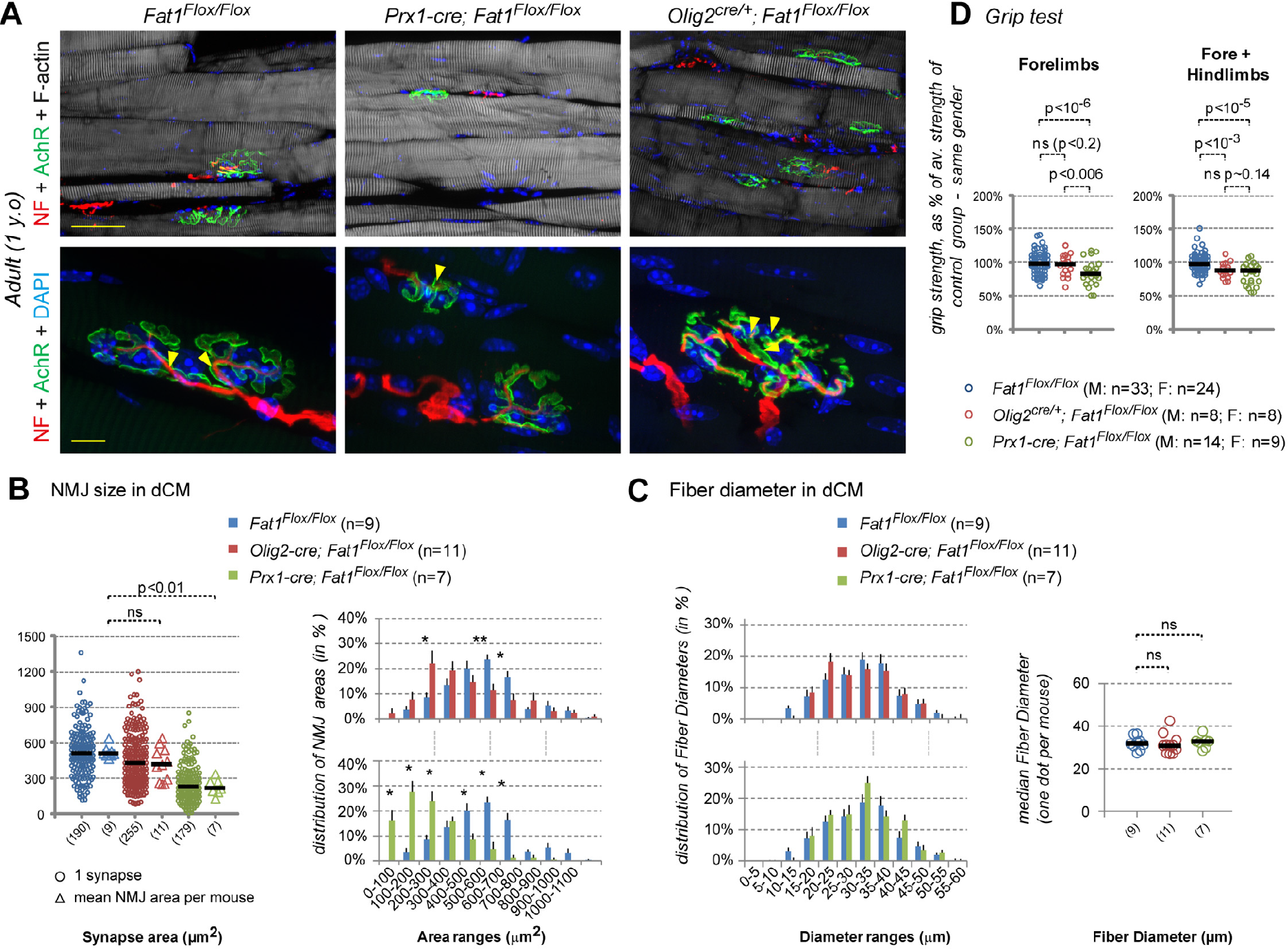
Impact of mesenchyme-specific and Motor Neuron-specific Fatl ablation on adult CM muscle anatomy and grip strength. **(A-C)** Analysis of NMJ morphology **(A)** was performed by immunostaining with α-Bungarotoxin (green, detecting AchR), phalloidin (white, detecting F-actin), and anti-Neurofilament antibodies on cryosections of the dorsal CM, in *Fatl^Flox/Flox^,* in *Olig2^cre/+^; Fatl^Flox/Flox^* and *Prxl-cre; Fatl^Flox/Flox^* adult mice (~1 year old). **(A)** Top pictures show low 20x magnifications, whereas bottom pictures show high magnification of individual synapses. **(B, C)** Histograms showing quantification of NMJ area **(B)**, and Fiber diameter **(C)** distributions in the dorsal CM muscle of *Fatl^Flox/Flox^* (n=9), *Olig2^cre/+^; Fatl^Flox/Flox^* (n=11) and *Prxl-cre; Fatl^Flox/Flox^* (n=7) mice. Underlying data are provided in S1 Data. **(B)** Left plot shows the synapse area, with for each genotype all synapses plotted (circles), as well as the median NMJ area per mouse (with each triangle representing one mouse). An average of 20 synapses per mouse was analyzed in the indicated number of mice. Total numbers of synapses and mice are indicated below the graph. Right plots: Average distribution of synapse areas in *Fatl^Flox/Flox^* (blue bars) and *Olig2-cre; Fatl^Flox/Flox^* (red bars) mice (top), and *Fatl^Flox/Flox^* (blue bars, same data as above) and *Prxl-cre; Fatl^Flox/Flox^* (green bars) mice (bottom). **(C)** Left plots: Average distribution of fiber diameters in the same samples. Right plots show the median fiber diameter in each mouse (one dot representing one mouse). Statistical significance: * indicates p<0.01, ** indicates p<0.001, Mann Whitney test. Underlying data are provided in S1 Data. **(D)** Measurements of grip strength in mice with the indicated genotypes. In the plots, each dot represents the average value for one mouse, assaying forelimb grip strength (left plot), or cumulative strength of forelimbs plus hindlimbs (right). For each mouse, the measured strength was normalized to the mean strength of the control group of the same gender, so that males and females could be expressed as percentage and pooled on the same graph. Underlying data are provided in S1 Data. Statistical significance: p values are indicated for the relevant comparisons, using unpaired student T-test for values with normal distribution and equal variance, or Mann Whitney test otherwise. Significance threshold: p<0.01. Scale bars: (A) upper images: 50 μm; (A) lower images: 10 μm.

We next thought to determine if there were consequences in terms of the force generating abilities of affected muscles. A typical assay is the grip test, which measures the grabbing force (also called prehension force) permitted by the coordinated activity of limb muscles (forelimbs and/or hindlimbs). Although phenotypes in the subcutaneous CM muscle are unlikely to contribute to forelimb movements, this assay allows detecting loss of strength in other limb muscles. The subtle defect in NMJ integrity observed in *Olig2^cre/+^; Fat1^Flox/Flox^* mice, is expected to be restricted to target muscles of *Fat1*-expressing MNs. These include the CM muscle, and a second pool of *Etv4*-negative neurons for which we did not identify the target muscle. Overall, this did not result in any significant loss of forelimb grip strength in *Olig2^cre/+^; Fat1^Flox/Flox^* mice compared to control mice (Fig 12C). Nevertheless, *Fat1* RNA, *Fat1^LacZ^* activity, and Fat1 protein were detected in subsets of lumbar MNs at E12.5 (Fig 1E) and E13.5 (S3B Fig). We therefore asked if *Fat1* ablation in these lumbar MNs had any impact in the force generation capacity of hindlimb muscles. A reliable way of approaching this force is to measure forelimb plus hindlimb strength (and inferring hindlimb strength by subtracting forelimb strength). Consistently, we observed a significant lowering of forelimb plus hindlimb grip strength in *Olig2^cre/+^; Fat1^Flox/Flox^* mice compared to *Fat1^Flox/Flox^* controls. This indicates that MN-*Fat1* is required for the functional integrity of subsets of lumbar MN pools innervating selected hindlimb muscles.

In contrast, mesenchyme-specific *Fat1* ablation in *Prx1^cre/+^; Fat1^Flox/Flox^* mice leads to significant lowering of both forelimb and the cumulated forelimb + hindlimb grip strength, with an effect more pronounced than in *Olig2^cre/+^; Fat1^Flox/Flox^* mice. This is consistent with the possibility that these mesenchyme-specific mice might exhibit myogenesis defects in muscles other than the CM and LD, including muscles in the scapulohumeral and lumbar regions, such defects being likely to affect the amount of muscle-derived factors supporting NMJ maintenance (Fig 7). Altogether, these results show that *Fat1,* by acting both in MNs and non-cell-autonomously in the mesenchyme, is required to shape CM and scapulohumeral neuromuscular circuits thus ensuring their anatomical and functional integrity.

## Discussion

While the processes of muscle morphogenesis and motor development occur simultaneously, what coordinates these two parallel processes is poorly understood. Here, we have dissected how the Fat1 cadherin controls neuromuscular morphogenesis. We studied this question in the context of morphogenesis of a subcutaneous muscle, the CM and the associated MN population. This system is an interesting model in which the process of myogenic differentiation is physically uncoupled from the progression of two pioneer populations - muscle progenitors and motor axons - which appear to lead the way hand in hand, both of them exposed to cues from the surrounding connective tissue (Fig 13). We show that loss-of-function of the *Fat1* cadherin simultaneously disrupt morphogenesis of this muscle, axonal wiring along this muscle and fate specification by the corresponding motor neuron population. Whereas we previously analyzed what happens when removing *Fat1* in myogenic progenitors, in the current study, we address the relative contribution of two other cell types expressing *Fat1*: these include a) the CM-innervating motor neuron population, and b) the subcutaneous mesenchymal connective tissue layer through which CM progression occurs. The present data establish that *Fat1* acts in both cell types to modulate distinct aspects of a tissue-crosstalk coupling muscle shape to MN pool specification (Fig 13): *Fat1* is required in mesenchymal cells to control muscle growth and differentiation in a non-cell-autonomous manner. This modulates the amount of a major CM-derived signal, GDNF, and consequently influences CM motor innervation and MN specification. In parallel, we show that Fat1 is required in MNs to modulate axon growth and fate specification.

**Fig 13:**
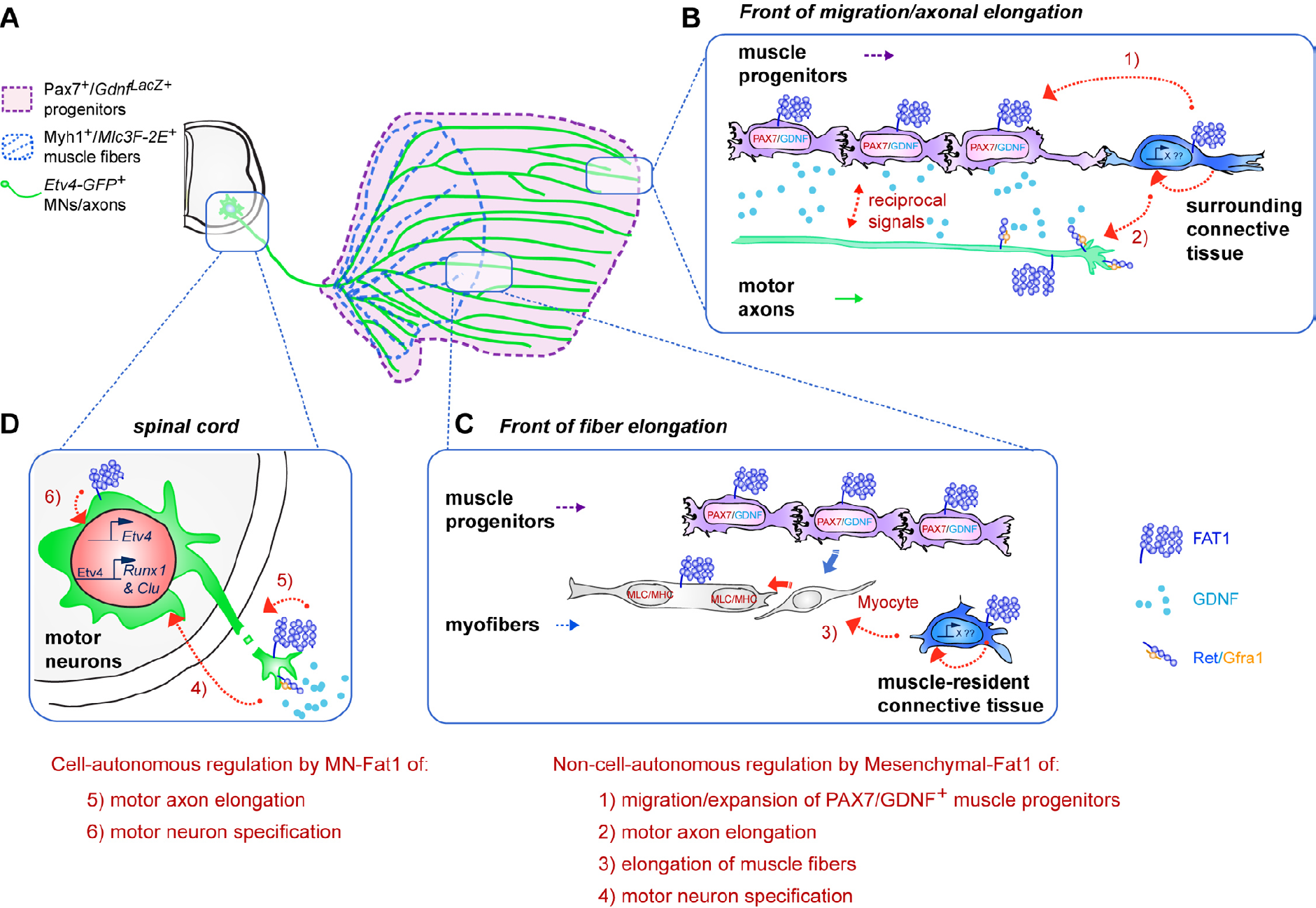
Fat1 coordinates neuromuscular morphogenesis by playing distinct roles in mesenchyme, muscles and motor neurons. **(A)** Scheme representing the CM neuronal circuit, with Etv4^+^ MNs in the spinal cord, and the CM muscle featuring the area covered by *Gdnf^LacZ+^* muscle progenitors in purple, *MLC3F-2E^+^* muscle fibers represented in blue, and CM-innervating axons represented in green. **(B)** Scheme summarizing event occurring at the migration front, where muscle progenitors and axons of CM motor neurons encounter the surrounding connective tissue, and the non-cell-autonomous functions exerted by Fat1 by acting in the mesenchyme. **(C)** Scheme summarizing event occurring at the front of fiber elongation, where new myocytes produced by the neighboring myogenic progenitors are added to elongating myofibers, and the non-cell-autonomous function exerted by Fat1 by acting in the resident mesenchymal cells. **(D)** Scheme of event occurring at the level of CM-innervating MNs, in which Fat1 cell autonomously promotes axonal growth and fine-tunes cell fate specification by modulating expression of the Etv4 target genes *Clusterin* and *Runx1* in the CM MN pools. **(E)** Summary of the roles played by Fat1 in MNs and in mesenchyme/connective tissue. Right: In muscle-associated connective tissue (both at the migration front and at the front of fiber elongation), Fat1 acts by controlling the production of a signal (Fat1 itself or a gene activated by a signaling cascade downstream of *Fat1*), which influences 1) expansion of muscle progenitors, 2) motor axon elongation; 3) myofiber elongation; and 4) MN specification. As a result of mesenchyme-specific *Fat1* ablation, the reduced CM muscle produces less GDNF, depletion of which mildly affects *Etv4* expression but severely impacts. Left: In CM MNs, Fat1 cell-autonomously modulates axon growth, and controls *Clusterin* and *Runx1* expression, either directly or as a consequence of the non-cell-autonomous impact of MNs on myogenic progenitor expansion.

### Mesenchymal *Fat1* controls CM muscle growth and impacts MN development

Ablating *Fat1* in the mesenchyme profoundly disrupts the progression of progenitor migration, and subsequently interferes not only with myogenic differentiation, but also with axonal growth and complete specification of the cognate MN pool. This suggests that *Fat1* controls the capacity of mesenchymal cells to influence: 1) the subcutaneous migration of Gdnf^+^/Pax7^+^ CM progenitors; 2) the subsequent elongation of CM muscle fibers; 3) the parallel progression of CM-innervating axons of the Etv4-expressing MNs; and 4) final molecular specialization of this MN neuron pool. The fact that MN-specific *Fat1* ablation only alters progenitor migration, but not the rate of myogenic differentiation, whereas mesenchyme-specific ablation affects both, suggests that the alteration of fiber extension observed in *Prx1-cre; Fat1^Flox/Flox^* embryos is not simply the consequence of an impaired progression of migration, but rather reflects a distinct independent function of *Fat1* in the mesenchyme, that regulates fiber elongation.

The mechanism by which *Fat1* acts in the mesenchyme, and whether this involves bidirectional Fat-Dachsous signaling, remains to be clarified. An interesting analogous situation occurs during kidney development, where a *Fat4-Dchs1/2* crosstalk controls mouse kidney morphogenesis [30–32]. In this context, Fat4 acts in the kidney stroma (analogous to the connective tissue), whereas Dchs1 and 2 are required in the Gdnf-expressing cap mesenchyme progenitors of renal tubules for their expansion (analogous to progenitors of the CM muscle) [30–32]. The mechanism identified in our study is also comparable to the feed-forward mechanism occurring in the drosophila wing imaginal disc, allowing wingless-dependent propagation of expression of the selector gene vestigial, where Fat-Dachsous signaling also contributes to several aspects, thereby coupling cell fate and tissue growth [83].

Our data establish that *Fat1* activity controls mesenchyme-derived cues acting on myogenesis. HGF and SDF1 (cxcl12) are two ideal candidates for mediating such mesenchyme-derived activity. Both factors are known to induce myoblasts motility and promote their migration towards the limb bud [62,84,85], and to act simultaneously on subsets of MNs to modulate their specification and axon guidance [46,47,86]. We recently showed that mesenchyme-specific overexpression of the Met RTK, without interfering with HGF expression and secretion, nevertheless prevented the release of biologically active HGF from mesenchymal cells, thereby leading to absence of migration in the limb [84]. This phenotype was however much more severe than the alterations of CM shape observed in mesenchyme-specific *Fat1* mutants, making it unlikely for significant changes in the amount of biologically active HGF to account for the muscle patterning defects observed in mesenchyme-specific *Fat1* mutants. The ultimate identification of mesenchyme-derived factors which production is altered by deficient Fat1 signaling will necessitate unbiased approaches such as analyses of the secretome or transcriptome of wild type and *Fat1*-deficient mesenchymal cells. In addition to seeking secreted factors, future lines of research will also evaluate whether the mesenchymal *Fat1* activity modulates the mechanical properties of the connective tissue or of the extracellular matrix it produces. Indeed, recent data showed that tissue stiffening on its own is a driver for the collective migration of neural crest cells [87].

Besides acting on myogenesis, we have established that *Fat1* activity in mesenchymal cells influence motor axon growth and MN fate specification. These non-cell autonomous consequences of mesenchymal-*Fat1* ablation on MNs and muscles could either represent independent phenotypes, or could be causally linked to one-another. Limb-innervating MNs are known to be pre-specified to find their peripheral targets [10,11], and can correctly execute pathfinding decisions in the surrounding limb mesenchyme in a context where muscles have been experimentally or genetically ablated [46,82]. In contrast, we provide experimental evidence supporting the idea that growth of MN axons innervating the CM, the trapezius or body wall muscles, is severely impacted by muscle ablation and the subsequent depletion in muscle-derived factors (S11 Fig). This is consistent with our previous work on the parallel activities of HGF/Met on muscle and MN development. Null mutation of the Met receptor abrogates muscle migration and the CM is part of the missing muscles [47,61]. This phenotype is associated with a complete absence of CM-innervating motor axonal arborization [60]. The null allele cannot distinguish whether this axonal defect is a consequence of the absence of CM muscle or of ablation of *Met* in MNs. Comparing changes in axonal patterns in null mutants and in neuronal-specific mutants indicated that *Met* is specifically required for motor innervation of another muscle (the pectoralis minor), but dispensable for the guidance of CM motor axons to and within the CM muscles [60]. Thus, the complete lack of axons matching a CM pattern in *Met* null mutant embryos is exclusively a consequence of the impaired migration of myogenic progenitors, also implying that CM motor axons along and within the CM is dependent on signals from myogenic cells. Together, these past and present elements support the possibility that altered extension of CM-innervating axons in mesenchyme-specific *Fat1* mutants could likewise be a consequence of the impaired CM muscle progression

We show that for a large part, MN phenotypes observed in mesenchyme-specific *Fat1* mutants mimic the phenotypes resulting from depletion of one essential muscle-derived factors, GDNF, which production by CM myogenic progenitors must be quantitatively affected by the reduced subcutaneous spreading. While mesenchymal *Gdnf* expression by plexus cells is unaffected, *Fat1* ablation in the mesenchyme drastically impairs the subcutaneous progression of the sheet of Gdnf^+^/Pax7^+^ progenitors (Fig 8). Our analysis of consecutive cross-sections supports the idea of a reduced number of Gdnf^+^/Pax7^+^ progenitors, raising the possibility that this effect could result not only from impaired migration, but also from impaired proliferation of the progenitor pool. As a result, MN specification defects exhibited by mesenchyme-specific *Fat1* mutants partially mimic the effect of *Gdnf* loss-of-function. Depletion of myogenic-*Gdnf* is sufficient to impair *Runx1* and *Clusterin* expression, with limited effect on *Etv4* induction. Finally, the additive effect of genetic lowering of *Gdnf* levels and mesenchyme-specific *Fat1* depletion indicates that left over *Gdnf* is a rate-limiting factor accounting for the residual induction of *Etv4* in CM motor pools in *Fat1* mutants.

These results do not exclude an alternative scenario, according to which other mesenchyme-derived factors regulated by *Fat1* activity would directly act on CM-innervating MNs to control their fate and axonal growth. The fact that we did not detect quantitative changes in the mesenchymal component of GDNF expression argues against the possibility of cell-autonomous regulation of *Gdnf* promoter by mesenchymal-*Fat1*. As discussed earlier, the likelihood for HGF amount to be drastically reduced is low, as this would have resulted in much more severe muscle defects. Thus, *Fat1*-dependent mesenchyme-derived factors influencing MN fate remain to be identified. A recent study described how regionalized mesenchyme-derived signals contribute to the acquisition of muscle-specific adaptation by proprioceptive afferents [88]. In this study, proprioceptive neuron specialization was neither affected by the MNs nor by muscle genetic ablation, pointing instead towards the contribution of mesenchymal cells. The identity of the regionalized mesenchyme-derived signals regulated by Lmx1b patterning activity, also remains unknown [88].

Which subtype of mesenchymal cells mediates such neuromuscular patterning *Fat1* activity? The importance of lateral plate-derived connective tissue for muscle patterning has been evidenced long ago through classical embryological studies [5], and is currently the subject of renewed interest [13]. The connective tissue subtypes likely to influence muscle development include tendon progenitors, at the interface between bones and muscles, but also a number of muscle associated subtypes. Genetic data supporting a contribution of the non-myogenic connective tissue to muscle development have previously been reported [13,77,89–91]. Mesenchyme-specific deletions of TCF4, β-catenin or Tbx4/5 lead to drastic alterations of muscle patterning reminiscent of the phenotypes of *Fat1* mutants [77,91]. Another example of transcription factor acting in the limb mesenchyme to pattern both muscle shapes and neuronal patterning is Shox2 [92]. These studies converge to identify CT subtypes by the combination of transcription factors they express. Among these transcription factors, Tcf4 expression highlights a subset of connective tissue at muscle extremities distinct from tenocytes, in addition to subsets of myogenic progenitors. Tcf4-expressing CT cells were shown to influence muscle growth and patterning during developmentç, but also adult skeletal muscle regeneration [77,78]. Co-cultures of Tcf4-expressing CT cells with myogenic cells indicated that these cells are capable of producing factors promoting muscle growth and differentiation in vitro [77]. However, we find that the *Fat1* expressing subcutaneous layer of connective tissue that delineates the path of CM progenitor migration/extension does not express Tcf4, but instead expresses Pdgfrα and TenascinC. Taking advantage of an inducible *Pdgfrα-cre/ERT2* line, we identify the Pdgfrα lineage as critical to for *Fat1* to influence CM growth and differentiation, since a rate of ablation of 30% of the cells in this lineage is sufficient to significantly delay progression of both progenitor expansion and myogenic differentiation. The severity of this phenotype is lower when driven by *Pdgfrα-iCre* than in the context of the broad deletion mediated by *Prx1-cre,* supporting the idea of a dosage-dependent effect affecting the concentration of secreted factors. It remains possible that some of the muscle patterning *Fat1* activity might be executed in other CT subtypes, as *Fat1* ablation in the Pdgfrα-dependent lineage did not reproduce the phenotypes observed in scapulohumeral muscles. It will be interesting in future to define the muscle-associated CT subtype that most accurately overlaps with *Fat1* expression domain and defines its domain of activity. CT at the interface between the CM muscle and the skin must be distinct from classical tendons which connect skeletal muscles to bones for force transmission. Candidate transcription factors that could highlight a signature for the *Fat1*-dependent CT subtype include Gata4, Tbx3, Osr1 and Osr2 [93,94], which were recently described as markers and essential players in defining distinct CT subtypes, respectively controlling morphogenesis of the diaphragm and of subsets of forelimb muscles [89,90,95]. The fact that *Fat1* deficiency did not alter diaphragm development or lead to diaphragmatic hernias such as those occurring in the absence of *Gata4* make it unlikely for *Fat1* to be required in the Gata4^+^/diaphragm CT subtype. In contrast, although CM development was not directly assessed in these studies, the changes in limb muscle patterning observed upon deletion of *Tbx3* or *Osr1* in the lateral-plate-derived lineage *(Prx1-cre)* include selective alterations of muscle attachment sites or fiber orientation reminiscent of those occurring in *Fat1* mutants [89,95].

The role played by the Pdgfrα-dependent lineage in *Fat1*-driven developmental myogenesis is interesting in light of the emerging role of muscle-resident CT lineages for adult skeletal muscle. Pdgfrα expression distinguishes a population of muscle-resident cells called fibroadipogenic progenitors (FAP) [96,97]. These cells do not directly contribute to the myogenic lineage, but are bipotential progenitors, capable of giving rise to adipocytes and to connective tissue fibroblasts [96,97]. Whereas FAPs are quiescent under homeostatic conditions, muscle lesions activate their proliferation and differentiation [97], in addition to activating muscle stem cells proliferation and to trigger the production of new muscle fibers [2]. This damage-induced FAP recruitment was shown to contribute to fibrosis and fat infiltrations, in the context of either acute lesions, or of chronic degeneration/regeneration cycles occurring in a mouse model of muscle dystrophy (mdx mice) [98,99]. In co-cultures of FAPs and muscle stem cells, FAPs promote myogenic differentiation, indicating that the recruitment and activation of FAPs in degenerating muscles could contribute to the efficiency of muscle repair [97]. Future work will establish whether *Fat1* activity is also necessary for the adult FAP lineage to exert its myogenesis-promoting activity during muscle homeostasis and repair.

### Motor neuronal Fat1 – dissecting direct and indirect contributions to axonal growth, MN specification and muscle spreading

We find that in addition to its mesenchymal contribution, *Fat1* plays additional cell-autonomous functions in the MN pool innervating the CM muscle. Fat1 protein is selectively expressed in the CM MN pool, likely distributed along their axons. MN-specific *Fat1* deletion negatively influences axonal growth of CM-innervating axons, and alters acquisition of MN fate characteristics. Unexpectedly, although MN-specific *Fat1* ablation does not significantly influence the rate of myogenic differentiation, our quantitative analysis uncovers a significant delay in the subcutaneous progression of muscle progenitor migration. This uncovers an unsuspected role of motor axons in the rate of progression of muscle progenitor migration.

The MN specification and axonal growth defects occurring in absence of MN-*Fat1* may reflect a cell-autonomous role of Fat1 signaling in MNs. Future lines of investigation will determine whether such Fat1 signaling cascade involves interactions with Dachsous cadherins, and how it influences axon growth and modulates *Runx1* and *Clusterin* expression. It will be relevant to discriminate whether MN-*Fat1*: 1) modulates the response to - or reception of - GDNF by controlling receptor expression, or efficiency of the GDNF/Ret signal transduction; 2) impinges on Etv4 transcriptional activity, by regulating its phosphorylation; or 3) acts directly on target gene expression via other cascades known to be downstream of FAT cadherins such as Hippo/YAP or the transcriptional repressor atrophin. As a protocadherin, Fat1 may also refine neural circuit assembly by mediating axonal self-recognition similar to functions of clustered protocadherins [100,101]. Alternatively, the two phenotypes could simply reflect the consequence of the reduction in GDNF levels resulting from the altered progenitor progression also resulting from MN-specific *Fat1* ablation. Both phenotypes are reminiscent of the consequences of *Gdnf* ablation (as illustrated here and in [39]). Gdnf is also known for exerting an axon growth promoting role [102], and in MN-specific *Fat1* mutants, the length of CM motor axons remains correlated with the size of the *Gdnf*-expressing progenitor sheet.

We have uncovered that CM motor axons progress hand in hand with Gdnf^+^/Pax7^+^ muscle progenitors, and show that disrupting *Fat1* activity in MNs interferes with the rate of subcutaneous progression of CM progenitors. This effect is however less predominant than the dependency of CM progenitor progression on mesenchymal *Fat1* activity. Such an influence of MNs on myogenic progenitors is counter intuitive, since previous studies supported the idea that muscle development can occur normally in absence of motor innervation [103,104]. These studies were examining NMJ maturation in the diaphragm muscle at fetal stages after genetic depletion of MNs. They showed that whereas a pre-pattern of Acetylcholine receptors (AChRs) prefiguring future synapses does occur in the diaphragm in absence of motor axons, the latter are required for subsequent NMJ maturation. The maturation-inducing activity elicited by MNs on muscles is mediated by signals such as Agrin, inducing AChR clustering [105]. These studies do not exclude however that the dynamics of muscle morphogenesis could have been altered at primary myogenesis stage in some vulnerable muscles, and compensated at secondary myogenesis stages. Such influence of MNs on the development of another tissue-type is also reminiscent of the role exerted by MNs, by way of VEGF secretion, on vascular development [106]. Finally, since axonal elongation is concomitant with spreading of the CM myogenic progenitor, motor axons and myogenic progenitors collectively expose themselves to the mesenchymal landscape, and to mechanical properties of the ECM, such as its stiffness. Interfering with the progression of one of the two cell types thus exposes the other on its own to the mesenchyme, rendering it less permissive, and slowing the collective progression.

In spite of this delayed progenitor progression, myogenic differentiation is not impacted in absence of MN-*Fat1* and the resulting anatomy of the adult CM is normal. MN-specific *Fat1* ablation leads to significant functional impact on NMJ integrity in absence of anatomical abnormalities of the muscle and of changes in muscle fiber diameter. This highlights the importance of MN-*Fat1* in ensuring NMJ integrity. This effect could be the direct or indirect consequence of: 1) *Clusterin* or *Runx1* lowering in MNs, 2) reduced GDNF levels in muscle progenitors, 3) a reduced number of muscle progenitors persisting in the adult muscle; or 4) the deregulation of other target genes influencing NMJ integrity, that remain to be identified.

### Relevance for FSHD

Muscle phenotypes caused by disruption of *Fat1* functions are highly regionalized and the topography is reminiscent of the map of muscles undergoing degeneration in Facioscapulohumeral dystrophy (FSHD) [14]. FSHD is the second most frequent muscular dystrophy, characterized by the selective map of affected muscles, involving facial expression and shoulder muscles [107]. In most cases, FSHD is caused by genomic alterations on chromosome 4q35 leading to excess production of DUX4, a transcription factor encoded in a repeated macrosatellite, normally active only during zygotic gene activation [107,108]. This overexpression is permitted by a genetic context simultaneously involving 1) a polymorphism stabilizing DUX4 RNA by creating a polyA signal [109]; and 2) the loss of epigenetic DUX4 repression [110]. Such epigenetic derepression results: a) in the most frequent form of the disease (FSHD1), from deletions that shorten the D4Z4 repeat array [110]; b) in less frequent cases (FSHD2), from mutations in genes involved in modulating the chromatin architecture, (SMCHD1 [111]), or in DNA methylation (DNMT3A [112]). DUX4 exerts a toxic effect on muscles by deregulating transcription of multiple targets [113–115]. Individuals carrying a pathogenic context exhibit a high variability in symptom severity [116,117], implying that other factors behave as disease modifiers.

The similarity between muscular and non-muscular phenotypes of *Fat1* mutants and FSHD symptoms indicates that *Fat1* deficiency phenocopies some FSHD symptoms [14], raising the possibility that *FAT1* dysfunction in FSHD-relevant tissue types could contribute to some of these phenotypes. This possibility was supported by the identification of pathogenic *FAT1* variants in human patients, in which FSHD-like symptoms were not caused by the traditional genetic causes of FSHD [14,118]. Furthermore, *FAT1* is located in the vicinity of the FSHD-associated D4Z4 array on 4q35, and its expression is downregulated in fetal and adult muscles of FSHD1 and FSHD2 patients [14,119], suggesting that traditional FSHD-causing contexts may not only enhance *DUX4* expression but also lead to lowered *FAT1* expression in affected muscles. Although exogenous DUX4 can downregulate *FAT1* expression when exogenously expressed in human myoblasts, shRNA-mediated *DUX4* silencing in FSHD1 myoblasts did not restore *FAT1* expression levels [119]. Thus, *FAT1* deregulation by a pathogenic 4q35 context is not attributable to DUX4 in FSHD1 myoblasts, and might be caused by DUX4-independent mechanisms. First, the loss of epigenetic repression of *DUX4* might be due to changes in the chromatin architecture, resulting from deletions of anchoring sites for chromatin folding proteins such as CTCF or SMCHD1 [111,120]. Given the proximity of *FAT1* with the D4Z4 array, the same changes in chromatin architecture could simultaneously lead to lowered *FAT1* levels. Second, among genetic variants in *FAT1,* a copy number variant (CNV) deleting a putative regulatory element in the *FAT1* locus, which has the potential to deplete *FAT1* expression in a tissue-specific manner, was enriched not only among FSHD-like patients but also among classical FSHD1 and FSHD2 [14,119]. Furthermore, given the chromosomal proximity, genetic alterations in *FAT1* have a high probability to co-segregate with a pathogenic FSHD context occurring on the same 4q chromosome. Support for this came from the recent identification of pathogenic *FAT1* variants associated with a classical FSHD1 context [121]. Thus, the combination of a pathogenic 4q35 context with pathogenic *FAT1* variants, whether functional or regulatory, has the potential to synergize to reduce *FAT1* expression and/or activity below a functional threshold, potentially contributing to worsen FSHD symptoms. Other genetic modifiers affecting chromatin methylation [111,112] and/or architecture of the 4q35 arm [122], hence affecting gene regulation, may simultaneously impact *DUX4* and *FAT1* expression.

Therefore, elucidating *Fat1* functions during development could provide instructive information to guide research on FSHD pathogenesis. Our findings indicate that myoblasts may not be the only relevant cell type where *FAT1* downregulation must occur to alter muscle development, and identify mesenchyme in developing embryos as another cell type in which *Fat1* disruption is sufficient to cause muscle shape phenotypes with an FSHD-like topography. Whether *FAT1* is indeed deregulated in connective tissue of FSHD patients, and whether such regulation change is accounted for by DUX4, or by 4q35 chromatin architecture changes, remains to be established. Several FSHD mouse models exist, some of them engineered for inducible and tissue-specific *DUX4* overexpression, and will be instrumental to assess to what extent and through which mechanism loss of *Fat1* function could synergize with enhanced *DUX4* levels [123–126]. Furthermore, our results also identify GDNF as a muscle-derived signal, which lowering is associated with muscular phenotypes of *Fat1* mutants and has consequences at least on neuronal development and possibly on NMJ integrity in adult muscle. Interestingly, GDNF appeared significantly down-regulated in FSHD biopsies, in at least two reports [127,128]. Another study suggests instead that FSHD involves enhanced Ret signaling in myoblasts, and proposes Ret inhibitors as a possible therapeutic strategy [129]. In view of these reports and together with our present findings, it may be relevant to determine whether and to what extent modulation of GDNF/Ret signaling, or of other mechanisms influenced by Fat1 signaling, can have any benefit to alleviate adult muscle symptoms and improve quality of life of FSHD patients. Finally, whereas developmental alterations such as those occurring in the Etv4/GDNF circuit might constitute a topographic frame in which muscles have deficient functional output, it will be necessary to understand whether and how this renders the muscles more susceptible to degeneration in adults in an FSHD context.

## Experimental Procedures

### Mouse Lines

All the mouse lines and genetic tools used here are described in the supplementary S1 Table, providing for each of them a link to the mouse genome informatics (MGI) information page, a reference describing their production, some brief explanations on what they are, how they were made, and how they are used in the present study. The alleles of the *Fat1* gene used in this study included a genetrap knockout/knock-in allele *Fat1^KST249^* (referred to as *Fat1^LacZ^)* [130], as well as a constitutive knock-out *Fat1^ΔTM^* allele (here referred to as *Fat1*^-^) and the conditional-ready *Fat1^Fln^* allele from which it derived, that we previously described [14]. As the original conditional-ready *Fat1^Fln^* allele still carried a neo selection cassette flanked by FRT sites [14], we have crossed *Fat1^Fln/+^* mice with mice carrying the Flp recombinase (*ActFlpe* mice, official name Tg(ACTFLPe)9205Dym [131]), thereby producing a new conditional-ready allele referred to as *Fat1^Flox^,* used throughout this study. The Cre lines used for tissue-specific ablation were either transgenic lines (official MGI nomenclature Tg(gene-cre)author), or knock-in/knockout alleles (official MGI nomenclature gene^tm(cre)author^). These include: *Olig2-cre* (knock-in/knock-out Olig2^tm1(cre)Tmj^, [72]); *Wnt1-cre* (transgenic Line Tg(Wnt1-cre)11Rth [75]); *Prx1-cre*(transgenic Line Tg(Prrx1-cre)1Cjt [73,74]), *Pax3-cre* (knock-in/knock-out line *Pax3^tm1(cre)Joe^* [64]), and the tamoxifen-inducible *Pdgfrα-iCre* line (BAC transgenic line Tg(Pdgfrα-cre/ERT2)1Wdr [79]. To produce embryos or mice *Cre^+^; Fat1^Flox/Flox^,* all cre alleles were transmitted by males only to avoid female germline recombination, by crossing *Cre^+^; Fat1^Flox/+^* males or *Cre^+^; Fat1^FloxFlox^* males with *Fat1^FloxFlox^* females, the latter carrying one of the reporter genes described below *(MLC3F-2E-LacZ, Gdnf^LacZ^, R26-YFP).* Knock-in/knock-out *Pea3-lacZ* mice (here referred to as *Etv4^-^)* were used with permission of S. Arber and T. Jessell, and genotyped as previously described [41]. Transgenic Mice carrying the BAC transgene *Etv4^GFP^* have been described previously [60]; knock-in/knock-out *Gdnf^LacZ^* mice were used with permission of Genentech, [39,132]; knock-in/knock-out mice carrying the *Met^d^* allele produce a signaling dead version of the Met receptor [61]. Additional lines used in this study are: *Rosa26-YFP* mice (Gt(ROSA)26Sor^tm1(EYFP)^Cos line (Jackson laboratory mouse strain 006148, [66]); *MLC3F-2E-LacZ* mice (transgenic line Tg(Myl1-lacZ)1Ibdml, [58,59]); *HB9-GFP* (transgenic line Tg(Hlxb9-GFP)1Tmj allele, [133]); *Myf5-cre* mice (Myf5^tm3(cre)Sor^ line (Jacson laboratory mouse strain 007893, [80]); *R26^Lox-LacZ-STOP-Lox-DTA^* mice (also called *R26^DTA^,* or *Gt(ROSA)26Sor^tm2(DTA)Riet^* line; [81]), All lines were maintained through back-crosses in B6D2 F1-hybrid background.

Animals were maintained and sacrificed in accordance with institutional guidelines. When collecting embryos, adult mice were euthanized by cervical dislocation. For collection of adult muscle samples, mice were first anesthetized with lethal doses of Ketamin and Xylasine, and euthanized by transcardiac perfusion with 4% PFA.

### X-gal and Salmon-Gal staining for β-galactosidase activity

Staining for β-galactosidase activity was performed using two alternative methods. To detect *LacZ* expression in embryos with the *MLC3F-2E-LacZ* transgene or *Fat1^LacZ^* allele, we performed classical X-gal staining with standard protocols using Potassium Ferricyanide and Potassium Ferrocyanide. Embryos were then post-fixed in PFA, partially clarified in 1 volume glycerol /1 volume PFA4%; and mounted in 100% glycerol on glass slides with depression, under coverslips with glass spacers for imaging. When detecting *LacZ* expression in embryos carrying the *Gdnf^LacZ^* allele or *Fat1^LacZ^* allele on cryosections, we selected the more sensitive method using Salmon Gal (6-Chloro-3-indolyl-beta-D-galactopyranoside; Appolo scientific UK, stock solution 40mg/ml in DMSO) in combination with tetrazolium salt (NBT (nitro blue tetrazolium, N6876 Sigma), stock solution 75mg/ml in 70% DMF), using the method we previously described [46]. Briefly, after initial fixation (for *Gdnf^LacZ^:* 15 minutes in 1%PFA; 0.2% Glutaraldehyde in PBS), embryos were rinsed in PBS, and incubated overnight, in order to avoid background staining, at 37°C in a solution containing 4 mM Potassium Ferricyanide, 4 mM Potassium Ferrocyanide, 4mM MgCl2; 0.04% NP40 but without substrate. Embryos were then rinsed 3 times in PBS, and incubated in a solution containing Salmon Gal (1mg/ml), NBT (330 μg/ml), 2mM MgCl2; 0.04% NB40, in PBS. After staining completion, embryos/sections are rinsed in PBS and post-fixed in 4% PFA. For *Gdnf^LacZ^,* the staining reaction is completed in 30-40 minutes. For staining of cryosections of *Fat1^LacZ^* embryos, the staining reaction is completed in 1-2 hours.

### Simple and double in situ hybridizations

Embryos were collected in cold PBS, and when necessary, spinal cords were dissected at that stage. Dissected spinal cords, and the remaining embryo trunks, or whole embryos were then fixed overnight in 4% paraformaldehyde (in PBS), dehydrated in methanol series, and stored in methanol at −20°C. In some cases, embryos were embedded in gelatin/Albumin, and sectioned on a vibratome (80um sections). These floating sections were then dehydrated in methanol series and treated like whole-mount embryos. In situ hybridizations were done as previously described [47]. Briefly, after methanol dehydration, tissues (embryos, spinal cords or vibratome sections) were rehydrated in inverse methanol series, treated with proteinase K, hybridized with Digoxigenin-labeled probes, washed with increasing stringency, incubated with anti-digoxigenin antibodies conjugated with alkaline phosphatase (Roche), and developed with NBT/BCIP substrates. For double in situ hybridizations, hybridization was done by mixing a Digoxigenin-labeled and a Fluorescein-labeled probe. Revelation of the two probes was done subsequently. We first incubated the samples with the AP-conjugated anti-digoxigenin antibody (1/2000, Roche), washed, and stained using NBT/BCIP (blue color). Samples were post-fixed, cleared with 50% glycerol, flat-mounted to be photographed with the first color only (blue). To start the second staining, the samples were then transferred back in PBT (PBS, 0.1% Tween-20). The remaining alkaline phosphatase was then inactivated for 10 minutes with 0.2M Glycine pH2.2; 0.1% Tween-20. Samples were further rinsed 4 times in PBT, after which they were incubated with AP-conjugated anti-fluorescein antibody (Roche, 1/2000). After washing, this second staining was revealed using INT/BCIP substrate (Roche, red color). Samples were post-fixed again, cleared in 50% glycerol, and flat-mounted for visualization.

### Immunocytochemistry

Embryos were collected in cold PBS and fixed 2-4 hr in 4% PFA on ice. Adult mouse tissues were collected after perfusing anesthetized mice with 4% PFA, and were post-fixed for 2-4h in 4% PFA. The samples were rinsed in PBS, cryoprotected in 25% sucrose-PBS solution, embedded in PBS, 15% sucrose / 7.5% gelatin, and cryostat sectioned (10 μm for embryos, 30 μm for NMJ analysis in adult muscle longitudinal sections). For Fluorescent immunohistochemistry, slides were thawed in PBS, then incubated in PBS, 0.3% triton-X-100, bleached in 6% H2O2 for 30 minutes, rinsed again in PBS, 0.3% triton-X-100. For some antibodies (see S2 Table), heat induced epitope retrieval was performed in 0.1M Citrate buffer pH6 at 95°C for 5-30 minutes (specific time depending on the antibody used). Slides were rinsed again 3x in PBS, 0.3% triton-X-100, and then incubated over night at 4°C with primary antibodies in 20% Newborn Calf serum, PBS, 0.3% triton-X-100 in a humidified chamber. Primary antibodies or detections tools used are described in S2 Table. After extensive washing in PBS, 0.3% triton-X-100, slides were incubated with secondary antibodies in blocking solution as above, for 1h30 at room temperature in a humidified chamber. Secondary antibodies used were conjugated with Alexa-555, Alexa-488 or Alexa-647 (see S2 Table). After extensive washing in PBS, 0.3% triton-X-100, slides were quickly rinsed in distilled water and mounted in mounting medium with ProlongGold antifade reagent and DAPI. Slides were kept at 4°C, and imaged with a Zeiss Z1 Microscope equipped with Apotome for structured illumination, using Zen software. Whole-mount anti-neurofilament staining of E12.5 embryos was performed as previously [46], using the mouse monoclonal Anti-neurofilament (2H3) antibody (bioreactor version, dilution 1/300), and anti-mouse-HRP secondary antibody (Sigma). Briefly, embryos were fixed 2-4h in 4% PFA (when relevant, an X-gal staining was also carried out), rinsed in PBS, then transferred in Dent’s fixative (80% Methanol; 20% DMSO), and kept at 4°C until use. A bleaching step with hydrogen peroxide was carried out by incubating embryos overnight in a solution with 1 volume 30% H2O2; 4 volumes Dent’s fixative. All washes were made in PBS, 1% triton X-100. Antibody incubations were done by incubating at least 3 days (at least one at room temperature) in a blocking solution containing 79% newborn calf serum, 20% DMSO, 1% triton X-100, and thimerosal. The final HRP color reaction was performed with DAB tablets (SIGMA) and H_2_O_2_, after rinsing in 0.1M Tris pH7.4. After staining completion, embryos were dehydrated in methanol, cleared in 2 volumes Benzyl-benzoate / 1 volume Benzy-Alcohol mix (BB-BA mix; once embryos are in 100% methanol, they are transfered in a mix 50%vol methanol; 50% volume BB-BA, for 30 minutes, the twice in 100% BB-BA mix), cut in half and mounted on glass slides with depression, under coverslips with glass spacers for imaging.

### Tamoxifen induction

In cases of crosses with the *Pdgfrα-iCre* line, Tamoxifen induction was performed by intra-peritoneal (IP) injection in pregnant females. Tamoxifen (T5648, Sigma) was dissolved first in ethanol (100mg/ml) under agitation (in a thermomixer) for 2h at 42°C, then diluted in sterile corn oil (C8267, Sigma) at 10mg/ml, kept under a chemical hood at room temperature to let the ethanol evaporate, and aliquoted (2ml aliquots) for storage at −20°C. One working aliquot was kept at 4°C for 1-2 weeks. To achieve recombination in mesenchyme-derived loose connective tissue, two successive injections were performed in pregnant females, first at E9.75, at a dose of 50 μg/g (per gram of tissue weight), and second the following day at a dose of 100 μg/g. Administration was done by IP injection, using a 26G syringe. Efficiency of recombination was assessed using the reporter line R26-YFP, first by following overall intensity YFP fluorescence on freshly dissected embryos, and later by performing anti-YFP immunohistochemistry on cryo-sections, combined with markers of tissue-types.

### Image processing for quantification of signal intensity

Quantifications of signal intensity were performed using the image J software (NIH) as previously described [46,84]. Briefly, color images were first converted to grey scale and inverted to negative scale (a scale with white for highest signal intensity (250) and black for the lowest intensity (0)). An area of fixed pixel width was cropped, and signal intensity was measured with ImageJ along a horizontal band of a the width of the cropped image (500 pixels, matching half a flat-mounted spinal cord, along dotted lines in quantified pictures, always ending in a fixed positional landmark, such as the spinal cord midline), and height of a few pixels. After background subtraction (background being the baseline value of the control samples in the same experiment, and baseline being the minimum value in a fixed window, systematically avoiding points such as the midline, where changes in tissue thickness influence background intensity), and threshold subtraction (to avoid negative values), the values were expressed as a percentage of the mean maximal amplitude of the control samples used in the same experiment (so that experiments developed independently can be normalized to their respective controls, and pooled). A mean distribution along the line is generated by averaging the normalized intensities between several samples of each genotype (considering left and right sides separately, the number of spinal cord sides used, for each plot being indicated in the corresponding figures) to generate an average signal distribution plot, shown ± standard deviation). The total signal intensity value was also calculated for each sample by measuring the area under the curve, and plotted individually as percentage of the mean control value.

### Area measurements

Area measurements were used to assess CM subcutaneous progression and NMJ size. Whole embryo images were acquired on a Leica Stereomicroscope, using Axiovision software. We used the area/distance measurement tools in the Axiovision software. For each embryo side, we measured the area occupied by the CM, detected with *MLC3F-2E-LacZ* by X-Gal staining, with *Gdnf^LacZ^* using Salmon Gal staining, or with anti-neurofilament immunohistochemistry. As reference measurements for normalization, we measured structures that grow proportionally with the embryo age. For this we measured, respectively, the area occupied by body wall muscles *(MLC3F-2E-LacZ),* the length of the trunk between two fixed *Gdnf^LacZ+^* landmarks (shoulder to posterior limit of the hindlimb), and the dorso-ventral length of a given thoracic nerve (T10 or T9) for anti-neurofilament stainings. For all measurements, the first graphs show the raw values, whereas the second plots represent the normalized area, obtained by calculation the ratio CM area/reference value, and dividing this value by the median ratio for a set of control embryos for normalization. For *LacZ*-stained embryos, measurements were done on embryos irrespective of the storage time and medium (PBS/PFA or glycerol, with different degrees of tissue shrinkage), normalization was typically done by comparing embryos of the same litter and mounted/imaged at the same time. For anti-neurofilament staining, as all embryos were stored in BB-BA (where tissue shrinkage is invariable) normalization was done by ranking embryos by size range (according to the thoracic nerve length), and comparing embryos in the same size range.

For NMJ area measurements, immunohistochemistry was performed on 30 μm thick longitudinal cryosections of the dorsal CM muscle to detect the neuromuscular junctions (with α-bungarotoxin (BTX)), the nerve end-plate (with anti-Neurofilament antibodies), and the muscle fiber F-actin (with Phalloidin) (see S2 Table for reagents). NMJ images were acquired on Zeiss Z1 Microscope equipped with Apotome for structured illumination, and processed using Zen Imaging software (Zeiss) or Axiovision software. After NMJ Image acquisition, a Maximal Intensity Projection image was generated. The latter was converted from .czi to .zvi file, and we used the area measurement tool in the Axiovision software. For each mouse, a minimum of 20 NMJs were measured in the dorsal CM, using pictures acquired at magnifications ranging from 10x to 63x (keeping the scale adapted to the magnification to measure areas in μm^2^), and the percentage of NMJs matching a set of area ranges was determined per mouse. As control, on the same set of images, the diameter of each muscle fiber was also measured (on longitudinal sections, the MIP shows the maximal width across the section thickness, thus accurately measuring fiber diameter). Even when no specific staining of the fiber was performed, background levels of the BTX staining are sufficient to accurately detect muscle fibers and measure their width). For each mouse, around 20-50 fiber diameters were measured, and the percentage of fibers matching a set of diameter ranges was determined in each mouse. The NMJ area and fiber diameter size distribution graphs in Fig 12 represent mean percentage in each area range for each genotype ± SEM, indicating the number of mice used next to the genotype indication). Control data from *Fat1^FloxFlox^* mice are shown twice (in blue) in both graphs, to make readability easier.

### Grip strength measurements

Grip tests were performed on adult mice using the Grip test apparatus (Bioseb), which measures the maximum force exerted by the mouse as it pulls a sensitive device (grid) until release of the grip. We measured either forelimb strength only or the cumulated force of forelimbs + hindlimbs (when a mouse uses all 4 limbs to pull the measurement grid). As raw force in the measured unit (Newton) was significantly different between males and females (controls), the data were normalized for each gender and genotype by dividing the measured value by the average value of the control group of the same gender. This allows expressing the output in percentage of the control value, set as 100% in average, hence allowing to pool data from males and females. In the results shown in Fig 12, male and female data have been pooled, as the separate analysis of both genders gave similar results. Three to four measurements were made for each mouse, and an average value was calculated. In both the pooled and non-pooled graphs, each dot represents the average value for one mouse. Statistic tests were carried out on the average values

### Statistical analysis

StatEL (add-in to Excel, by Ad Science, Paris) was used for statistical analyses. For comparisons between two groups, differences were assessed either using the unpaired Student t-test when data were showing a normal distribution and equal variance, or using the non-parametric Mann-Whitney test otherwise. All p values are indicated in the figures, except for Fig 9, where * indicates p value < 0.01, whereas ** indicates p value < 0.001. Differences were considered significant, when p<0.05. For signal intensity distribution curves, average values ± standard deviations are shown (in light blue). For dot plots, all individual data are plotted (one per sample), and the median value is shown with a black bar.

## Author Contributions

FH conceived the project, obtained funding, performed 95% of the experiments, analyzed the data, prepared all the figures and wrote the paper.

## Acknowledgements

I thank F. Maina and R. Kelly for comments on the manuscript and suggestions throughout the work; F Maina for *Met^d^* mutant mice; T. Jessell, and S. Arber for *Etv4* null and *HB9-GFP* mice, Genentech for *Gdnf^LacZ^* mice, Irina Dudanova for sending *Prx1-cre* mice, Aziz Moqrich for *Wnt1-cre* mice; Andrea Huber-Brösamle for *Olig2-cre* and *R26^DTA^* mice, with permission of T. Jessell; Pascale Durbec for providing the *Pdgfrα-Cre/ERT2* mice (from William Richardson). *Scleraxis* RNA probe was provided by Delphine Duprez, *Clusterin* RNA probe by C.E. Henderson and K. Dudley; *Runx1* probe obtained from the RZPD repository, *Gdnf* RNA probe cloned by B. Herberth. Antibodies against Pax7 (developed by A. Kawakami), 2H3 (by T. Jessell), and MF20 (by D.A. Fischman), were obtained from the Developmental study hybridoma bank (DSHB), created by the NIHCD of the NIH, kept under the auspices of the Iowa University. Nathalie Caruso for mouse colony genotyping and technical support in early phases of this study; Dominique Fragano for mouse colony PCR genotyping; Virginia Girod-David, and the IBDM animal house staff for help with mouse husbandry at the IBDM; Angela K Zimmermann for contribution to histological analysis for part of Fig 12B (top graph). Imaging was performed using the PiCSL-FBI core facility (IBDM, AMU-Marseille, France) supported by the Agence Nationale de la Recherche through the ‘Investments for the Future’ program (France-BioImaging, ANR-10-INBS-04). This work was supported by grants to FH from the AFM-Telethon (Association Française contre les myopathies), FSHD global research foundation, and FSH society.

## 6 SUPPORTING INFORMATION

**S1 Data.** Excel file with values used to make all plots in all figures.

**S1 Table: List of mouse lines used in this study,** recapitulating their mouse genome informatics nomenclature, reference of origin, and brief description of how they were made.

**S2 Table: List of primary antibodies and other staining reagents**, including provider, references, species, IgG type, dilutions, requirement for antigen retrieval, and secondary antibody.

**S1 Fig:**
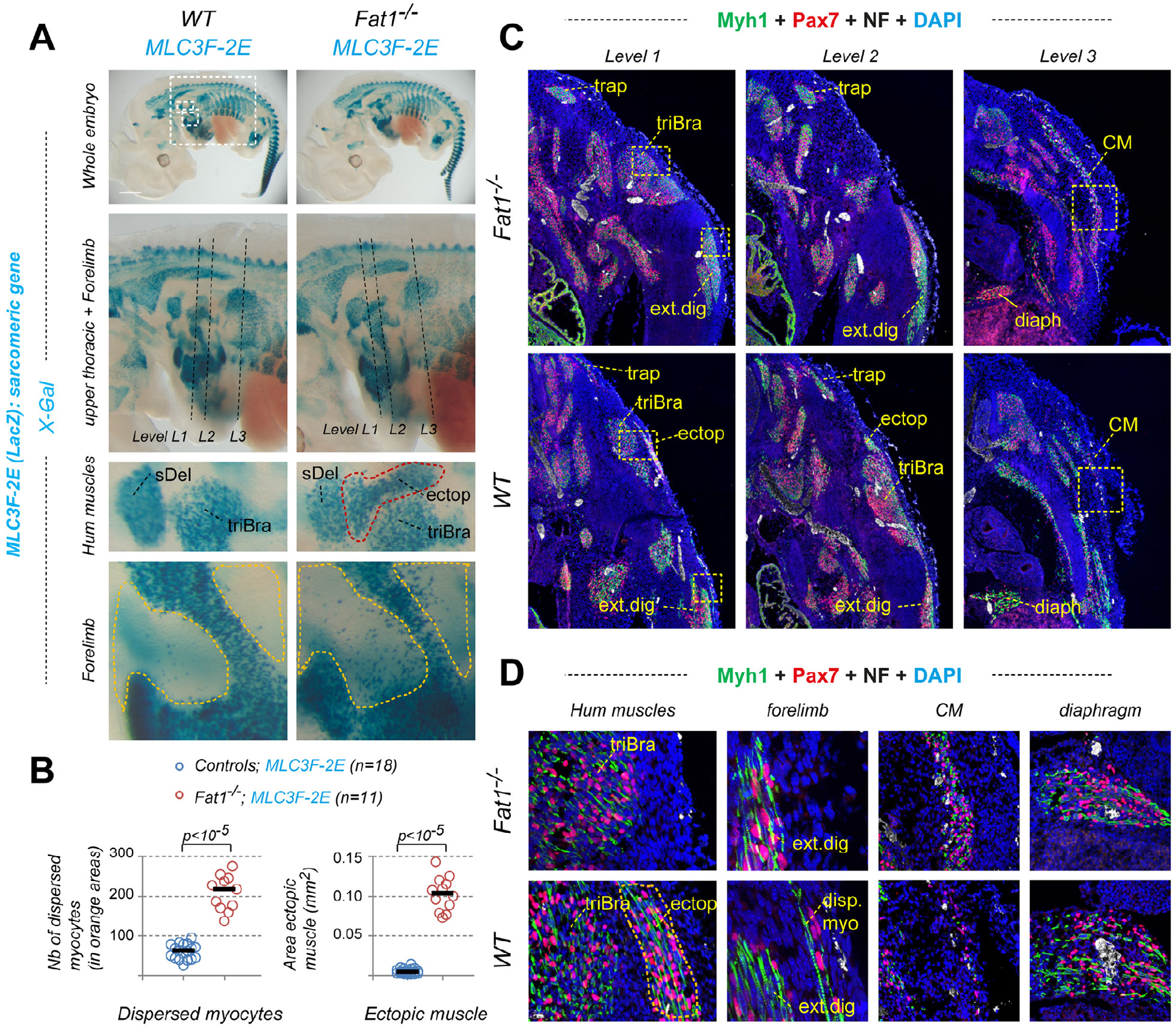
**(A)** Whole-mount β-galactosidase staining was performed on *Fat1^+/+^;* (left) and *Fatl^-/-^* (right) embryos carrying the *MLC3F-2E* transgene. Top images represent a side view of entire embryos, and indicate the position of higher magnification areas, including that of top pictures in Fig 1A,B, as well as the upper forelimb region (whith scapulohumeral muscles), and distal forelimb, showing a region where dispersed myocytes accumulate in *Fatl^-/-^* embryos. The second row images highlight a closer view of the upper thoracic and forelimb region. The dotted lines indicate for each genotype the levels and angle corresponding to the 3 consecutive sections shown in **(C)** and **(D)**. **(B)** Quantification of the number of dispersed myocytes found in orange areas in the forelimb (Left plot) and of the area occupied by the ectopic humeral muscle (ectop) appearing between the spinodeltoid (Del) and the triceps brachii (TriBra) muscles (right plot). These data reproduce and confirm our own previous results. Underlying data are provided in S1 Data. **(C, D)** Cross sections of control *Fat1^+/-^* and mutant *Fat1^-/-^* E12.5 embryos, featuring three consecutive sections at forelimb levels (Level 1 and level 2) and upper thoracic level (Level 3), immunostained with antibodies against Pax7 (red), Myh1 (Type I myosin heavy chain, green), Neurofilament (white), and with DAPI (blue). Images in **(D)**, represent high magnification views of the area highlighted with the yellow dotted square in **(C)**. These data confirm 1) the severe reduction in thickness of the CM muscle (level 3, and higher magnification in **(D)**),2) the presence of a robust ectopic muscle next to the triceps brachii (levels 2 + 3, and higher magnification in **(D)**), 3) the presence of dispersed myogenic progenitors and muscle fibers in ectopic subcutaneous position in the forelimb (image in **(D)** shows higher magnification of an area between the digit extensors and the skin). Lack of obvious phenotype in the diaphragm is also shown.

**S2 Fig:**
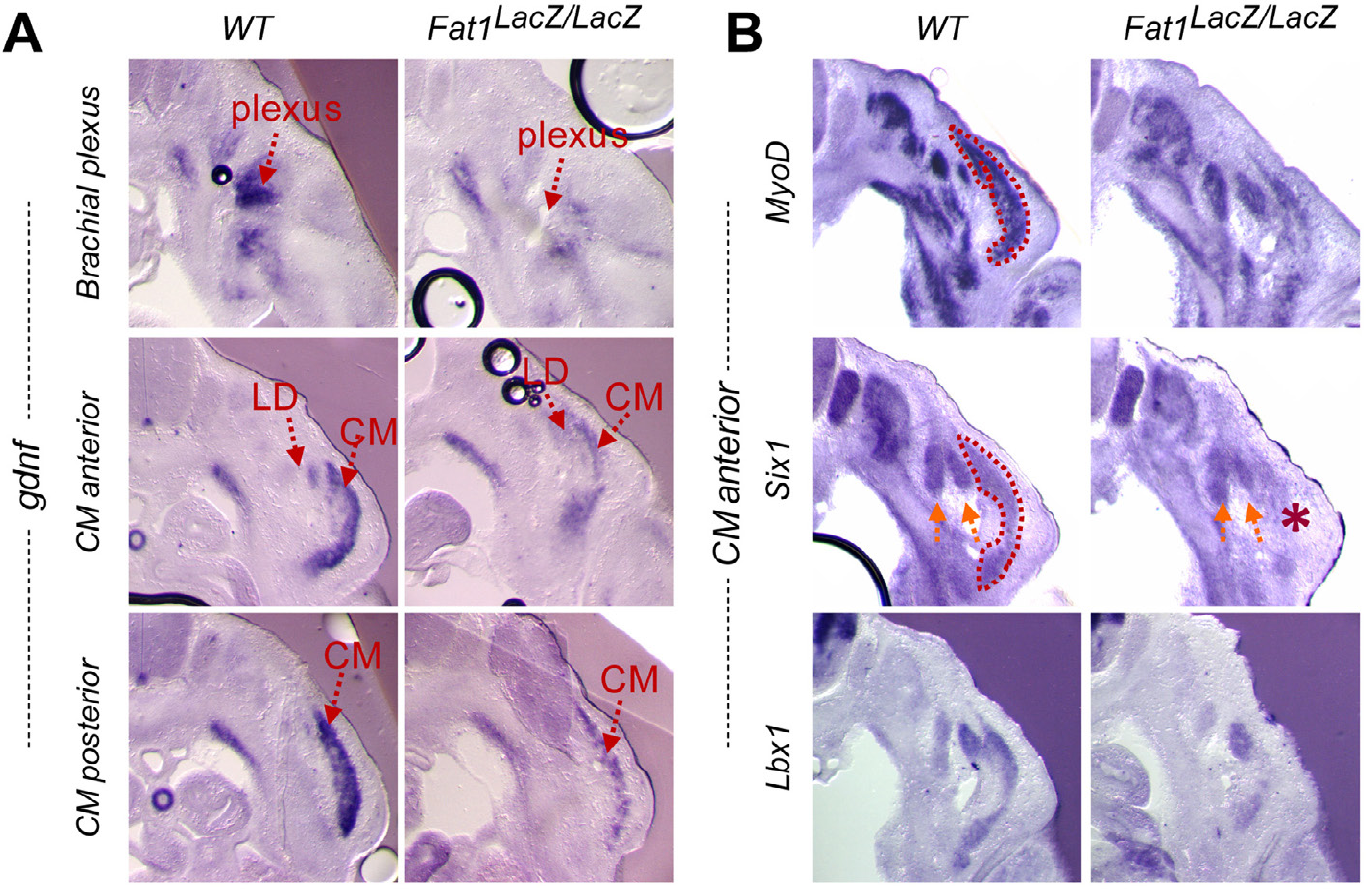
Analysis of muscle phenotype in *Fat1^LacZ/LacZ^* embryos. Expression of *Gdnf* **(A)**, and of three markers of developing muscles *(MyoD, Sixl,* and *Lbxl,* **(B)**), was analyzed by in situ hybridization on floating vibratome sections of E12.5 wild type (left panels) and *Fat1^LacZ/LacZ^* embryos (right panels). **(A)** Gdnf expression is visualized at 3 successive anteroposterior levels, showing a hotspot at the brachial plexus (mesenchymal cells around passing nerves), where Gdnf expression is drastically reduced by the absence of *Fat1,* and two successive levels in the CM and LD muscles, where *Fat1^LacZ/LacZ^* embryos exhibit a thinner CM with overall less *Gdnf* signal. (B) On sections corresponding the the anterior part of the CM muscle, expression of markers of muscle differentiations *(MyoD, Six1,* and *Lbx1)* also shows that *Fat1^LacZ/LacZ^* embryos exhibit a selective loss of staining in the CM and not other neighboring muscles masses.

**S3 Fig:**
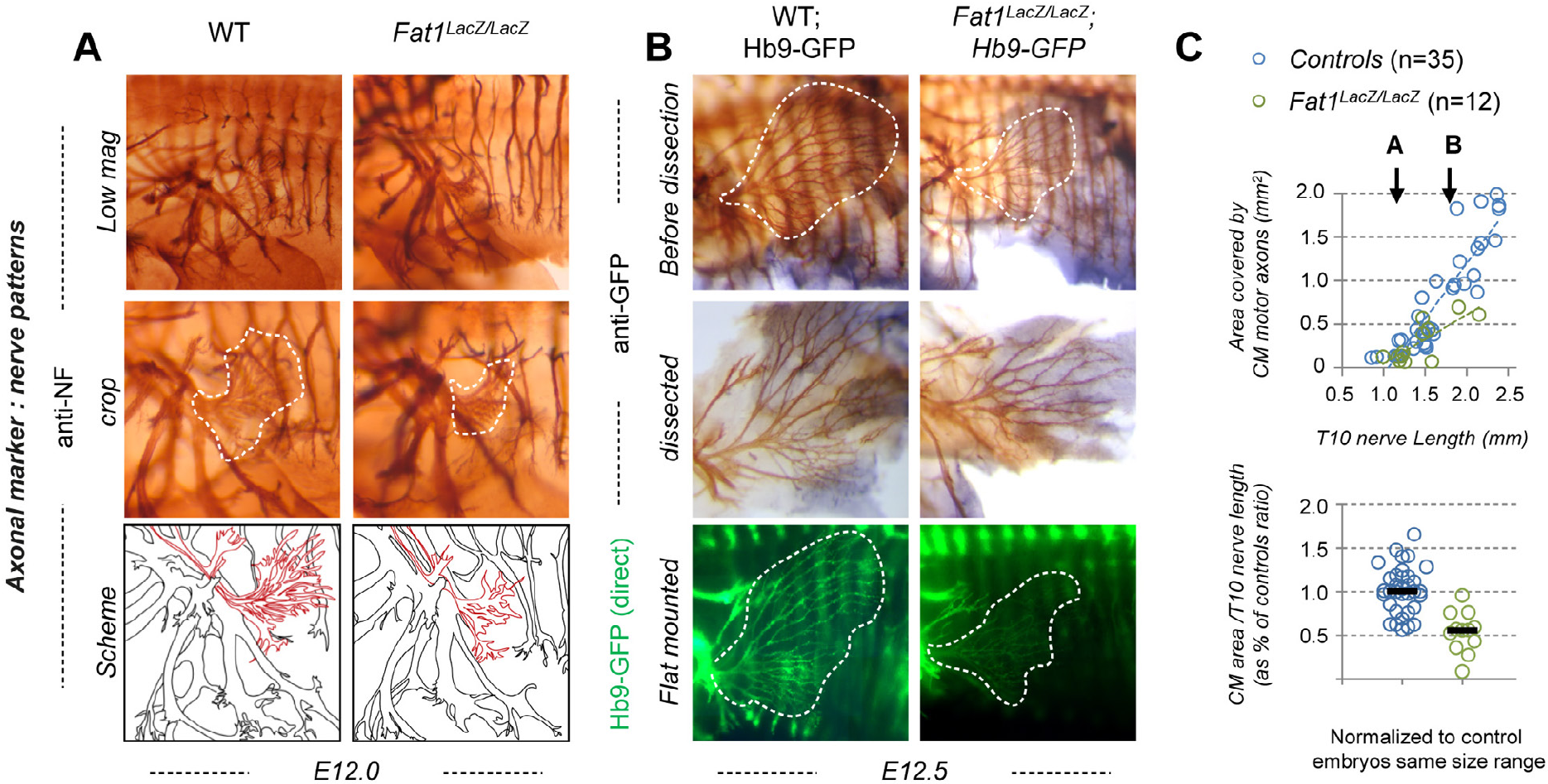
Loss of Fat1 alters motor innervation of the Cutaneous Maximus muscle. **(A, B)** The nerve pattern was analyzed by immunohistochemistry with antibodies against Neurofilament (2H3) **(A)** or by taking advantage of the Hb9-GFP transgene **(B)**, which labels motor neurons and their axons. (A) anti-neurofilament histochemistry on whole-mount wild type and *Fatl^-/-^* embryos at E12.0. **(B)** Hb9-GFP was visualized with antibodies against GFP (top and middle images), or by direct fluorescence imaging, in *Fatl^+/+^; Hb9-GFP^+^* and *Fatl^LacZ/LacZ^; Hb9-GFP^+^* embryos at E12.5. For immunohistochemistry, embryos were cut in half, cleared in BB-BA, and flat-mounted. Upper panels **(A, B)** are low magnification images of the embryo flank. Lower panels show high magnification views of the area containing the CM muscle. The area covered by CM innervated axons is highlighted in white (middle panels). In the middle panels in **(B)**, axons of vertically oriented thoracic spinal nerves have been manually removed by dissection to improve visibility of CM axons. In lower panels in **(B)** direct GFP imaging was done on PFA fixed embryos, after flat-mounting of the flank. **(C)** Quantifications of the relative expansion of the area covered by CM-innervating axons. Upper plot: for each embryo side, the area covered by CM-innervating axons is plotted relative to the length of a thoracic nerve (T10, from dorsal root origin to ventral tip). Arrows point the stages of representative examples shown in **(A)** and **(B)**. Bottom plot: for each embryo, the CM-innervated area/T10 Length was normalized to the median ratio of control embryos, by size range. Blue dots: *Fat1^+/+^* (n=35, same sample set as in controls of Fig 2); Red dots: *Fat1^LacZ/LacZ^* (n=12). Underlying data are provided in S1 Data.

**S4 Fig:**
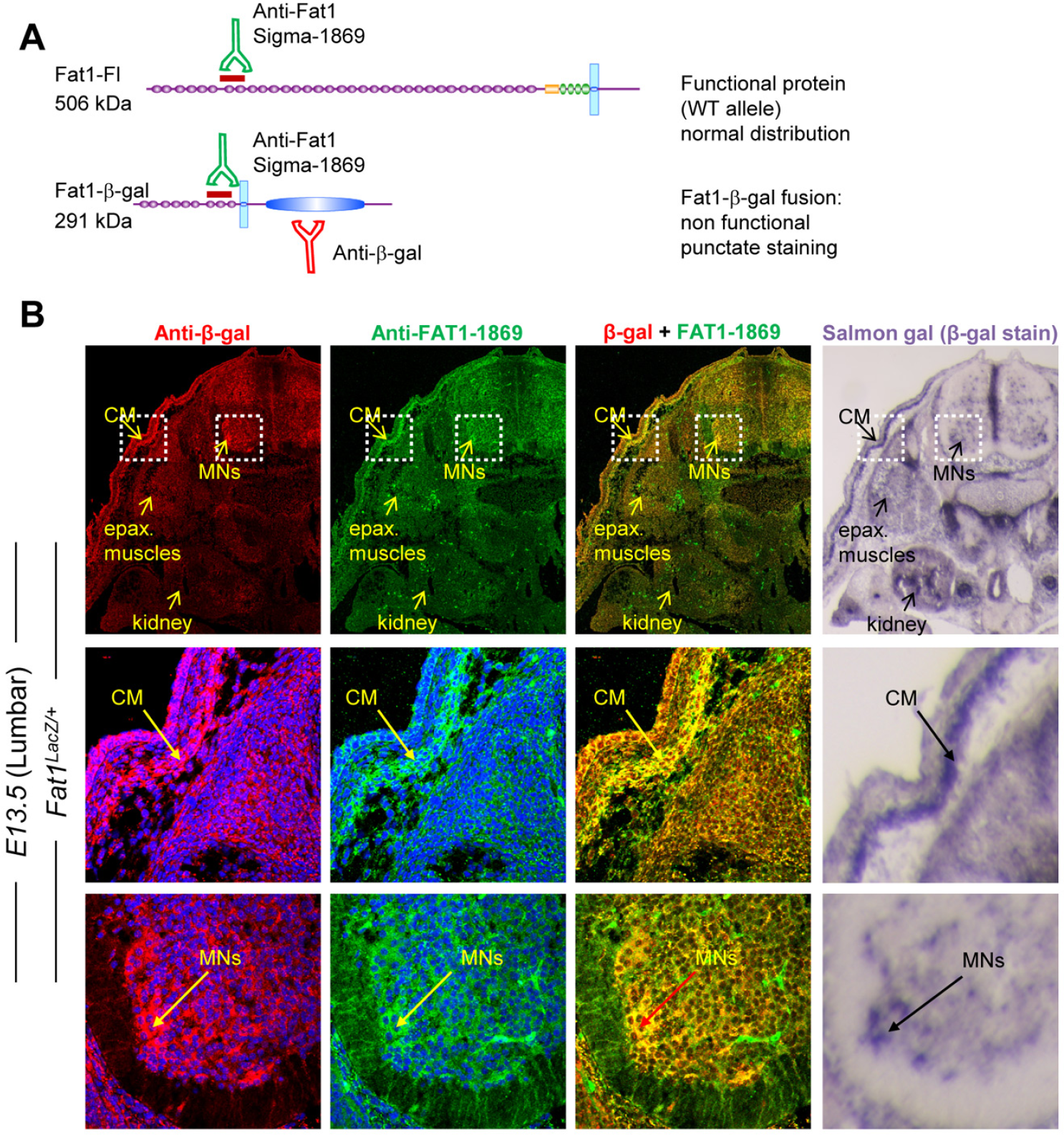
Validation of Fat1 immunohistochemistry with antibodies against the Fat1-LacZ fusion. **(A)** Scheme of the protein products of a wild type *Fat1* allele (full length Fat1) and of a *Fat1^LacZ^* allele (Chimaeric protein with the first 8 cadherin domains of Fat1 extracellular domain, fused to an exogenous transmembrane domain in frame with β-galactosidase as intracellular domain. An antibody to Fat1 (Sigma 1869) directed against a portion of the common segment of Fat1 extracellular domain, recognizes both proteins, whereas an antibody to β-galactosidase recognizes only the Fat1-β-gal fusion protein, most of which being sequestered in the golgi apparatus and not localized at the cell membrane. **(B)** Comparison of immunohistochemical detection of Fat1 in a *Fatl^LacZ^* embryo using the anti-β-galactosidase antibody (red), the Fat1-1869 antibody (green), and the pattern of β-galactosidase activity revealed by Salmon-Gal staining on cross-sections of an E13.5 mouse embryos at Lumbar levels, where it is possible to detect both the expression in subsets of MNs, and in the caudal most part of the mesenchymal subcutaneous layer towards which the CM extends. The Fat1-β-gal fusion protein is mainly detected by both antibodies as punctae in the golgi, most likely because the protein is misfolded and does not reach the cell membrane. The two stainings perfectly overlap (except for some weak staining of blood vessels in the green channel, mostly due to the secondary antibody), and correspond to the signal detected by Salmon-Gal staining. **(C)** Comparison between expression of *Fatl^LacZ^* (visualized with an anti-β-galactosidase antibody (red)) and that of Pax7 (green, muscle progenitors) on cross-sections of forelimbs of E13.5 mouse embryos. Whereas β-gal expression is detected within the Pax7-positive CM progenitors, many Pax7-negative cells within or around the muscle also express significant amount of β-gal.

**S5 Fig:**
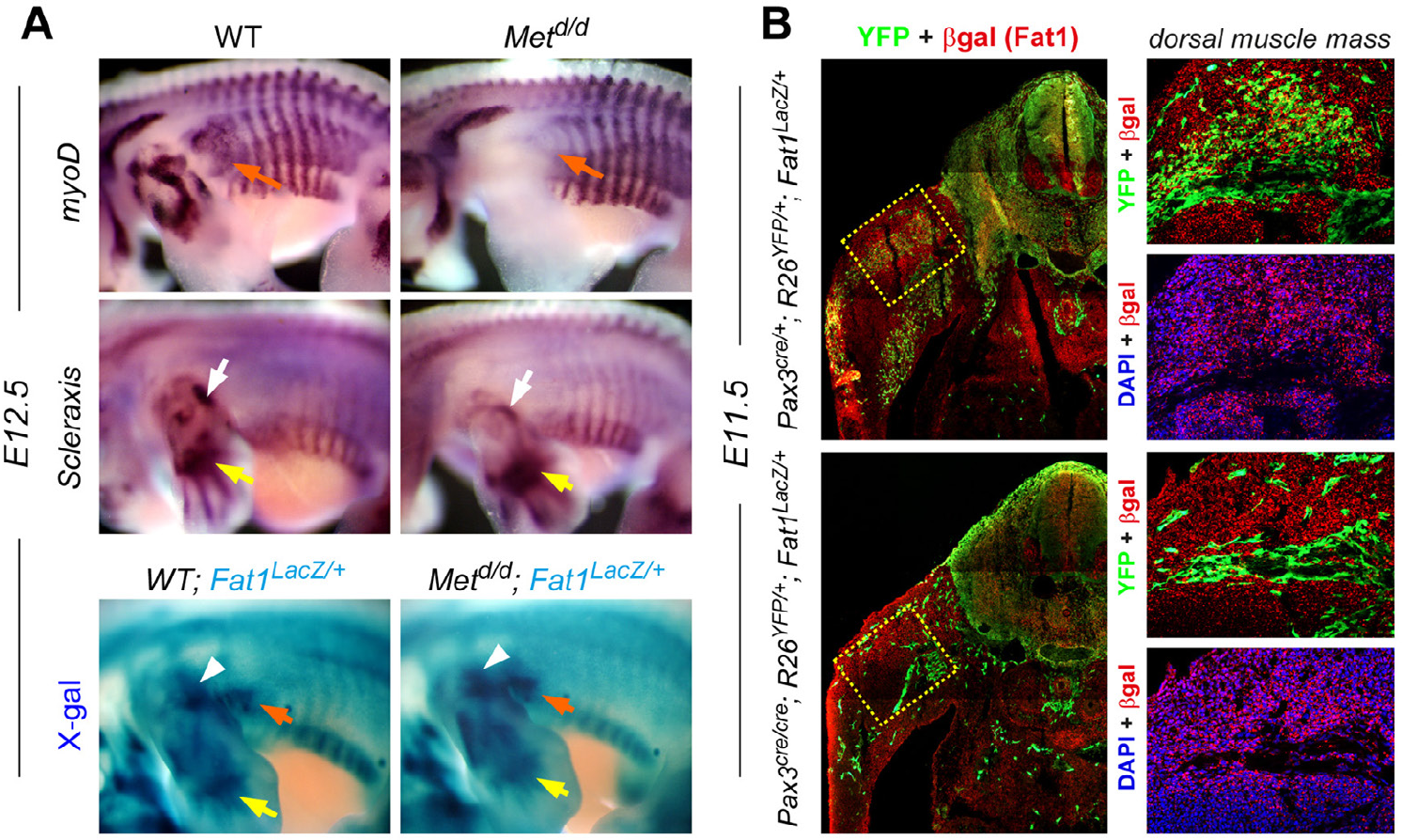
Mesenchymal *Fat1* expression persists in embryos devoid of migratory muscles. **(A)** Analysis of *MyoD* and *Scleraxis* expression in *Fatl^LacZ/+^* and *Met^d/d^* E12.5 embryos, and of *Fatl^LacZ^* expression in *Fatl^LacZ/+^* and *Met^d/d^*; *Fatl^LacZ/+^* E12.5 embryos was performed by whole-mount in situ hybridization and X-gal staining, respectively. Peripheral sites of *Fatl^LacZ^* and *Scleraxis* expression are maintained in spite of the absence of migratory appendicular muscles, indicating that expression of both genes occurs in non-myogenic/mesenchymal cells. Orange arrowheads indicate the position of the CM muscle; white arrowheads indicate the mesenchyme hotspot in the humeral region of the forelimb, yellow arrowheads indicate positions of Scleraxis-positive tendons in the distal limb (autopod). **(B)** Comparison of β-gal (red) and YFP (green) immunohistochemistry in *Pax3^cre/+^; R26^YFP/+^; Fatl^LacZ/+^* **(top)** and *Pax3^cre/cre^; R26^YFP/+^; Fatl^LacZ/+^* **(bottom)** E11.5 embryos on transverse sections at forelimb levels. The region indicated with a dotted area is shown in higher magnification on the right panels.

**S6 Fig:**
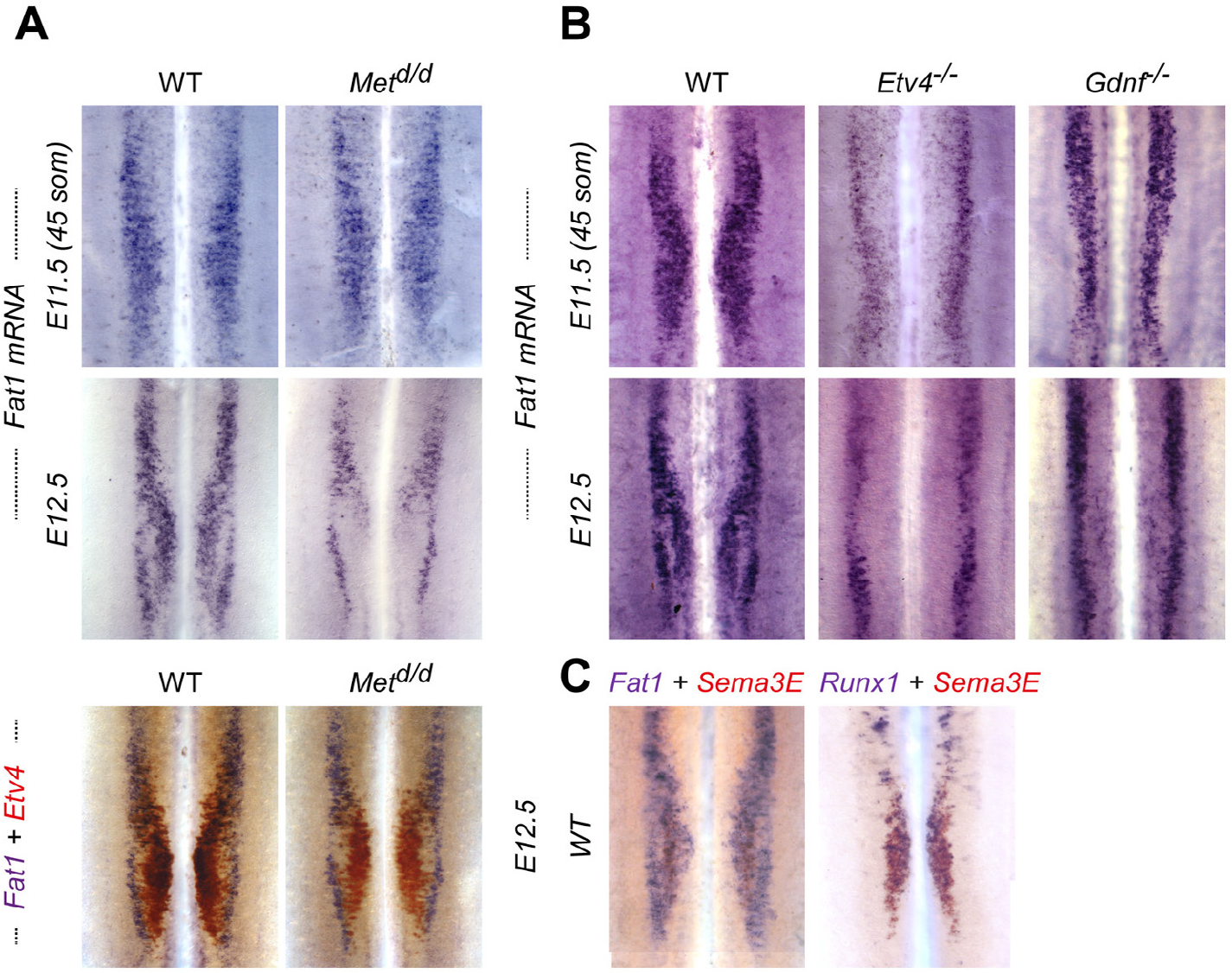
**(A)** Double in situ hybridization analysis of *Fat1* (purple) is combined with *Etv4* (red) expression in WT and *Met^d/d^* E12.5 spinal cords in lower panels. **(B)** In situ hybridization analysis of *Fat1* in WT, *Etv4^-/-^* and *Gdnf^-/-^* spinal cords at E11.5 (upper panels) and E12.5 (lower panels). **(A, B)** Onset of *Fat1* expression occurs independently of *Met* **(A)**, of *Etv4* and of *Gdnf* **(B)** functions and is preserved in *Met^d/d^*, *Etv4^-/-^* and *Gdnf^-/-^* embryos at E11.5. In contrast, analysis of *Fat1* expression at E12.5 reveals alterations consistent with 1) defective maintenance of *Fat1* expression in the CM pool in *Met^d/d^* spinal cords **(A)**, possibly due to the lack of migrating muscles and subsequent growth factor depletion, and 2) aberrant positioning of the CM pool in *Pea3^-/-^* and *Gdnf^-/-^* spinal cords **(B)**. **(C)** Comparison of *Fat1, Sema3E* and *Runx1* expression in the brachial spinal cord at E12.5 was performed by double in situ hybridization (ISH) with *Fat1* (purple) and *Sema3E* (red) on the left, and with *Runx1* (purple) and *Sema3E* (red) on the right. **(D)** *Fat1* expression in the mouse thoracic spinal cord is shown X-gal staining on sections of *Fatl^LacZ/+^* spinal cords at E12.5, showing expression in all neural progenitors in the ventricular zone, and lack of high level expression in thoracic motor columns.

**S7 Fig:**
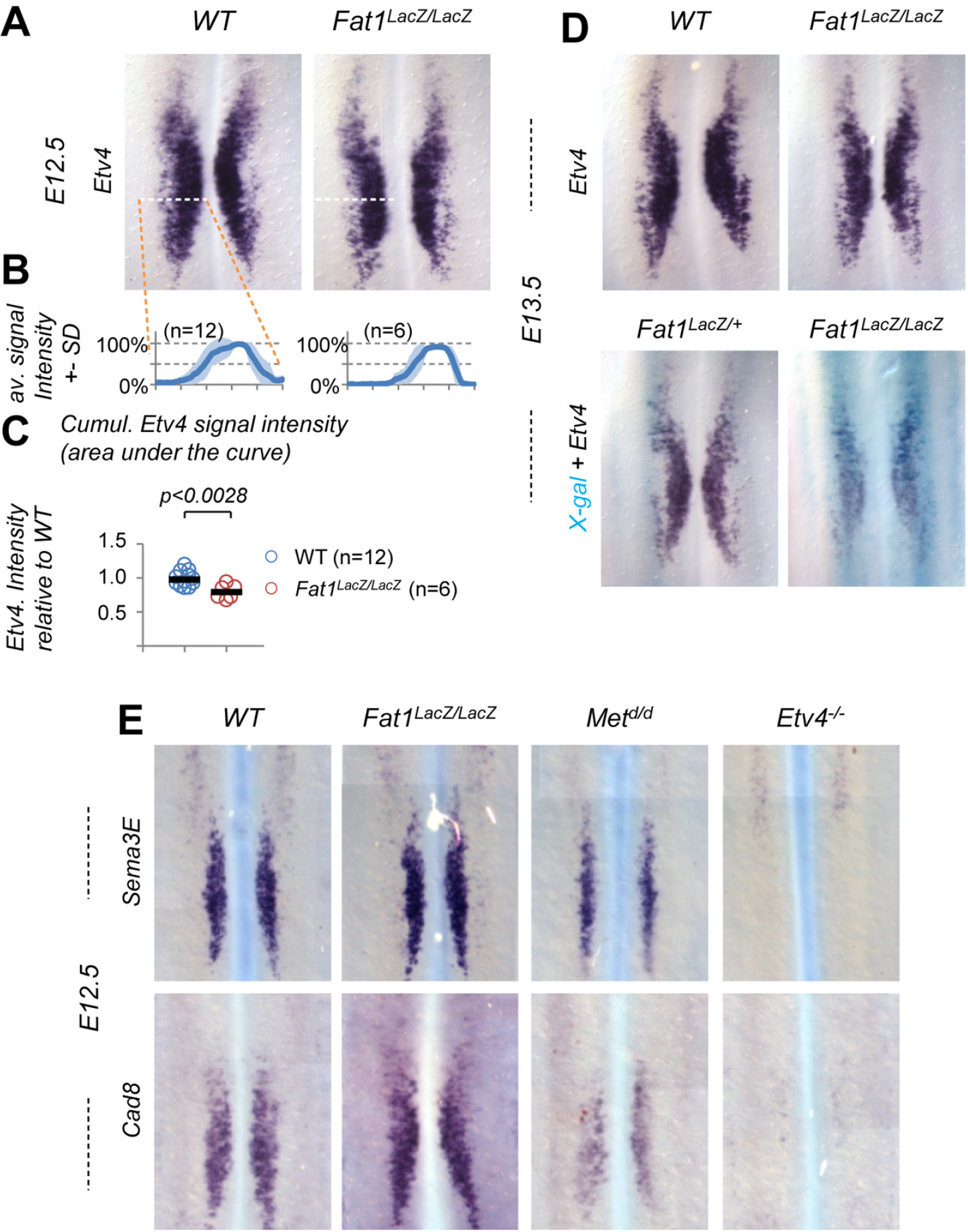
**(A-C) (A)** *Etv4* expression was analyzed by in situ hybridization in E12.5 wild type, and *Fatl^LacZ/LacZ^* spinal cords. The images represent flat-mounted spinal cords in the brachial region. **(B)** Quantifications of *Etv4* signal: each plot represents the average signal distribution (± standard deviation in light blue) measured on the indicated number of spinal cord sides along the orange dotted line in each image in **(A)** *(Fatl^+/+^* (n=12; this set of controls is the same as that shown in Fig 6); *Fatl^LacZ/LacZ^* (n=6). **(C)** Quantifications and statistical analyses of the sum of signal intensity corresponding to the area under the curves in plots shown in **(C)**: each dot represents the sum of *Etv4* intensity for each spinal cord side, the number of samples being indicated (the two sides of each embryo considered independent). (B-C): Underlying data are provided in S1 Data. **(D)** Analysis of *Etv4* expression was carried out by ISH with *Etv4* probe alone (top) on WT and *Fatl^LacZ/LacZ^* E13.5 spinal cords, or with *Etv4* probe combined with prior X-gal staining (bottom) on *Fat1^LacZ/+^* and *Fatl^LacZ/LacZ^* E13.5 spinal cords. **(E)** Analysis by ISH of *Sema3E* and *Cadherin8* expression in flat-mounted brachial spinal cords from wild type, *Fatl^LacZ/LacZ^, Met^d/d^* and *Etv4^-/-^* embryos at E12.5: Expression of *Sema3E* and *Cadherin8* in the C7-C8 segments is lost in *Etv4^-/-^* and reduced in *Met^d/d^* mutants, whereas they are unaffected in *Fatl^LacZ/LacZ^* spinal cords.

**S8 Fig:**
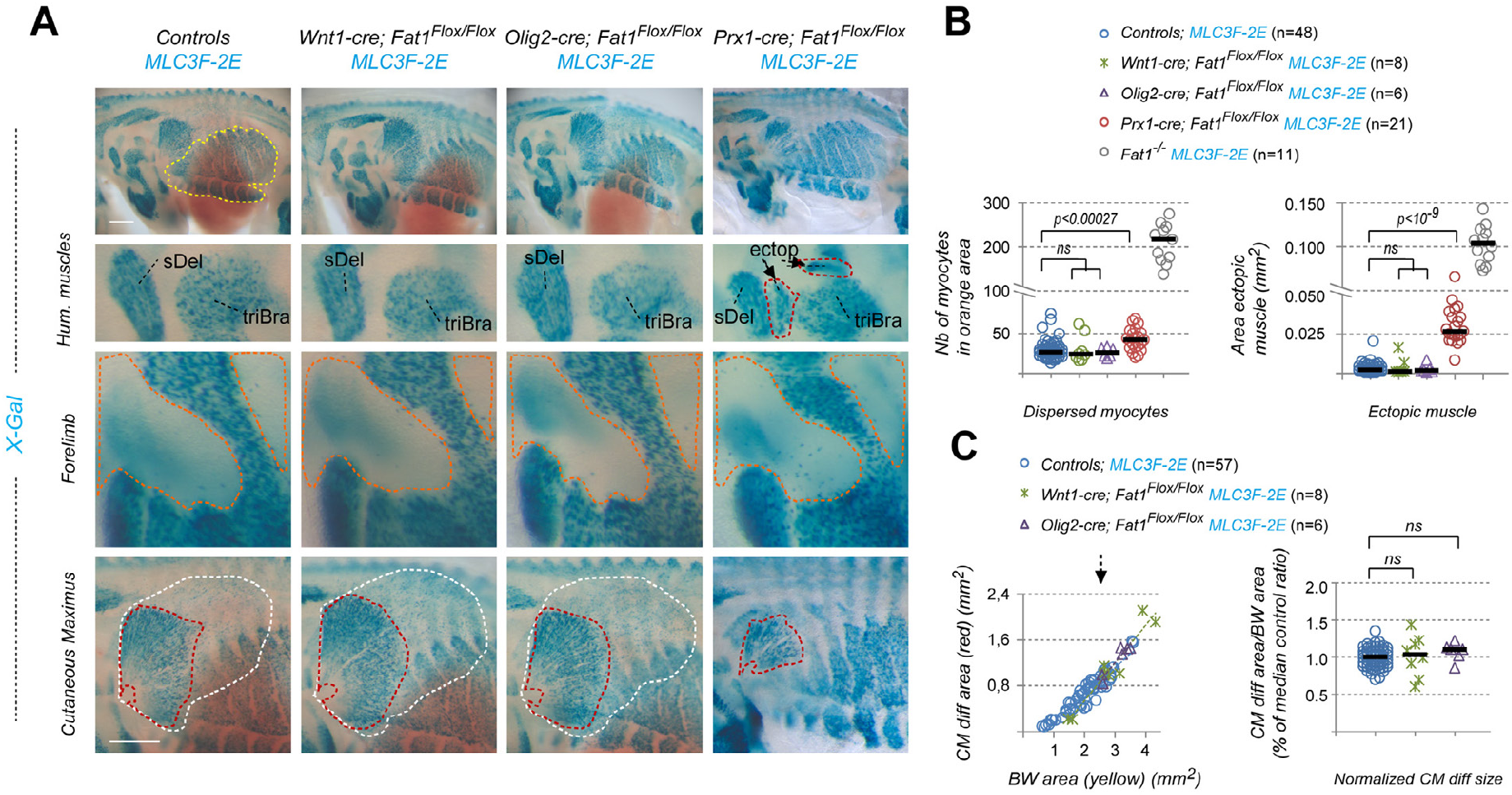
Effect of tissue-specific *Fat1* ablation on myogenic differentiation at E12.5. **(A)** Embryonic musculature was visualized by X-gal staining on E12.5 embryos carrying the *MLC3F-2E-LacZ* transgene in different contexts of tissue-specific deletion of *Fatl,* with *Wntl-cre* (neural crest cells), *Olig2-cre* (MN precursors), and *Prx1-cre* (limb/lateral mesoderm-derived mesenchyme). Top images are images of the entire flank of *Fatl^Flox/Flox^; MLC3F-2E* controls, and *Wntl-cre; Fatl^Flox/Flox^; MLC3F-2E*, *Olig-cre; Fatl^Flox/Flox^*; *MLC3F-2E,* and *Prx1-cre; Fatl^Flox/Flox^*; *MLC3F-2E* embryos, respectively. Higher magnification pictures focus (from top to bottom) on humeral muscles, on the forelimb area, and on the flank region in which the CM spreads, respectively. **(B)** Quantification of the number of dispersed myocytes found in orange areas in the forelimb (Left plot), and of the area occupied by the ectopic humeral muscle (ectop) appearing between the spinodeltoid (Del) and the triceps brachii (TriBra) muscles (right plot). In both graphs, data from *Fatl^-/-^* embryos (from plots in S1B Fig) have been added in the last lane, for comparison of effect size. Note that on both graphs there is an interruption on the Y axis, and a change of scale. Only *Prx1-cre; Fatl^Flox/Flox^*; *MLC3F-2E* embryos exhibit a mild increase compared to controls. Underlying data are provided in S1 Data. **(C)** Quantification of the expansion rate of differentiated CM fibers in *Fatl^Flox/Flox^*; *MLC3F-2E* controls, *Wntl-cre; Fatl^Flox/Flox^; MLC3F-2E,* and *Olig-cre; Fatl^Flox/Flox^; MLC3F-2E.* Quantifications of *Prxl-cre; Fatl^Flox/Flox^; MLC3F-2E* embryos are shown in Fig 7B. Left graph: For each embryo side, the area covered by differentiated CM fibers was plotted relative to the area occupied by body wall muscles. Right plot: for each embryo, the CM area/Body wall area was normalized to the median ratio of control embryos. Blue dots: *Fat1^Flox/Flox^; MLC3F-2E* (n=57, includes the same set of controls as Fig 7B); Green crosses: *Wnt1-cre; Fatl^Flox/Flox^; MLC3F-2E* (n=8); Purple triangles: *Olig2-cre; Fatl^Flox/Flox^; MLC3F-2E* (n=6). Underlying data are provided in S1 Data. Scale bars **(A)** 500 μm.

**S9 Fig:**
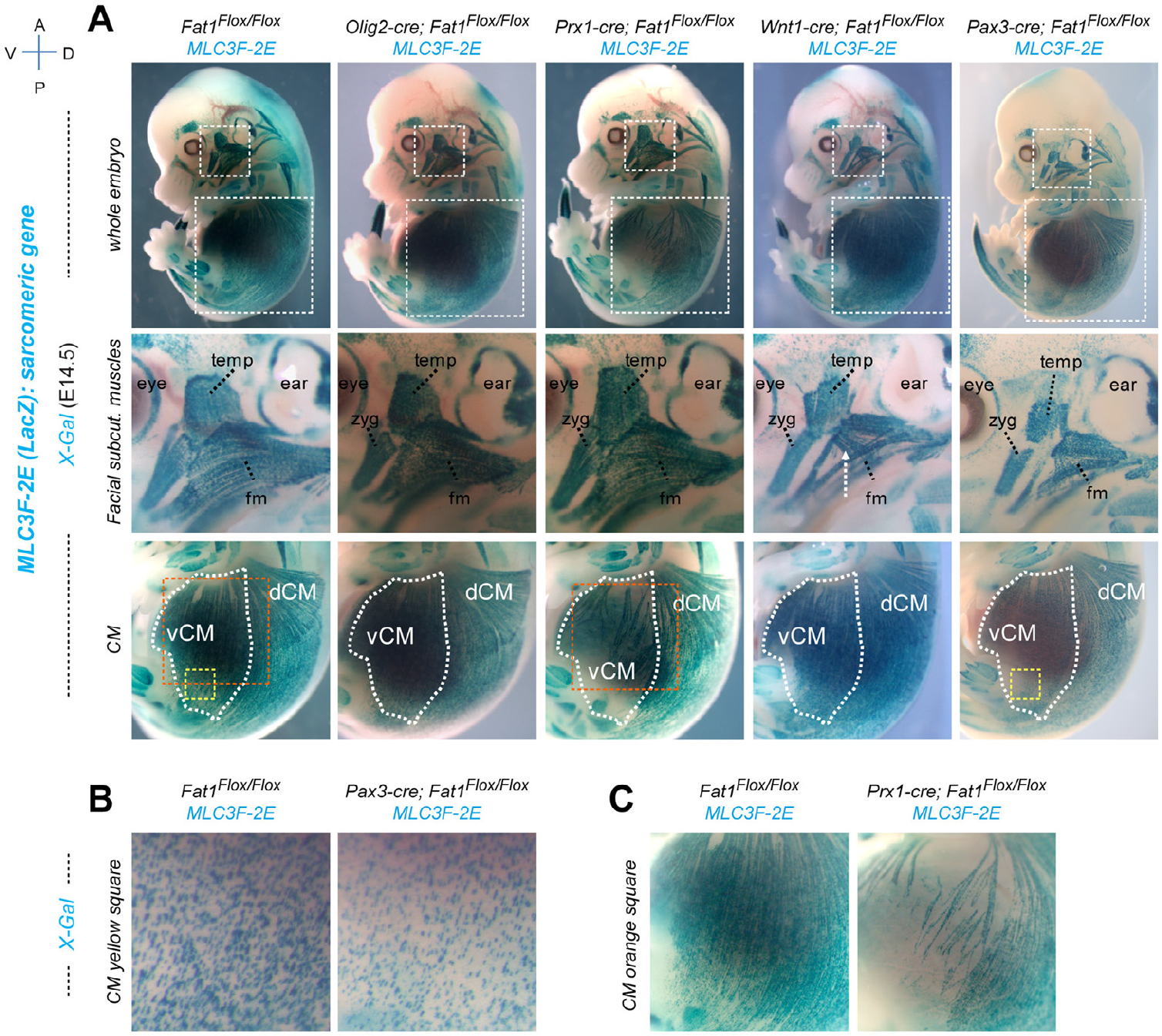
Effect of tissue-specific *Fat1* ablation on fetal musculature at E14.5). **(A)** The respective effects of depletion with *Wnt1-cre, Olig2-cre, Prx1-cre,* and Pax3-cre (premigratory myogenic cells + neural crest lineage) were assessed by whole mount X-gal staining on E14.5 *Fat1^Flox/Flox^; MLC3F-2E* controls (1^st^ column), *Olig-cre; Fat1^Flox/Flox^; MLC3F-2E* (2^nd^ column), *Prx1-cre; Fat1^Flox/Flox^; MLC3F-2E* (3^rd^ column), *Wnt1-cre; Fat1^Flox/Flox^; MLC3F-2E* (4^th^ column), and *Pax3^cre/+^; Fat1^Flox/Flox^; MLC3F-2E* (5^th^ column) embryos. Top panels: Low magnification pictures show whole-embryo side-views. Boxed areas show the position of higher magnification pictures shown in lower panels. Middle panels focus respectively on facial subcutaneous muscles. Lower panels focus on the cutaneous Maximus muscle. *Prx1-cre; Fat1^Flox/Flox^* embryos display normal musculature in the face, but abnormal muscle shape and intramuscular orientation in the CM (higher magnification of the orange box shown in **(C)**), while *Wnt1-cre; Fatl^Flox/Flox^* embryos have a normal CM, but severely affected subcutaneous muscles in the face, compared to controls (*Fatl^FloxFlox^*). Thus, ablation in neural crest cells causes severe alterations in shape of the facial subcutaneous muscles in both *Pax3^cre/+^; Fatl^Flox/Flox^* and *Wnt1-cre; Fatl^Flox/Flox^* embryos. In contrasts, muscle shapes of *Olig2-cre; Fatl^Flox/Flox^* embryos are indistinguishable from those seen in controls. **(B, C)** Higher magnification views of fiber density in the CM in yellow and orange boxed areas outlined in lower panels in **(A)**, respectively comparing control with *Pax3^cre/+^; Fatl^Flox/Flox^* **(B)**, and with *Prxl-cre; Fatl^Flox/Flox^* **(C)** embryos. The effect of *Fat1* ablation in *Pax3^cre/+^; Fatl^Flox/Flox^* embryos on CM appearance **(B)** is modest at E14.5 (high magnification images illustrate a subtle reduction in the density of *LacZ*-positive nuclei). In contrast, the reduction in fiber density observed in *Prxl-cre; Fatl^Flox/Flox^* embryos **(C)** is very severe in the ventral CM, in contrast to more dorsal regions, coinciding with the limit above which *Prx1-cre* is not active.

**S10 Fig:**
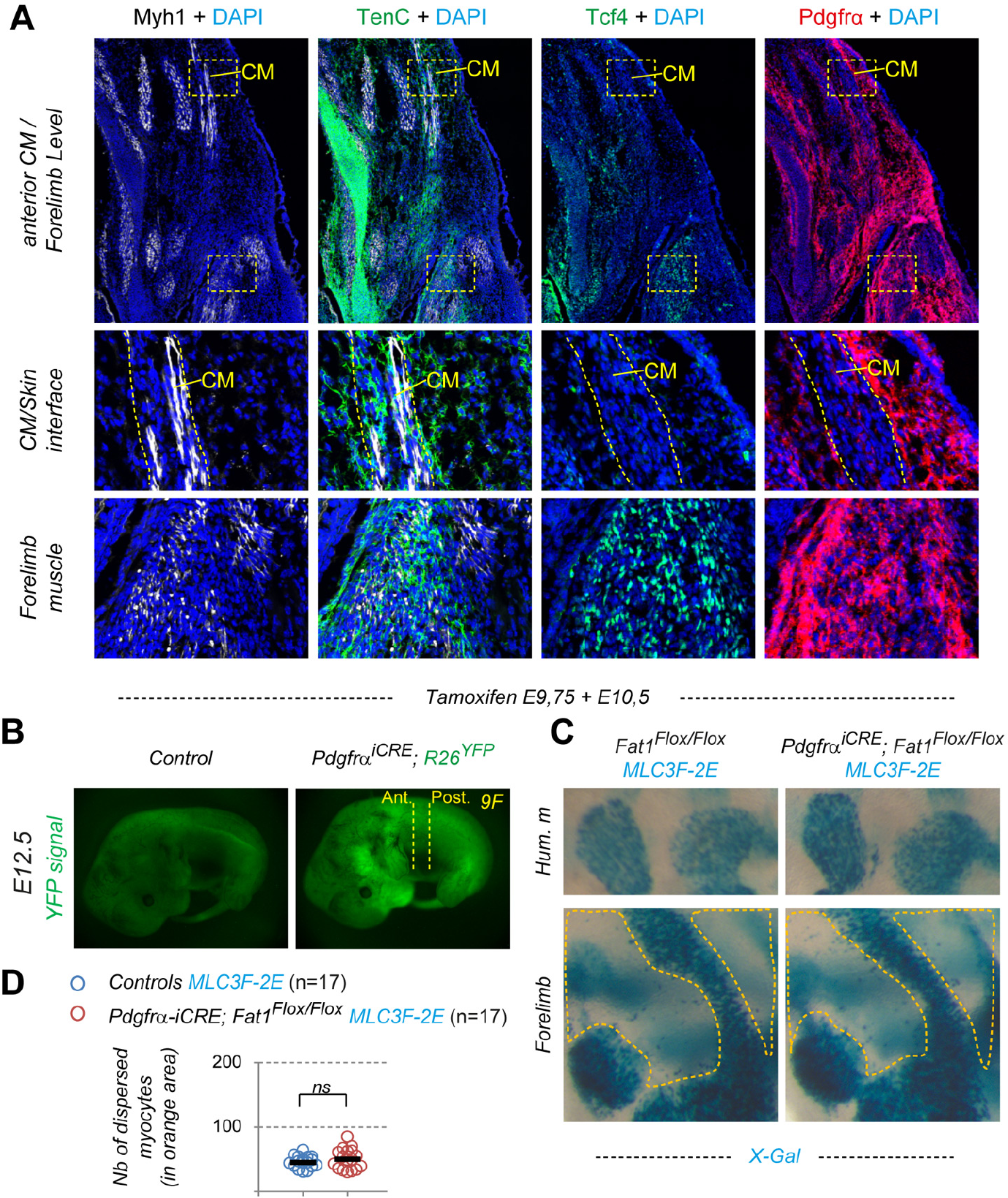
The CM surrounding connective tissue belongs to the Pdgfrα lineage. **(A)** Characterization of the mesenchyme subtype at the CM-skin interface. Sections of an E12.5 wild type embryo at the level of the anterior CM (and upper forelimb) were immunostained with antibodies against Myh1 (white), TenascinC (green), Tcf4 (green), Pdgfrα (red), and DAPI (blue). The lower panels represent higher magnifications of the two bowed areas indicated in the low magnification views, respectively focusing on the mesenchyme surrounding the CM and at the interface between CM and skin (middle panels) and the extremity of one forelimb muscle (lower panel). Whereas Tcf4 is detected in myogenic cells and connective tissue fibroblasts associated with the forelimb muscle, no expression is detected in the CM/skin connective tissue, which in contrasts expresses TenascinC and Pdgfrα. **(B)** Control and *Pdgfrα-iCre; R26^YFP/+^* embryos were collected at E12.5 from a pregnant female treated with Tamoxifen at E9.5 + E10.5. Images show the direct fluorescence for acquired at the same exposure time. The vertical lines in the lower picture highlight the level of sections corresponding to Fig 9F (anterior CM and Posterior CM). **(C)** Whole-mount β-galactosidase staining was performed on Tamoxifen treated *control; MLC3F-2E* (left) and *Pdgfrα-iCre; Fatl^Flox/Flox^; MLC3F-2E* (right) embryos at E12.5. Top images represent the upper forelimb region (whith scapulohumeral muscles), bottom images represent the forelimb regio where myocyte dispersal is being quantified in **(D)**. **(D)** Quantification of the number of dispersed myoblasts observed in the forelimb (counted in the areas delimited by orange lines). Blue dots: *Controls; MLC3F-2E* (n=17, control embryos from Tamoxifen-treated litters); Red dots: *Pdgfrα-iCre; Fatl^Flox/Flox^; MLC3F-2E* (n=17). Underlying data are provided in S1 Data.

**S11 Fig:**
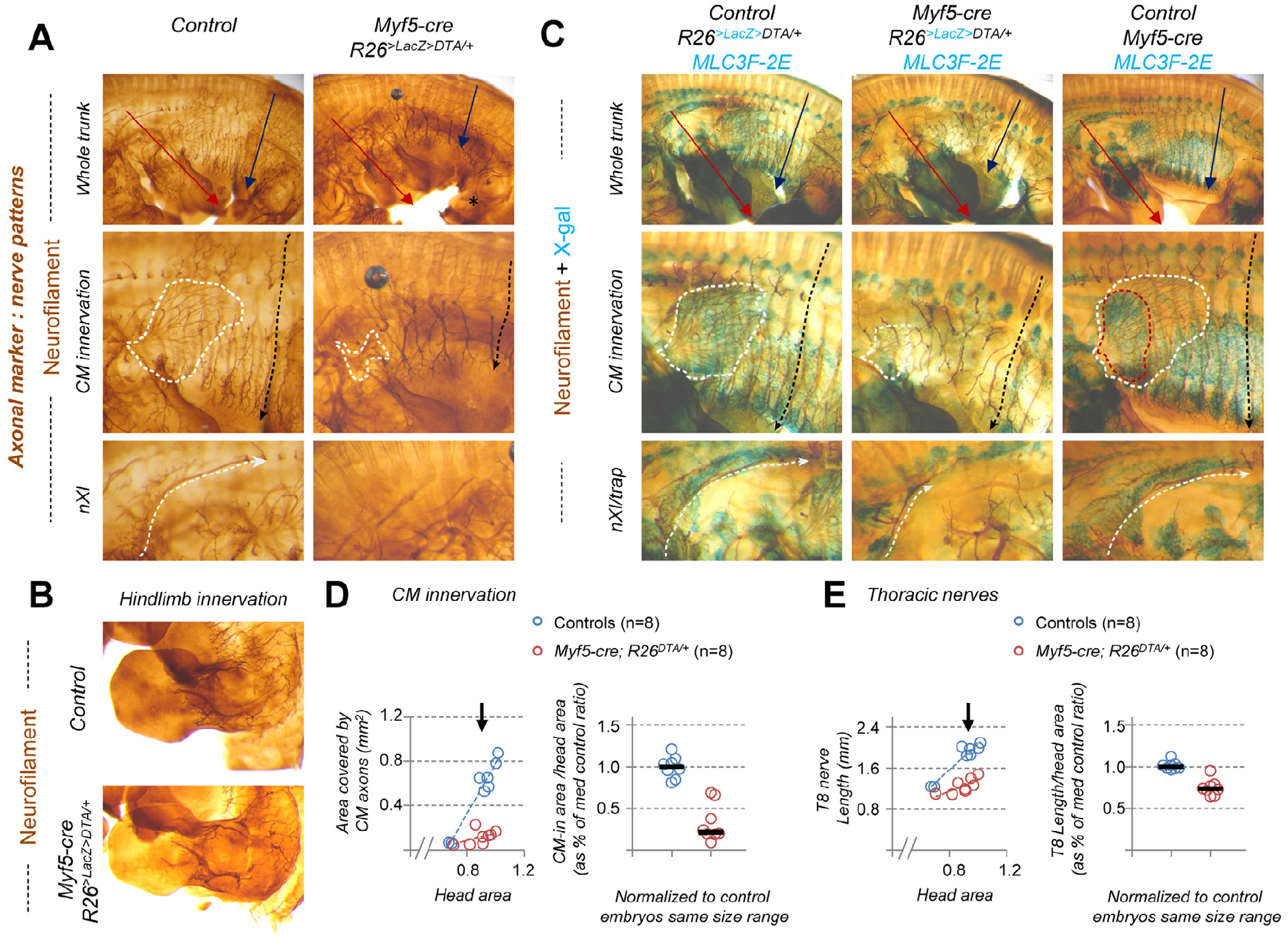
Genetic ablation of myogenic cells interferes with axonal growth of several classes of MNs comprising those innervating the CM, the trapezius muscle, and hypaxial bodywall muscles. **(A-C)** Anti-Neurofilament immunohistochemistry was performed on E12.5 embryos, in the context of *Myf5-cre*-driven myogenic lineage depletion, mediated by conditional Diphtheria Toxin A (DTA) expression under regulatory control of the *R26* locus (*R26^Lox-LacZ-STOP-Lox-DTA^* mouse line). This nerve pattern is either visualized on its own **(A, B)** or combined with X-gal staining, when embryos also carry the *MLC3F-2E-LacZ* transgene **(C)**. In cases with *MLC3F-2E* **(C)**, X-Gal staining also detects *LacZ* expression driven by the *R26* locus in un-recombined cells (the relative expression levels in the two loci *(R26* and *MLC3F-2E-LacZ* transgene) allows visualizing muscles in the trunk area but not in the limb). After BA-BB clearing, the embryos were cut in half, and the internal organs and skin removed to visualize the CM and brachial plexus. Genotypes shown are: control *R26^Lox-LacZ-STOP-Lox-DTA^;* and *Myf5-cre; R26^Lox-LacZ-STOP-Lox-DTA^* embryos **(A, B)**; control *R26^Lox-LacZ-STOP-Lox-DTA^; MLC3F-2E* (left), *Myf5-cre; R26^Lox-LacZ-STOP-Lox-DTA^; MLC3F-2E* (middle), or *Myf5-cre; R26^Lox-LacZ-STOP-Lox-DTA^; MLC3F-2E* (right) **(C)**. Upper panels show the entire flank of embryos at comparable stages. The area covered by CM-innervating axons is outlined with white-dotted lines. The extent of the T8 thoracic nerve is outlined with a black arrow. The length of the forelimb (slightly affected by muscle depletion) is highlighted by a red arrow. Middle panels show higher magnification of the upper thoracic region containing the CM and corresponding motor axons as well as thoracic nerves, after manual removal by dissection of most cutaneous sensory nerves. Lower panels show the upper cervical region, with the trapezius muscle, innervated by the cranial nerve XI, also called spinal accessory nerve. **(D, E)** Quantifications: of the progression of CM innervation **(D)** and of thoracic nerve growth **(E)**. Left plots: For each embryo side, the area covered by CM-innervating axons **(D)**, or the length of the T8 nerve **(E)** were plotted relative to the head area, comparing *Myf5-cre; R26^Lox-LacZ-STOP-Lox-DTA^;* embryos (red dots) to controls (blue dots). The T8 value represents the distance between the dorsal tip of sensory nerve roots and the ventral-most axon tips on the flank, thus including both sensory and motor components. Head area was used as reference as it grows proportionally to embryo growth, and it is not affected by muscle depletion, in contrast to other distances/areas in the trunk or limbs. Right plots: for each embryo side, we calculated the ratios CM-innervated area/head area **(D)** and T8 Length/head area **(E)**, and these values were normalized to the median ratio of control embryos, by size range. Blue dots: Controls (n=8, pooling all control littermates); Red dots: *Myf5-cre; R26^Lox-LacZ-STOP-Lox-DTA^* (n=8). **(D, E)**: Underlying data are provided in S1 Data. Arrows point to the stage of representative examples shown in **(A-C)**.

**S12 Fig:**
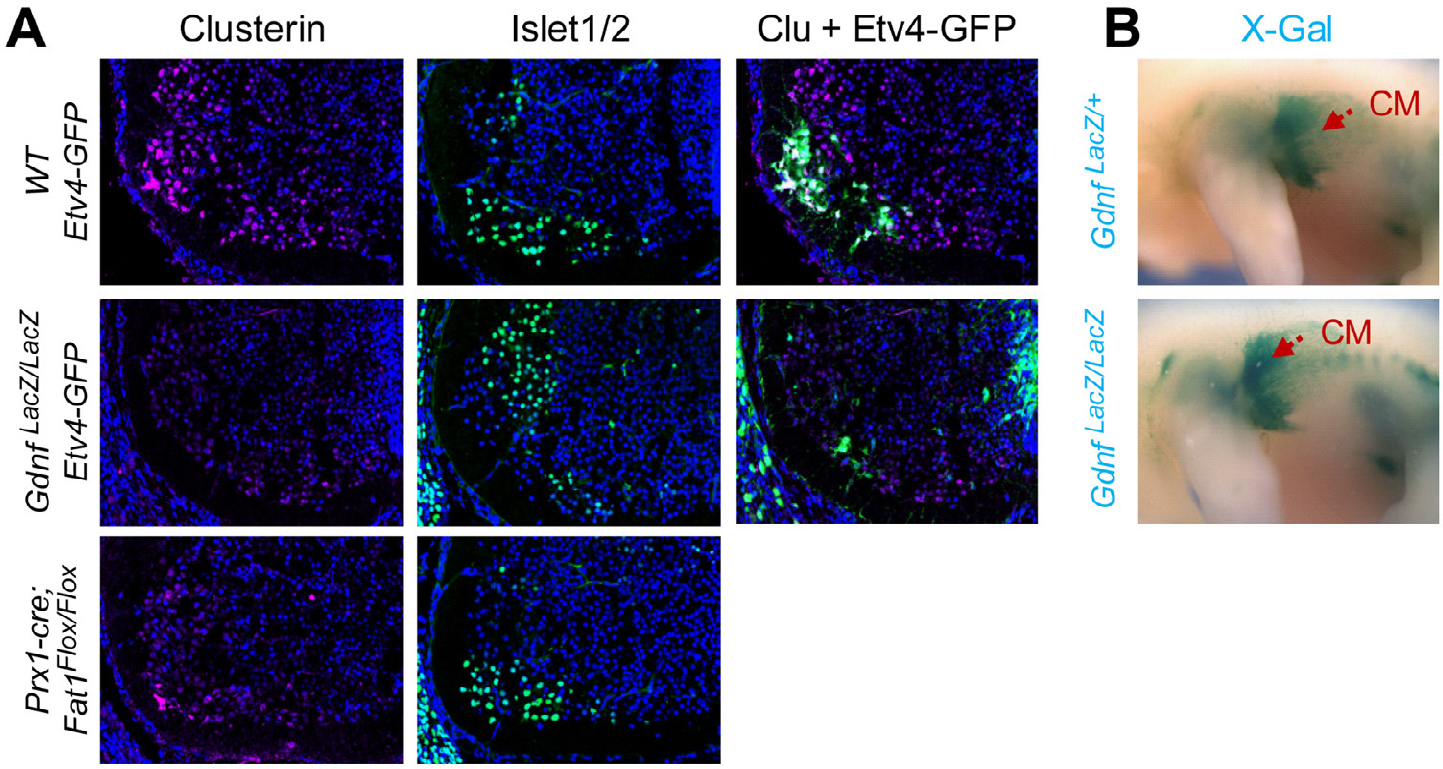
Complementary analyses of MN and CM muscle phenotypes in *Gdnf* mutants. **(A)** Analysis by immunohistochemistry of Clusterin, islet1, and GFP expression on spinal cord sections at C7-C8 level from E13.5 *WT; Etv4-GFP/+,* from *Gdnf^LacZ/LacZ^; Etv4-GFP/+,* and from *Prxl-cre; Fatl^Flox/Flox^* embryos. Clusterin and GFP staining were done one sections neighboring those analyzed with Islet1/2 IHC. *Etv4-GFP* could not be combined with *Prx1-cre,* as this combination appears to cause a phenotype (short limbs, with impaired elongation, visible from E12.5 onward) unrelated to *Fat1* alleles, and most-likely linked to the site of integration of both transgenes. **(B)** X-gal staining of *Gdnf^LacZ/+^* and *Gdnf^LacZ/LacZ^* embryos, showing that expansion of the area covered by *Gdnf^LacZ^*-expressing CM progenitors is not affected by the lack of *Gdnf.*

